# Tuning Encodable Tetrazine Chemistry for Site-Specific Protein Bioorthogonal Ligations

**DOI:** 10.1101/2025.05.20.655094

**Authors:** Subhashis Jana, Alex J. Eddins, Yogesh M. Gangarde, P. Andrew Karplus, Ryan A. Mehl

## Abstract

Using genetic code expansion (GCE) to encode bioorthogonal chemistry has emerged as a promising method for protein labeling, both in vitro and within cells. Here, we demonstrate that tetrazine amino acids incorporated into proteins are highly tunable and have extraordinary potential for fast and quantitative bioorthogonal ligations. We describe the synthesis and characterize reaction rates of 29 tetrazine amino acids (20 of which are new) and compare their encoding ability into proteins using evolved Tet ncAA encoding tRNA/RS pairs. For these systems, we characterized on-protein Tet stability, reaction rates, and ligation extents, as the utility of a bioorthogonal labeling group depends on its stability and reactivity when encoded into proteins. By integrating data on encoding efficiency, selectivity, on-protein stability, and in-cell labeling for Tet tRNA/RS pairs, we developed the smallest, fastest, and most stable Tet system to date. This was achieved by introducing fluorine substituents to Tet4, resulting in reaction rates at the 10⁶ M⁻¹s⁻¹ level while minimizing degradation. This study expands the toolbox of bioorthogonal reagents for Tet-sTCO-based, site-specific protein labeling and demonstrates that the Tet-ncAA is a uniquely tunable, highly reactive, and encodable bioorthogonal functional group. These findings provide a foundation to further explore Tet-ncAA encoding and reactivity.

## Introduction

The encoding of bioorthogonal chemistry into proteins has been critical for engineering protein materials, developing new protein therapeutics, and advancing studies on protein function *in vitro* and in cells.^[1,2]^ The most powerful method for the general encoding of these chemistries into proteins is genetic code expansion (GCE), an approach in which a reactive functional group is included in the side chain of a noncanonical amino acid (ncAA) that is installed into a protein during translation by using a tRNA/amino acyl-tRNA synthetase (tRNA/RS) pair engineered to be specific for the ncAA.^[3,4]^ While many bioorthogonal ligation reactions have been encoded using GCE, the ligation chemistries that launched the field – oxime, Staudinger, photo-click, strain-promoted and copper catalyzed azide-alkyne cycloaddition ligations ^[5–7]^ – have relatively slow reaction rates (10^-3^ - 50 M^-1^ s^-1^) (Figure 1) that are not fast enough to be suitable for applications involving physiologically relevant protein concentrations or intracellular protein labeling.^[8,9]^

**Figure 1.**
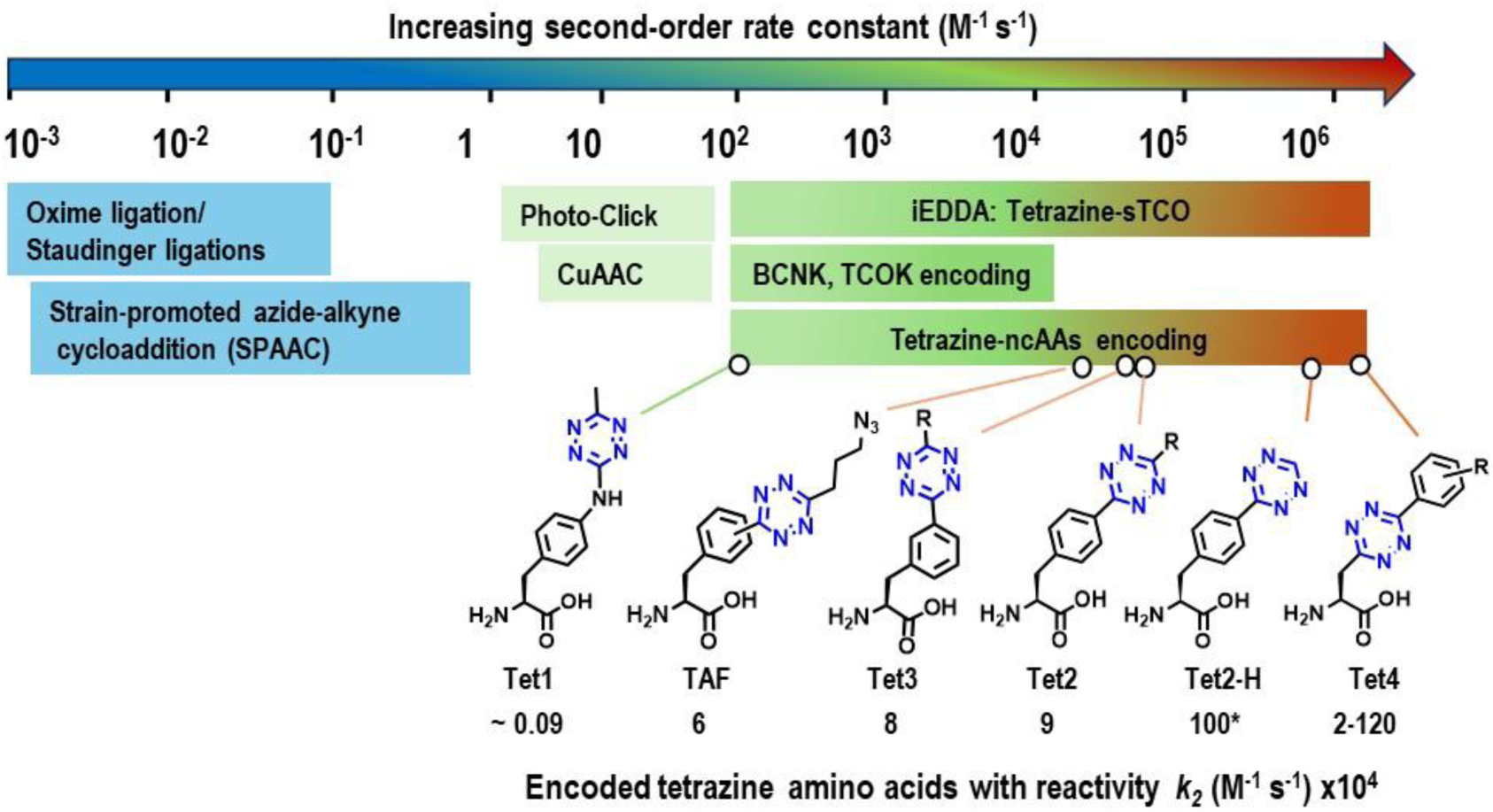
Rate constants for common bioorthogonal ligations. Bioorthogonal ligation chemistries are organized on a color-coded log scale by their reaction rates (colors start slow as blue-to-green-to-red as fastest). The structures of tetrazine amino acids previously encoded by GCE are shown below the scale along with the second-order rate constants for reaction with sTCO. *The estimated rate constant of Tet2-H with sTCO is based on data from a reported paper.^[10]^

The subsequent introduction of the initial GCE systems encoding the faster inverse electron demand Diels-Alder (IEDDA) reaction between tetrazines (Tet) and dienophiles^[8,11]^ – specifically transcyclooctenes (TCOs) and strained transcyclooctenes (sTCOs) introduced by Fox et al.^[8,12,13]^ – provided access to much faster reaction rates. These reaction rates of 10^2^ – 10^4^ M^-1^ s^-1^ met the criteria for ideal bioorthogonal ligations^[14]^ that would be sufficient for intracellular quantitative labeling at biologically relevant protein concentrations and time-sensitive biological studies (Figure 1). Knowing that IEDDA reactions between tetrazines and TCOs could, in principle, deliver even faster reaction rates that could improve the homogeneity (i.e. completeness) of live cell protein labeling, ^[15,16]^ we and others have explored encoding ncAAs with more reactive TCO and Tet moieties. Through this work novel encodable Tet-ncAAs showed up to an ∼100-fold increase in reaction rate (to over 10^6^ M^-1^ s^-1^; Figure 1) while maintaining minimal degradation of functional groups and site-specific encoding.^[14,17,18]^ In contrast, by encoding strained dienophile ncAAs it has not been possible to exceed reaction rates of ∼10^3^ M^−1^ s^−1^ while avoiding problematic degradation. ^[19,20]^

Since the 2012 encoding of the first Tet-ncAA, designated Tet1, additional encodable and more reactive Tet ncAAs have been developed (Figure 1).^[21]^ For Tet1, the maximal on-protein reaction rate with sTCO reagents was 9 × 10^2^ M^−1^ s^−1^ and the resulting secondary amine linkage to the protein was susceptible to low levels of elimination. We next developed GCE encoding machinery for two alternate Tet-ncAA scaffolds, Tet2 and Tet3 (Figure 1), which achieved on-protein reactivities exceeding 10^4^ M^-1^ s^-1^, fast enough to be considered ideal bioorthogonal labeling reagents for use in cells. *Methanococcus jannaschii* and *Methanosarcina barkeri* tRNA/RS pairs were selected to function on the Tet2-Me and Tet3-Me derivatives, respectively, and designated Tet2-tRNA/RS_Mj_ and Tet3-tRNA/RS_Mb_, respectively.^[9,17]^

We and others realized that these tRNA/RS pairs had permissive substrate profiles, and so could also encode additional Tet-ncAA structures, expanding the scope of Tet-ncAAs for functional evaluation. Initially, we focused on improving Tet encoding efficiency by exploring larger alkyl substituents predicted to better fill the active site, and achieved a 1.5-fold increase improvement by using Tet2-ethyl with the Tet2 tRNA/RS pair^[22]^ and a 3-fold increase in using Tet3-butyl with the Tet3 tRNA/RS pair.^[9]^ The Kwon lab demonstrated that the Tet2-tRNA/RS_Mj_ pair could encode the smaller Tet2-proton ncAA (Figure 1) which reacted 10 times faster, presumably due to reduced steric clash for the IEDDA ligation. The Liu lab added a second azide bioconjugation functional group to the Tet-ncAA, encoding both pTAF and mTAF (Figure 1) using the Tet2-tRNA/RS_Mj_ and Tet3-tRNA/RS_Mb_, respectively opening access to one-pot preparation of protein multiconjugates.^[23]^

To minimize the Tet-ncAA size and bring the tetrazine ring closer to the protein backbone, we developed the Tet4 scaffold (Figure 1) and selected two Tet4-tRNA/RS_Mb_, pairs to encode the Tet4-Ph ncAA. To our surprise, we discovered that bringing the tetrazine functionality closer to the protein surface provided a ∼10-fold increase in on-protein labeling rates compared to free Tet-ncAA as opposed to the ∼3-fold rate increase as seen by the Tet2 and Tet 3 systems. Additionally, seeking a still faster-reacting Tet4-derivative, we showed that one of the Tet4-tRNA/RS_Mb_ pairs could accommodate Tet4-pyridyl encoding which pushed on-protein labeling rates to above 10^6^ M^-1^ s^-1^. These increases in bioorthogonal protein labeling rates when combined with site-specifically attaching spin-labels allowed for the first double electron−electron resonance (DEER) spectroscopy to monitor structural states and the conformational equilibria of biomacromolecules inside of cells. ^[18]^

Another attractive attribute of tetrazine dienes, as compared to the TCO strained dienophiles, is their wide chemical space for tuning reactivity and substituent size. Since the report of the bioorthogonal potential for the tetrazine reaction by Fox and co-workers in 2008, there have been numerous studies that explore the chemical range of functional groups on tetrazine, and found that by altering electron withdrawing abilities, distortion effects and structural sizes, Tet reactivity is tunable over 6 orders of magnitude.^[24–26]^ Given that encoding Tet ncAAs with the Tet2, Tet3, and Tet4 scaffolds have already led to ideal bioorthogonal labeling of proteins in live cells (i.e. with quantitative yields at low concentrations and with exquisite chemoselectivity), having more choices among properties could enhance their functionality for protein bioconjugation.^[9,17,18]^ Thus, we sought here to explore more broadly the vast chemical space of tetrazine amino acids and assess the encodability and potential utility for fast, and quantitative bioorthogonal ligations.

We chose to synthesize Tet-ncAAs by varying the substituents on the Tet2, Tet3 and Tet4 scaffolds, so the new Tet-ncAAs would have the greatest likelihood of compatibility with the currently available Tet-RS/tRNA pairs. We focused especially on the Tet3 and Tet4 scaffolds since these systems are eukaryotic cell compatible. Here we describe the synthetic accessibility and reaction rates of **29** Tet-ncAAs, **20** of which have never been tested for incorporation into proteins, and assess their encodability using the existing evolved Tet2, Tet3 and Tet4 tRNA/RS pairs. Because the utility of a bioorthogonal labeling group depends on its properties when encoded into a protein, we assessed the on-protein stability of the encoded tetrazines as well as their rate and extent of reactivity. Among the three Tet-ncAA scaffolds, Tet4 variants demonstrate the fastest reactivity for a given modification. However, the fastest variant, Tet4-Pyr showed self-degradation in biological systems. To address this, we have developed 3,5 di-, and 3,4,5 tri-fluoro phenyl derivatives of Tet4 and incorporated using our evolved synthetases. These Tet derivatives achieve a competitive reaction rate while preserving stability within proteins. Eukaryotic compatible Tet-ncAAs encoding systems were also evaluated for encoding efficiency, selectivity, and in-cell labeling in mammalian cells. This work also provides a general strategy for evaluating four key attributes useful to consider when developing ncAAs for GCE-encoded bioorthogonal ligations: synthetic accessibility, encodability, stability, and labeling rate.

## Results

### Strategy

The effectiveness of a synthetically accessible ncAA for GCE-based bioorthogonal ligation depends on its encodability as well as its stability and its on-protein stability and labeling rate, and the tools we use in this study to assess these properties are generally applicable for determining the critical encoding and reaction properties of any new Tet-ncAA. To assess the encoding of Tet-ncAAs (Figures 2, 3, and 9; labeled with color), we use our standard permissivity assay^[9]^ in which Tet-tRNA/RS pairs are evaluated in *E. coli* for their ncAA-dependent suppression of a TAG codon at the Asn150 site in the superfolder green fluorescent protein (sfGFP) gene. Successful encoding of a Tet-ncAA will lead to full-length sfGFP with the ncAA at position 150 (sfGFP^150^), so comparing the sfGFP^150^ fluorescence produced with and without ncAA present quantifies how well that ncAA is incorporated. It is important to note that Tet-labeling is not limited to GFP, it is also encodable and reactive in many other proteins as demonstrated by previous work.^[27–35]^ Here, we categorize the incorporation efficiencies into four levels based on how they compare with expression levels achieved using the ncAA for which the RS was selected: “good efficiency” means similar to the reference ncAA; “moderate efficiency” means ∼30-60% that of reference ncAA; “low efficiency” means still lower expression; and “no detectable encoding” means expression levels indistinguishable from that without ncAA supplementation. We also use Tet-sfGFP^150^ fluorescence to evaluate encoding efficiencies in HEK293 cells, and the ratio of sfGFP^150^ fluorescence to dye fluorescence gives a readout for the in-cell reactivities of expressed Tet-proteins with TCO-dyes.^[9]^

**Figure 2:**
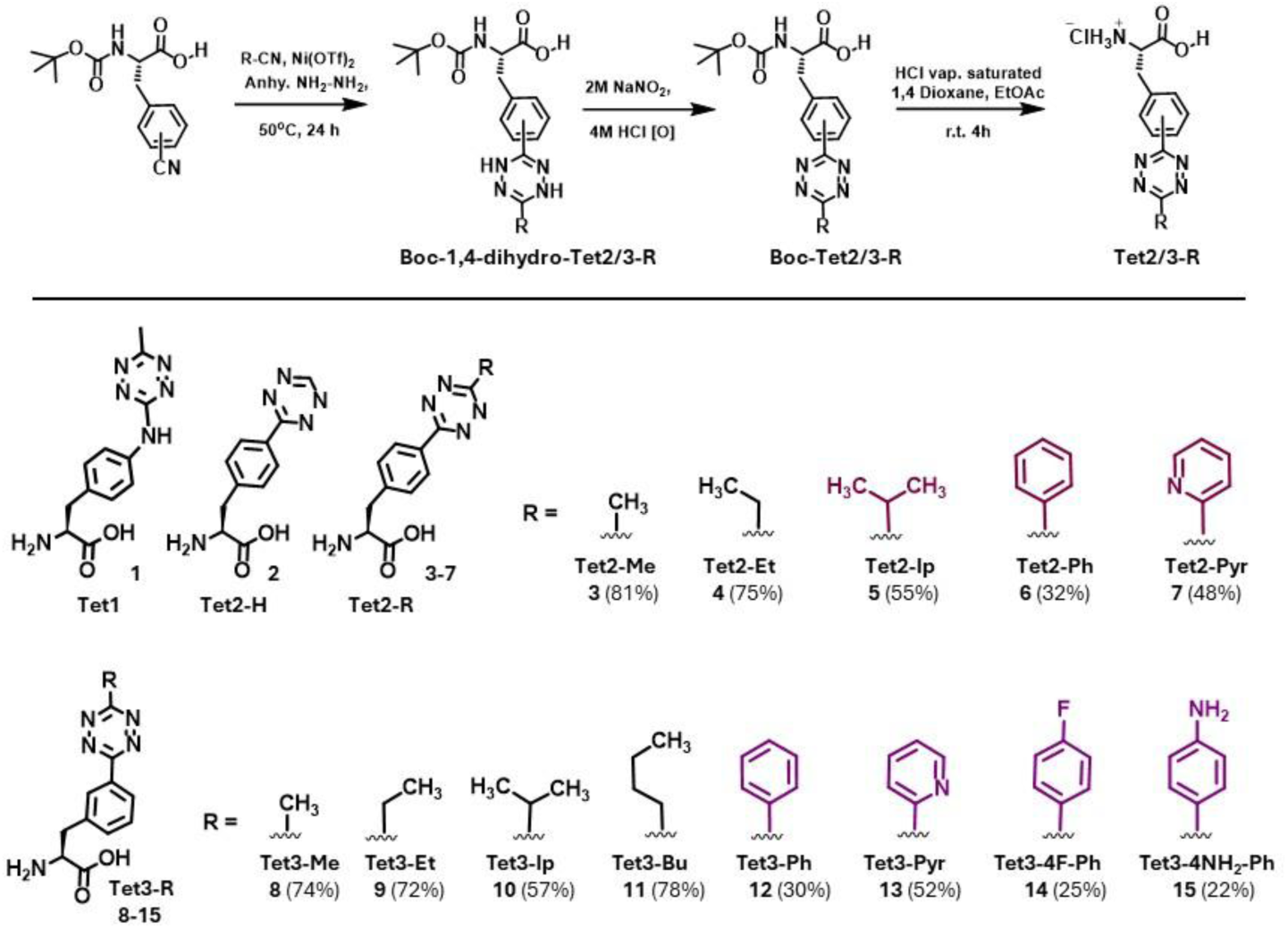
Synthetic scheme of Tet2- and Tet3-ncAA derivatives. Synthetic yields are given in parentheses, and novel derivatives are shown in color.

**Figure 3:**
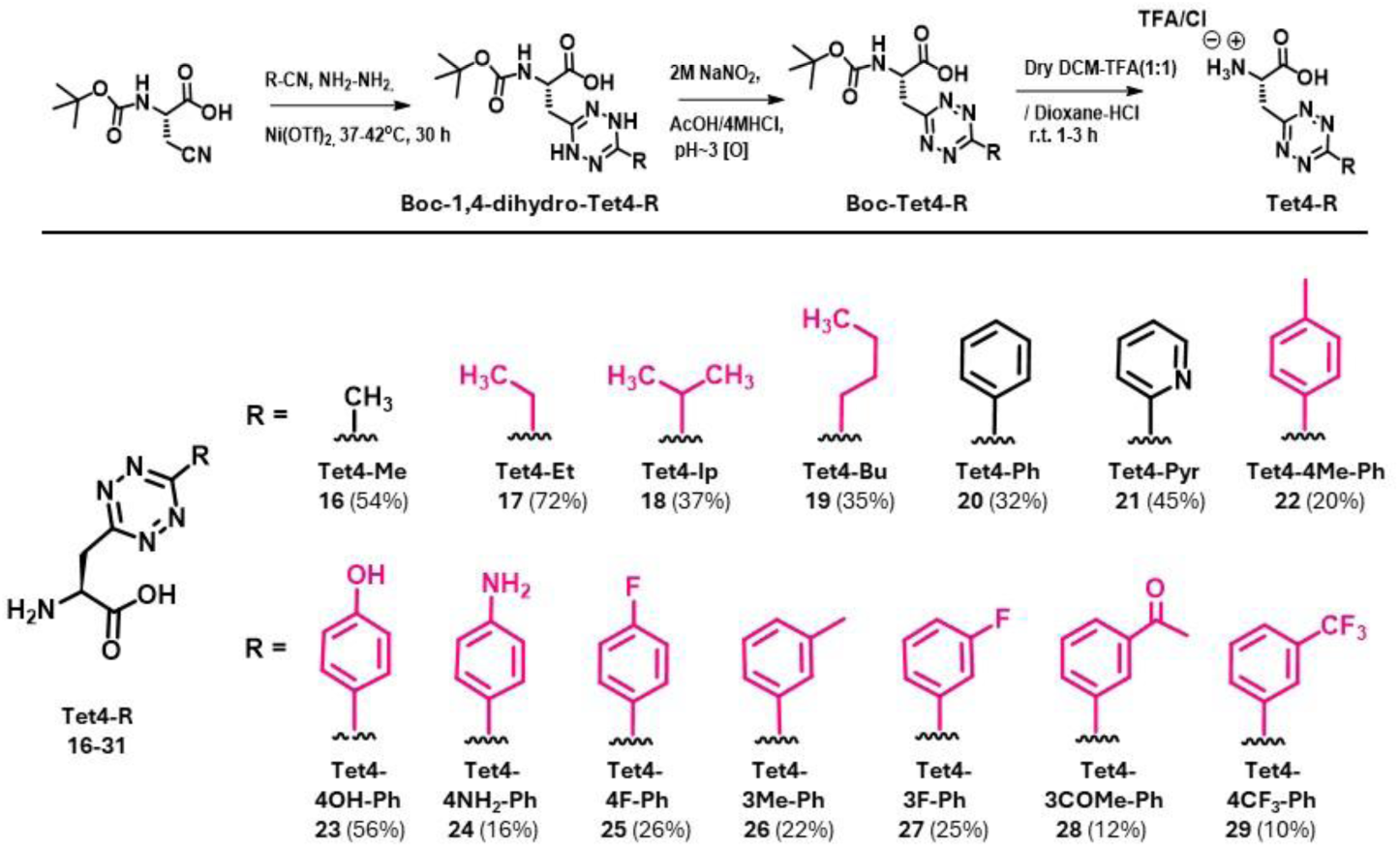
Synthetic scheme of Tet4-ncAA derivatives. Synthetic yields are given in parentheses, and novel derivatives are shown in color.

Since the full-length sfGFP^150^ produced may not all have the intact target ncAA at position 150, we next quantify the extent of on-protein Tet reactivity using a semi-quantitative mobility shift assay. The presence of a C-terminal His-tag on the sfGFP reporter allows all full-length protein to be easily purified, and when this pure protein is reacted with sTCO-PEG5k, the reactive portion of the sample undergoes a mobility shift in SDS-PAGE.^[35]^ Thus, densitometry analysis of the gel gives the reactive Tet-containing fraction of the *E. coli* produced protein. In our experience, this mobility shift assay is more sensitive than mass spectrometry (MS) at detecting trace levels (<10%) of unreactive protein, especially if multiple contaminants are present. This gel shift assay can also be used to assess the stability of purified Tet-protein or the resulting protein-peg conjugate that has been incubated at different conditions before or after reaction with sTCO-PEG5k. Complementing these reactivity-based analyses, MS of full-length Tet-sfGFP^150^ before or after reaction with an sTCO-OH provides a quantitative assessment of Tet-ncAA encoding and reactivity and allows us to identify the molecular nature of significant contaminating species.

Finally, it is on-protein IEDDA reaction rates that are relevant to the utility of the Tet-ncAAs, and we and others have demonstrated that this class of bioorthogonal reactions is significantly influenced by solvent polarity^[36]^ and incorporation into proteins.^[17]^ Our strategy to measure the on-protein reaction rates of the encoded Tet-ncAAs uses analysis of the fluorescence increase always associated with the Tet-sfGFP^150^ reaction with a dienophile, due to relief of the partial quenching of GFP fluorescence. This is possible, because when Tet-amino acids are incorporated at sfGFP residue 150, the GFP fluorescence is quenched by ∼80%, and this is recovered upon reaction with sTCO.^[9]^

### Selection and synthesis of Tet-ncAA derivatives

Since the major application we are seeking to improve the labeling of proteins inside of mammalian cells, in this exploration we focused more on Tet3 and Tet4 derivatives, because the RSs for those two structural scaffolds function in mammalian cells. As an initial set of derivatives to explore for all three scaffolds, we included three aliphatic substituents methyl, ethyl, isopropyl and two aromatic substituents phenyl and pyridyl. This allows the behavior of these five substituents to be compared across all of the scaffolds. Then for Tet3 and Tet4 scaffolds, we also included butyl, 4-fluoro-phenyl and 4-amino-phenyl derivatives. And finally, for the Tet4 scaffold, we added six further phenyl derivatives with varying geometries and electron donating/withdrawing properties; these were the 4-hydroxy, 4-methyl, 3-methyl, 3-fluoro, 3-acetyl, and 3-trifluoromethyl derivatives (Figure 2 and 3).

For efficient synthesis of these diverse Tet2, Tet3 and Tet4 ncAA derivatives, we adapted the method from Devaraj et al.^[37]^ for a metal-catalyzed coupling between two nitriles in the presence of hydrazine. Using Ni(OTf)_2_-as a catalyst,^[9]^ the Tet2- and the Tet3-ncAAs were synthesized in 22-81% yields by the addition of hydrazine to boc-4-nitrile phenylalanine and boc-3-nitrile phenylalanine, respectively, in presence of various nitrile derivatives of the targeted substituents (Figure 2). For the smaller Tet4 scaffold, 10-50% yields were obtained for reacting diverse nitrile-substituents with boc-nitrile-alanine (Figure 3). Among the derivatives synthesized for this study, the ones not tested before for incorporation into proteins were three of the Tet2 derivatives **5-7** (Figure 2), four of the Tet3 derivatives **12-15** (Figure 2), and eleven of the Tet4 derivatives **17-19** and **22-29** (Figure 4).

**Figure 4:**
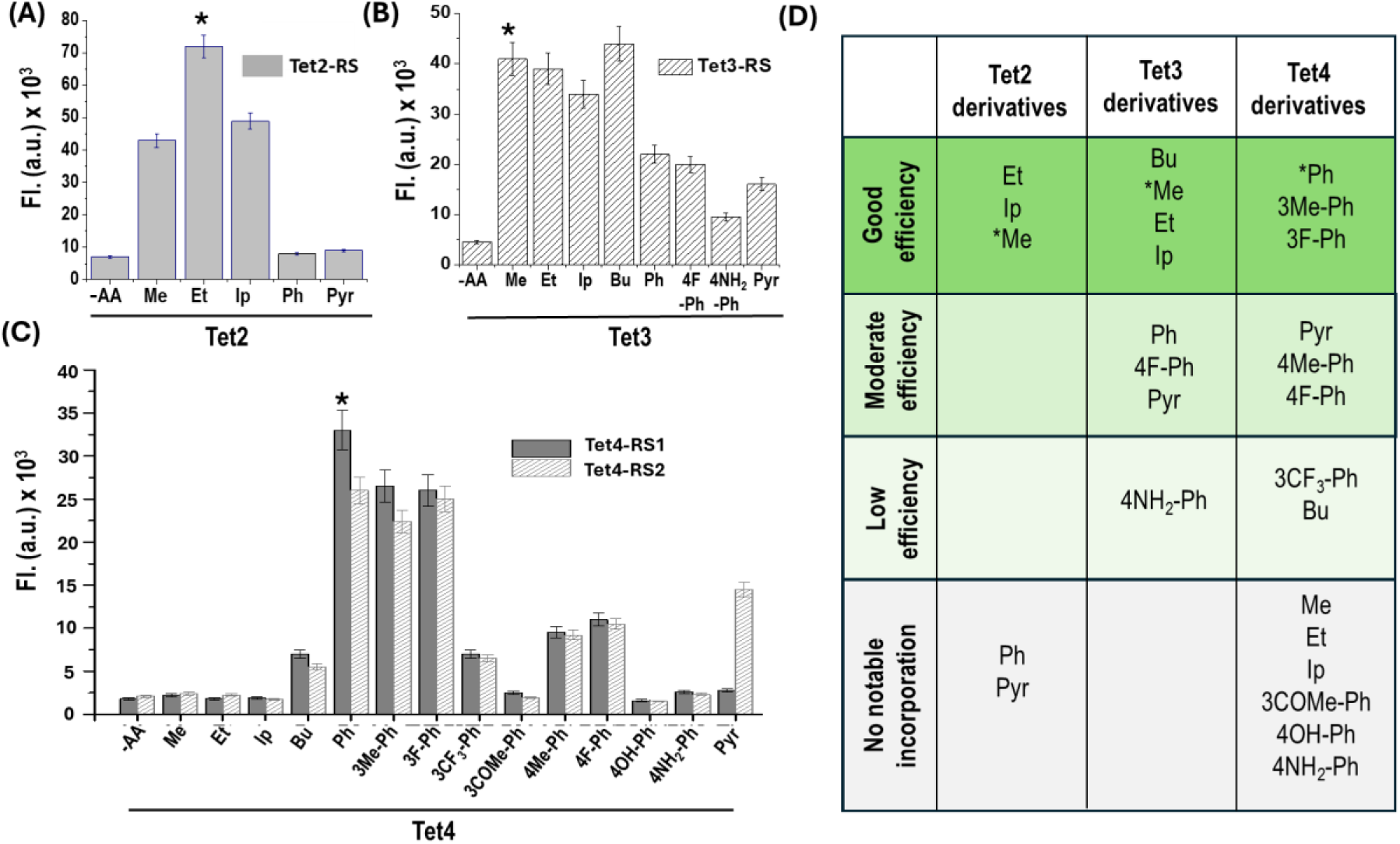
Genetic incorporation of Tet-ncAAs. Permissibility and efficiency of evolved synthetases measured by florescence of expressed sfGFP-TAG150 in absence and presence of Tet-ncAA. (A) Evaluation of Tet2 (D12) synthetase^[17]^ with 0.5 mM Tet2-ncAAs. (B) Evaluation of Tet-3 (R2-84) synthetase^[9]^ with 0.5 mM Tet3-ncAAs. (C) Evaluation of Tet4-RS1 (E1) and Tet4-RS2 (D4) synthetases^[18]^ with 0.3 mM Tet4-ncAAs. All expressions are in 25 mL AIM at 37 °C for 32 hrs. (D) Overview table of encoded Tet2, Tet3 and Tet4 derivatives. Asterisks (*) represents Tet-ncAAs used to evolve orthogonal aaRS/tRNA_CUA_ pair.

### Genetic incorporation Tet-ncAAs into protein

#### Toxicity screens

As the cellular toxicity of the ncAA can hinder its incorporation into proteins, we assessed the impact of alkyl and phenyl derivatives of the synthesized Tet-ncAAs on the viability of the *E. coli* DH10B cells to be used in test expressions (see SI, Figures S1-4). Tests at concentrations of up to 1 mM revealed that Tet-alkyl derivatives had minimal effects on cell growth. For all the Tet-Pyridyl compounds and the Tet4-Aryl scaffold, significant toxicity was observed at 1 mM concentration, and for these derivatives toxicity was minimized by lowering concentrations (0.3 mM) of Tet-ncAA, and adding the Tet-ncAA to the media 4 to 6 h after cell inoculation, once the cells reached a healthy growth phase (OD_600_ ∼ 0.5 to 0.7) (Figures S3 and S4).

#### Tet-ncAAs incorporation

To screen the existing Tet2, Tet3 and Tet4 RSs for their ability to incorporate into proteins the wider range of Tet-ncAA derivatives, we used our standard fluorescence-based assay^[9]^ as discussed above in the Strategy section. The results of the permissivity screens for each Tet scaffold are summarized in Figure 4D. For the Tet2 scaffold, all alkyl derivatives of Tet2 incorporated well, on par with or better than the methyl derivative against which the RS was selected; in contrast, the fidelity ratios for both aromatic derivatives **6** and **7** (Tet2-Ph and Tet2-Pyr) were near 1, indicating no useful incorporation (Figure 4A; TableS1). For the Tet3 scaffold, the Tet3-RS incorporated all derivatives tested, with good efficiency for the alkyl derivatives **9**-**11** (Tet3-Et, -Ip, and -Bu), moderate efficiency for three of the aryl derivatives **12**-**14** (Tet3-Ph, -Pyr, and −4F-Ph), and low efficiency for **15** (Tet3-4NH_2_-Ph) (Figure 4B). For the Tet4 derivatives, both the Tet4-RS1 and Tet4-RS2 were tested, as it is well-documented that RSs that work similarly for the ncAA against which they were selected can have very different permissivity profiles.^[9,18]^ In addition to efficiently encoding Tet4-Ph, against which the RSs were selected, both Tet4-RSs encoded two of the new Tet4 derivatives with high efficiency (3-Me-Ph and 3F-Ph) and four new Tet4 with moderate to low efficiency (4Me-Ph (**22**), 4F-Ph (**25**), 3CF_3_-Ph (**29**) and butyl (**19**)), while neither Tet4-RS incorporated the **16**-**18** (Tet4-Me, -Et, and -Ip) derivatives or the Tet4-aryl −4OH (**23**), −4NH_2_ (**24**) or −3COMe (**28**) derivatives (Figure 4C). In addition, Tet4-RS2 uniquely incorporated Tet4-Pyr (**21**).

Next, we carried out mass spectrometry (MS) on some of the purified sfGFP^150^ proteins to directly assess the ncAA incorporation fidelities (Figures 5B-D, S8 and Tables S4A-B). All the ncAAs that gave high-efficiency expression exhibited a single prominent peak consistent with incorporation of the respective Tet-ncAA at position 150, with no evidence of misincorporation. These were: Tet2-Me and -Et (**3**, **5**); Tet3-Me, -Et, and -Bu (**8**, **9** and **11**); and Tet4-Ph, −3Me-Ph, −3F-Ph (**20** and **26**-**27**). All the other ncAAs assessed (Tet3-Ph, −4F-Ph, −4NH_2_-Ph and -Pyr (**12**-**15**) as well as Tet4-3CF_3_-Ph and -Pyr (**29**-**21**)) had peaks in the MS that indicated between 10 and 50% of the protein produced incorporated something other than the ncAA (Figures 5D, S8 and Tables S1-3). In all cases, the alternate mass was consistent with the incorporation of glutamine or lysine, so we infer that the misincorporation is due to near-cognate suppression (NCS) which is a well-known process that can lead to glutamine insertion at TAG codons.^[35,38]^ The level of misincorporation was generally anti-correlated with the efficiency of protein expression, with the Tet3-4NH_2_-Ph having the lowest efficiency sfGFP150 production and, at ∼50%, the highest level of misincorporation due to NCS (Figure S8C, and Table 2).

**Figure 5:**
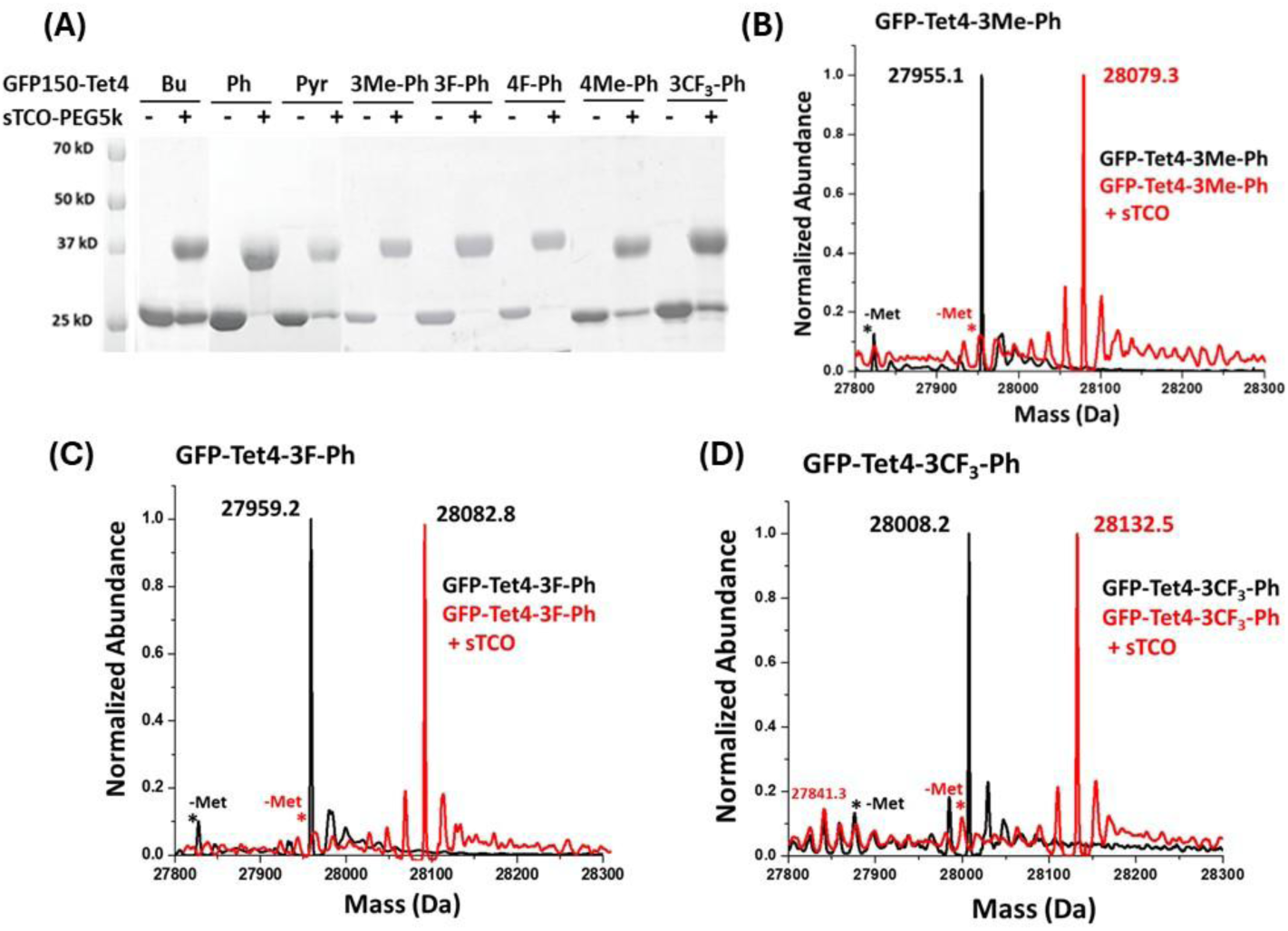
SDS-page gel mobility shift assay and MS analysis confirmed Tet4 derivatives incorporation into GFP and the labeling efficiency with sTCO reagent. (A) Fidelity of Tet4 incorporation examined by reacting with sTCO-PEG5k and the mobility shift of the reacted proteins in SDS-page gel. ESI mass spectrometry analysis of purified GFP-Tet4 derivatives (black) and the reaction with 5-fold molar excess of sTCO for 10 minutes (red) shows as expected 124 Da increase in mass corresponding to the addition of sTCO and loss of molecular nitrogen. No unreacted sfGFP-Tet4 were detected, verifying the reaction of genetically encoded Tet4 derivatives with sTCO was quantitative. Cal. Mass of sfGFP-wt: 27827.1 Da avg; (B) sfGFP-Tet4-3Me-Ph observed: 27955.1 Da avg, (expected: 27954.1 Da avg); sfGFP-Tet4-3Me-Ph + sTCO observed: 28079.3 Da avg, (expected: 28078.2 Da avg). (C) sfGFP-Tet4-3F-Ph observed 27959.2 Da avg (expected: 27958.05 Da avg.); sfGFP-Tet4-3F-Ph + sTCO observed: 28082.8 Da avg. (expected: 28082.1 Da avg.). (D) sfGFP-Tet4-3CF_3_-Ph observed 28008.2 Da avg (expected: 28008.1 Da avg.) sfGFP-Tet4-3CF_3_-Ph + sTCO observed: 28132.5 Da avg. (expected: 28132.1 Da avg.). The peak at 27841.3 Da avg. observed due to near-cognate suppression of amber codon. The lower mass peak labeled with asterisk (*) is a loss of n-terminal methionine (-Met) and upper mass peaks are salt of sodium and potassium adducts.

In summary, we have found that the existing Tet ncAA RSs can encode one, four and six of the newly tested Tet2 (**5**), Tet3 (**12**-**15**) and Tet4 ncAAs (**19**, **22**, **25**-**27**, and **29**) into proteins. And even while some of these ncAAs are incorporated along with some misincorporation due to NCS, these results provide proof of principle that these ncAAs can be incorporated via translation into proteins. Worth noting is that we and others have shown that because NCS misincorporation occurs in competition with ncAA incorporation, it is often possible to overcome the NCS simply by evolving a more efficient next-generation synthetase.^[35,39]^ Also, important to note that even with contaminating protein present, the stabilities and reactivities of the incorporated Tet residues can still be characterized, because the protein containing a misincorporated glutamine in place of the Tet-ncAA will not react with the TCO reagent.

### Chemical properties of the Tet-containing proteins

#### On-protein Tet-ncAA reactivity

Because the utility of a biorthogonal labeling group depends on its stability and reactivity when encoded onto proteins, we next studied the on-protein reactivity. The various purified Tet-sfGFP^150^ were evaluated based on their reaction with sTCO-PEG5k, which resulted in a mobility shift in SDS-PAGE gel, as discussed in the strategy section.

For Tet2-scaffold, we previously demonstrated the labeling efficiency of Tet2-Me and Tet2-Et on the protein.^[35]^ Tet2-Ip showed similar labeling efficiency, likely due to the similar functionality they all share (Figure S5A). For Tet3 and Tet4-derivatives, reactivities for the freshly purified proteins were consistent with the MS analyses of the pure proteins. Those that had shown 100% ncAA incorporation by MS of **11** (Tet3-Bu) and **20**, **25**-**27** Tet4 (Ph, 4F-Ph, 3Me-Ph, 3F-Ph) showed quantitative labeling (>90%) with sTCO-PEG5k, while the Tet3 and Tet4 derivatives which showed NCS contamination (Tet3-Ph (**12**), -Pyr (**13**), −4NH_2_-Ph (**15**) and Tet4-Bu (**19**), -Pyr (**21**), −4Me-Ph (**22**), −3CF_3_-Ph (**29**)) also showed unreacted bands at levels 3 – 15% higher than was indicated by MS (Figures 5A, S5B and Tables S2-3). As in the MS analysis, the ncAA leading to the most unreactive protein was Tet3-4NH_2_-Ph (**15**), which exhibited 51% unreactive protein by MS and 57% unreactive protein in the mobility-shift assay.

Mass spectrometry analysis of sTCO-OH labeling reactions showed full reactivity of the Tet-containing proteins, with the entire peaks corresponding to Tet3 and Tet4-ncAA proteins shifting 124.2 Da higher, indicative of sTCO addition with the loss of N_2_. And as expected, the peaks inferred to be protein with glutamine incorporated (produced by NCS) were unaltered (Figures 5B-D, S8 and Tables S4A-B). No additional peaks were detected, confirming the quantitative reaction of each Tet3 and Tet4-sfGFP^150^ with sTCO without side reactions.

#### On-protein Tet-ncAA stability

We next assessed how well the Tet3 and Tet4 ncAAs, after incorporation into the protein, maintained their reactivity over time. As it is known that stability is anti-correlated with reactivity, we focused our studies on the sfGFP^150^ proteins with the electron-withdrawing Pyr derivatives of Tet3 and Tet4, which we expect to be the most reactive, along with the electronically neutral Ph derivative of each Tet (Tet2-Tet4), and also including four Tet4 derivatives with different aryl substitutions – 4Me-Ph (**25**), 3Me-Ph (**26**), 3F-Ph (**27**) and 3CF_3_-Ph (**29**). These proteins were incubated for 8 days in a neutral buffer at 4 °C and at room temperature (RT), and the gel-shift assay was used to test their reactivity. Tet-sfGFP incubated at 4 °C, over the 8 days, none of the tested proteins lost any reactivity with sTCO-PEG5k, and even at RT, for 3 days of incubation, there was no notable degradation for any of the Tet3- and Tet4-phenyl derivatives were observed whereas, upon 8-days of incubation, ∼20% of the protein was found degraded (Figures 6C, S6B and S7). However, for both derivatives with the electron-withdrawing pyridyl moiety, degradation was much more rapid, and approximately 10% reactivity was observed after 5 days (Figures 6D and S6B).

**Figure 6:**
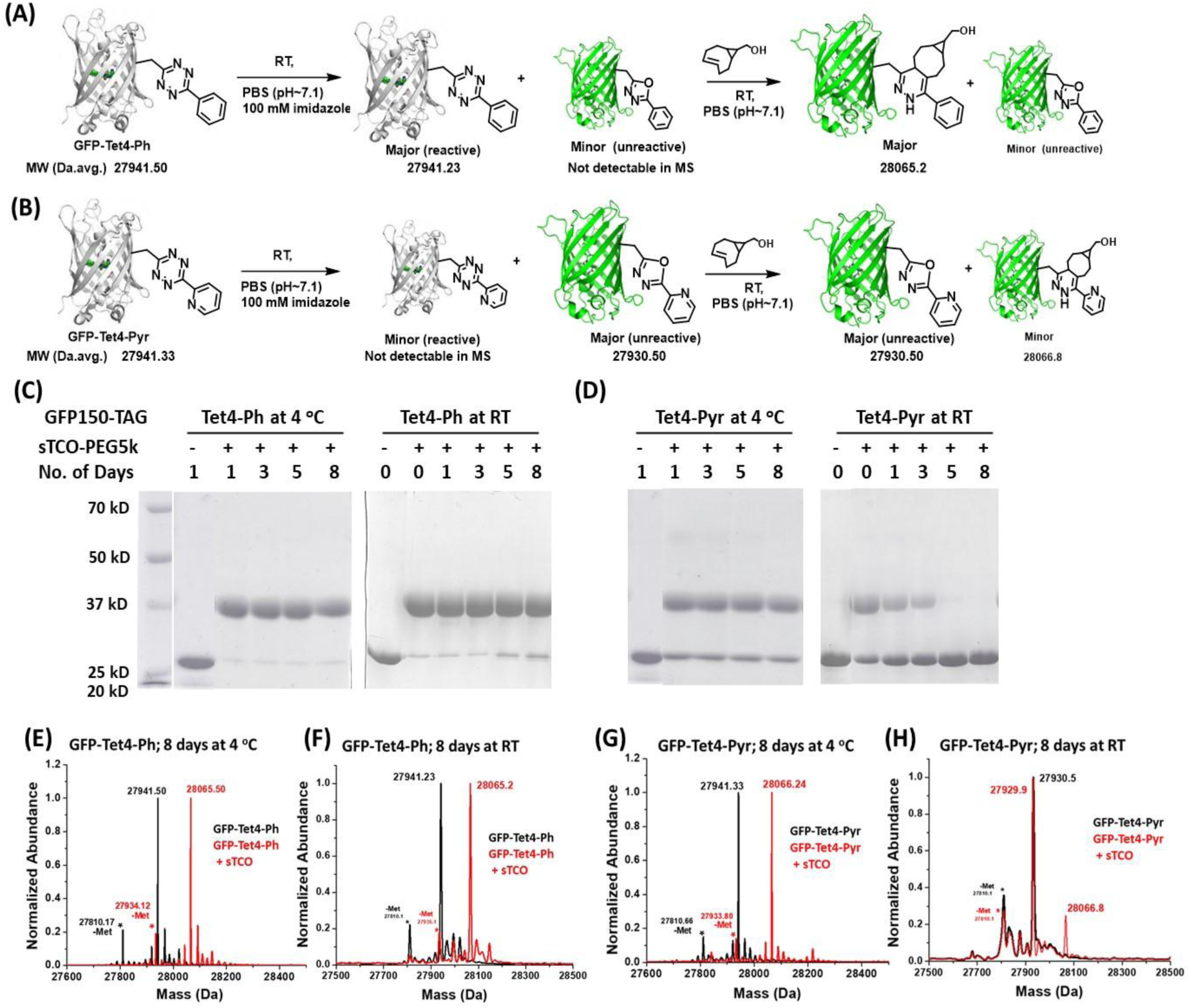
Assessing the stability of Tet4-Ph and Tet4-Pyr in a protein context. (A) and (B). Schematic of the RT stability assay results for sfGFP^150^-Tet4-Ph and sfGFP^150^-Tet4-Pyr, respectively. (C) and (D). SDS-PAGE gel of mobility shifts seen at time points taken during the 4 °C and RT incubations of sfGFP^150^ containing Tet4-Ph (**20**) and Tet4-Pyr (**21**), respectively. (E) through (H). ESI-Q-TOF mass spectrometry of purified sfGFP-Tet4-Ph/Pyr (black) and upon reaction with 5-fold molar excess of sTCO-OH for 10 minutes (red) after 8 days at 4 °C and RT as indicated in the figure reacts quantitively with sTCO; sfGFP-Tet4-Ph observed: 27941.23 Da avg, (expected: 27940.1 Da avg); sfGFP-Tet4-Ph + sTCO observed: 28065.2 Da avg, (expected: 28064.12 Da avg). whereas (G) at 4°C, sfGFP-Tet4-Pyr observed 27941.33 Da avg (expected: 27941.1 Da avg.) sfGFP-Tet-v4.0Pyr+ sTCO-OH observed: 28066.24 Da avg. (expected: 28065.11 Da avg.) (H) sfGFP-Tet4-Pyr degraded with time at room temperature. There was a small peak observed for a reaction with sTCO and the majority of sfGFP-Tet4-Pyr remains unreactive. sfGFP150-Tet4-Pyr observed 27930.5 Da avg (expected: 27941.1 Da avg.) sfGFP-Tet4-Pyr + sTCO observed: 27929.9 (unreactive) and 28066.76 Da avg. (expected: 28065.11 Da avg.) The unreacted single major peak at 27930.5 Da avg. indicates that the sfGFP-Tet4-Pyr degraded under the following conditions and converted to its oxadiazole derivative which is 11 Da lower molecular mass than Tet4-Pyr. Note: Figures (E) and (G) contain MS data at 4°C, which were sourced from reference 18.

To characterize the degradation products of the Tet-Ph and Tet-Pyr scaffolds, the samples after 8-day incubation were reacted with sTCO-OH and analyzed by MS. For the 4 °C and RT incubations of the Tet3-Ph (**12**) and Tet4-Ph (**20**) proteins as well as the 4 °C incubations of the Tet3-Pyr (**13**) and Tet4-Pyr (**21**) proteins, a dominant single mass peak corresponding to the sTCO adduct was observed, indicating that the level of degradation that occurred was below the limit of detection by MS (Figures 6E-G and S9A). However, at the RT incubation of the Tet3-Pyr (**13**) and Tet4-Pyr (**21**) proteins, only a minor peak represented the sTCO adduct and a major peak 11 Da less than the expected mass of the unreacted protein observed which represented the 1,3,4-oxadiazol breakdown product (Figures 6H and S9B and Tables S4C-D).

The loss of 11 Da is consistent with the tetrazines being converted to their oxadiazole derivative (Figures 6A-B, S10), which is a well-known breakdown product^[40]^ and is not reactive to sTCO. These findings conclusively demonstrate that this range of Tet2, Tet3, Tet4 alkyl and aryl derivatives are stable enough for protein labeling over a period of multiple days at RT, whereas the pyridyl-attached Tet scaffolds are labile enough that they must be kept cold or used within a few hours to maintain its full reactivity.

#### On-protein Tet-ncAA kinetics upon reaction with sTCO

Since the utility of the Tet derivatives for various applications, particularly for fast and complete labeling at low concentrations, depends on their reaction speeds, we measured the kinetics of reactions with sTCO-OH for some of the Tet-ncAAs in both their free and on protein contexts. For a few selected free Tet-ncAA from each of the Tet2, Tet3 and Tet4 sets, kinetics was measured using stopped-flow spectroscopy to monitor tetrazine absorbance at 270 nm. The reaction kinetics in a protein context were measured by the time course of increasing sfGFP^150^ fluorescence during reaction with sTCO (Figure S11).^[9]^ Based on our earlier work with Tet2 and Tet3 derivatives, we anticipated observing ∼3-fold enhanced reactivity in the protein context, which we attributed to the hydrophobic local reaction environment provided by the protein surface.^[36]^

The free Tet2 and Tet3 ncAAs tested exhibited second-order rate constants (*k*_2_) ranging from about 0.5 – 20 × 10^4^ M^-1^ s^-1^ (Tables S1 and S2). Notably, Tet3-Ip (**11**) was the slowest reacting, presumably due to steric hindrance, and Tet2-Pyr (**7**) and Tet3-Pyr (**13**), with their electron-withdrawing pyridyl groups, were the fastest with *k*_2_ values of 20.4 × 10^4^ M^-1^ s^-1^ and 9.6 ×10^4^ M^-1^ s^-1^, respectively. In a protein context, the Tet2 and Tet3 derivatives generally reacted more rapidly, with second-order rate constants ranging from 2 – 23 × 10^4^ M^-1^ s^-1^. Despite their structural differences, Tet2 and Tet3 exhibited similar reactivity (Figure 7, Figure S12).

**Figure 7:**
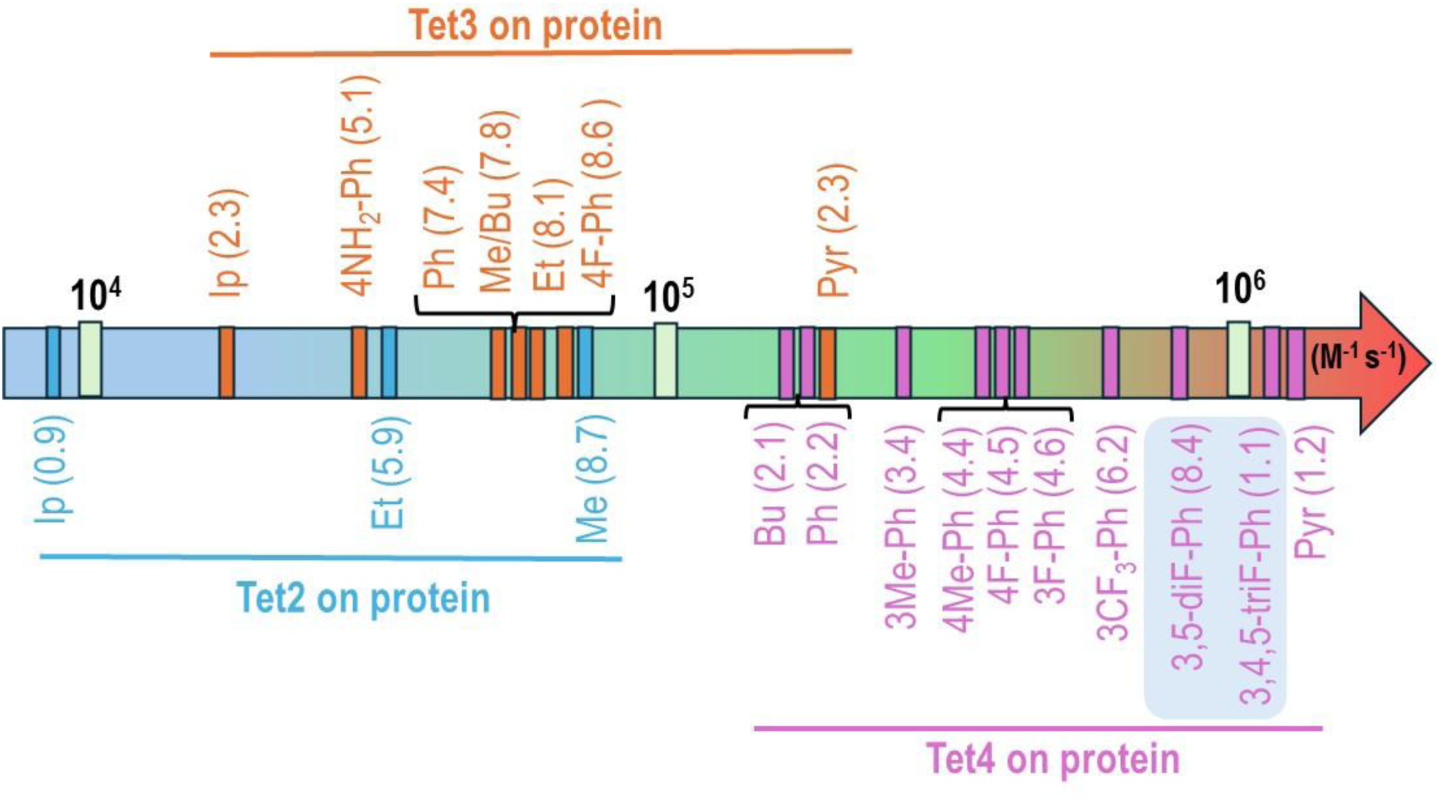
On-protein reaction rates of Tet-ncAAs upon reaction with sTCO. Increasing order of second-order rate constants (*k*_2_ with units of M^-1^ s^-1^) of encoded Tet2, Tet3 and Tet4 derivatives into sfGFP^150^ with sTCO-OH in PBS, pH 7.0, at room temperature. The two highlighted Tet4 derivatives (light blue background) were a second set of variants designed to have very high reactivity and sufficient stability for in-cell studies.

Furthermore, free-Tet4-ncAAs derivatives also exhibited a broad range of kinetics, spanning from 1 – 8 × 10^4^ M^-1^ s^-1^. Tet-ncAA including, Tet4-Ph (**20**), Tet4-4NH_2_-Ph (**24**), Tet4-4F-Ph (**25**), displayed a good reaction rate of ∼2.4 × 10^4^ M^-1^ s^-1^, while Tet4-Pyr (**21**) was 4-fold faster at 8.5 × 10^4^ M^-1^ s^-1^. Particularly striking about the Tet4-derivatives, is that all three for which both free and on-protein reactivities were determined exhibited a 10-20 -fold enhancement, not just the 3-fold on-protein rate enhancement we have seen for Tet2 and Tet3 on proteins. (Figures 7, S13, and Table S3). We hypothesize that the observed rate enhancement for the shorter Tet4 ncAAs compared to Tet2 and Tet3 ncAAs is due to the tetrazine group’s closer proximity to the protein surface and that preorganization of the Tet reduces the entropy of activation for protein surface IEDDA reactions. While the rates of IEDDA reactions between tetrazines and TCOs are well documented increase by using more polar solvents,^[41]^ the polarity of the environment is not predicted to change significantly between Tet ncAAs free in solution as compared to the longer Tet2/3 on proteins and shorter Tet4 on proteins. Entropy has been identified as the major contributor to free energy of activation for Tet/TCO reaction.^[42,43]^ A similar ∼10 fold rate increase was observed when Tet-glycan was integrated into the peptidoglycan surface of bacteria as compared to the free Tet-glycan in solution.^[44]^ Additionally, this on-protein preorganization hypothesis matches the ground state distortion model by Mikula, which identifies how prepositioning the ground state in a transition state like geometry can accelerate IEDDA reactions.^[45][43]^ This on-protein preorganization explanation for rate increase also matches our previous study, where nearly identical Tet4-Ph (**20**) reaction rates were obtained on two different locations on sfGFP, one in a positively charged environment and one a neutral electrostatic environment.^[18]^ This means that it is not the intrinsic reactivities of the Tet4 ncAAs, but this “on-protein” boost which allows the Bu (**19**), Ph (**20**), 3Me-Ph (**26**), 4Me-Ph (**22**), 3F-Ph (**27**), 4F-Ph (**25**), 3CF_3_-Ph (**29**) derivatives of Tet4 to achieve their unusually fast reactivities ranging from 21 – 62 × 10^4^ M^-1^ s^-1^, and allows theTet4-pyridyl derivative to achieve its ultra-fast *k*_2_ value of 120 × 10^4^ M^-1^ s^-1^.

### Expression and labeling of Tet-proteins within eukaryotic cells

All the above work involved expressing Tet-containing sfGFP^150^ in *E. coli*. To test the potential utility of the newly developed range of rapidly-reacting Tet3 and Tet4 derivatives for live-cell labeling work in mammalian cells, we followed our previously optimized eukaryotic expression protocol for producing ncAA-sfGFP^150^ in HEK293T cells, where we found that cell viability was compromised at Tet-ncAA concentrations above 0.1 mM.^[9]^ Moderate to high fidelity expressing three Tet3 derivatives (Ph (**12**), 4-F-Ph (**14**), Pyr (**13**)) and five Tet4 derivatives (Ph (**20**), 3Me-Ph (**26**), 3F-Ph (**27**), 4F-Ph (**25**), and Pyr (**21**)) were tested in HEK cells, and all were successfully incorporated into sfGFP^150^ using Tet3-RS and both the Tet4-RS1 or Tet4-RS2 synthetases, respectively (Figures 8A and 8C). Impressively, it was observed that R284 efficiently encoded Tet3-Ph (**12**) and Tet3-Pyr (**13**) at similar levels to Tet3-Bu (**11**) in HEK293T cells, comparable to the results in *E. coli*. A surprise result is that Tet4-RS1, which is unable to incorporate Tet4-Pyr (**21**) in *E. coli*, does encode it in HEK cells. Additionally, Tet4-RS2 demonstrates much better encoding of Tet4-Pyr (**21**) in HEK cells compared to in *E. coli*. We do not know the reason for this difference, but one simple explanation would be if Tet4-Pyr (**21**) enters HEK cells better than it enters *E. coli* or has higher stability in mammalian culture conditions.

**Figure 8.**
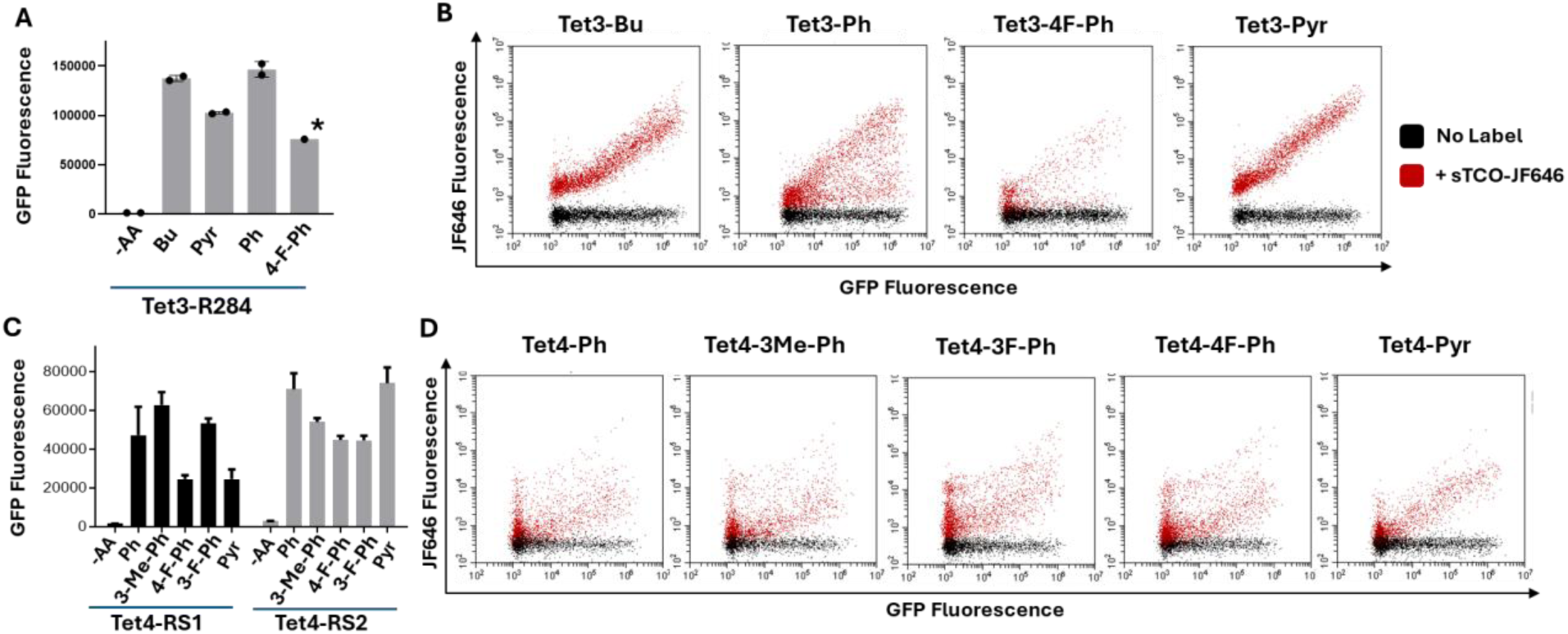
Expression and labeling of Tet3 proteins in mammalian HEK293T cells analyzed by FACS. (A) Efficiency of the R284 Tet3 aaRS/tRNA system for encoding Tet3 variants (30 µM), indicated by the fluorescence of sfGFP^150^-Tet3 (*expressed individually). (B) Live in cell labeling of Tet3 expressions. Cells were incubated with sTCO-JF646 (100 nM) for labeling. Tet3-Bu (**11**) and Tet3-Pyr (**13**) were labeled for 60 min prior to analysis and Tet3-Ph (**12**) and Tet3-4F-Ph (**14**) for 90 min. (allowed 30 more minutes for complete labeling to relatively less reactive Tet3-ncAA) (C) Efficiency of the Tet4-RS1(E1) and Tet4-RS2 (D4) aaRS/tRNA systems for encoding Tet4 variants (100 µM), indicated by the fluorescence of sfGFP^150^-Tet4. (D) Live in cell labeling of Tet4 expressions using Tet4-RS2. Cells were incubated with sTCO-JF646 (100 nM) for 60 min prior to analysis. Expressions were performed for 24 h before labeling assays and FACS measurements.

Next, the HEK293T cells expressing sfGFP^150^-Tet derivatives were allowed to react with the cell-permeable fluorescent dye sTCO-JF646,^[18]^ and labeled cells were tracked by measuring JF-646 fluorescence using flow cytometry (Figures 8B and 8D). The increase in JF-646 fluorescence proportionally to the Tet-sfGFP^150^ fluorescence indicates the successful labeling of the expressed Tet-sfGFP^150^, with minimal non-specific labeling. Notably, the sfGFP^150^ with Tet3-Pyr (**13**), Tet4-3F-Ph (**27**) and Tet4-Pyr (**21**) showed both greater and more consistent labeling as indicated by a steeper diagonal distribution, as expected based on their higher reactivities. We conclude that all these Tet3 and Tet4 derivatives are of potential use for the in-cell labeling of eukaryotic proteins.

### Two additional Tet4-derivatives designed to combine high incorporation efficiency, high stability, and high reactivity

#### Design, synthesis and encoding of Tet4-3,5-diF-Ph (**30**) and Tet4-3,4,5-triF-Ph (**31**)

Given the above results, we wondered if we could use the information gained to create a Tet derivative that would have reactivity similar to the fastest Tet derivative, Tet4-Pyr (**21**), but would maintain sufficient stability to allow for reliable protein labeling over multiple days at room temperature. We noticed that introducing a single fluorine into Tet4-Ph at the 3-position (compound **27)** nearly doubled the reaction rate while maintaining high stability and compatibility with Tet4-RS. Based on this and considering fluorine’s small size and strong electron-withdrawing nature, we hypothesized that di- and tri-fluoro derivatives could be even faster, still relatively stable, and could also be encoded by existing Tet4-synthetases. We designed and synthesized two new Tet4 derivatives: Tet4-3,5-diF-Ph (**30**) and Tet4-3,4,5-triF-Ph (**31**) (Figure 9A).^[46,47]^ An evaluation of their encoding by Tet4-RS1 and Tet4-RS2 showed that Tet4-RS1 incorporated both amino acids very well, at as high or even higher efficiency than the Ph derivative for which the RS was developed (Figure 9B), and also with high fidelity as confirmed by MS (Figure 9D-E). The encoding efficiency of two new Tet4 derivatives was compared to that of Tet4-Ph (**20**) using the previously described in cell eukaryotic assays use for Tet3 and Tet4 studies at a range of ncAA concentrations (30-400 μM) with the expected drop off at above 200 μM due to cellular toxicity (Figure S14A, & B). Good encoding efficiency (80-100%) for Tet4-3,5-diF-Ph (**30**) and moderate efficiency (40-60%) for Tet4-3,4,5-triF-Ph (**31**) was observed.

**Figure 9:**
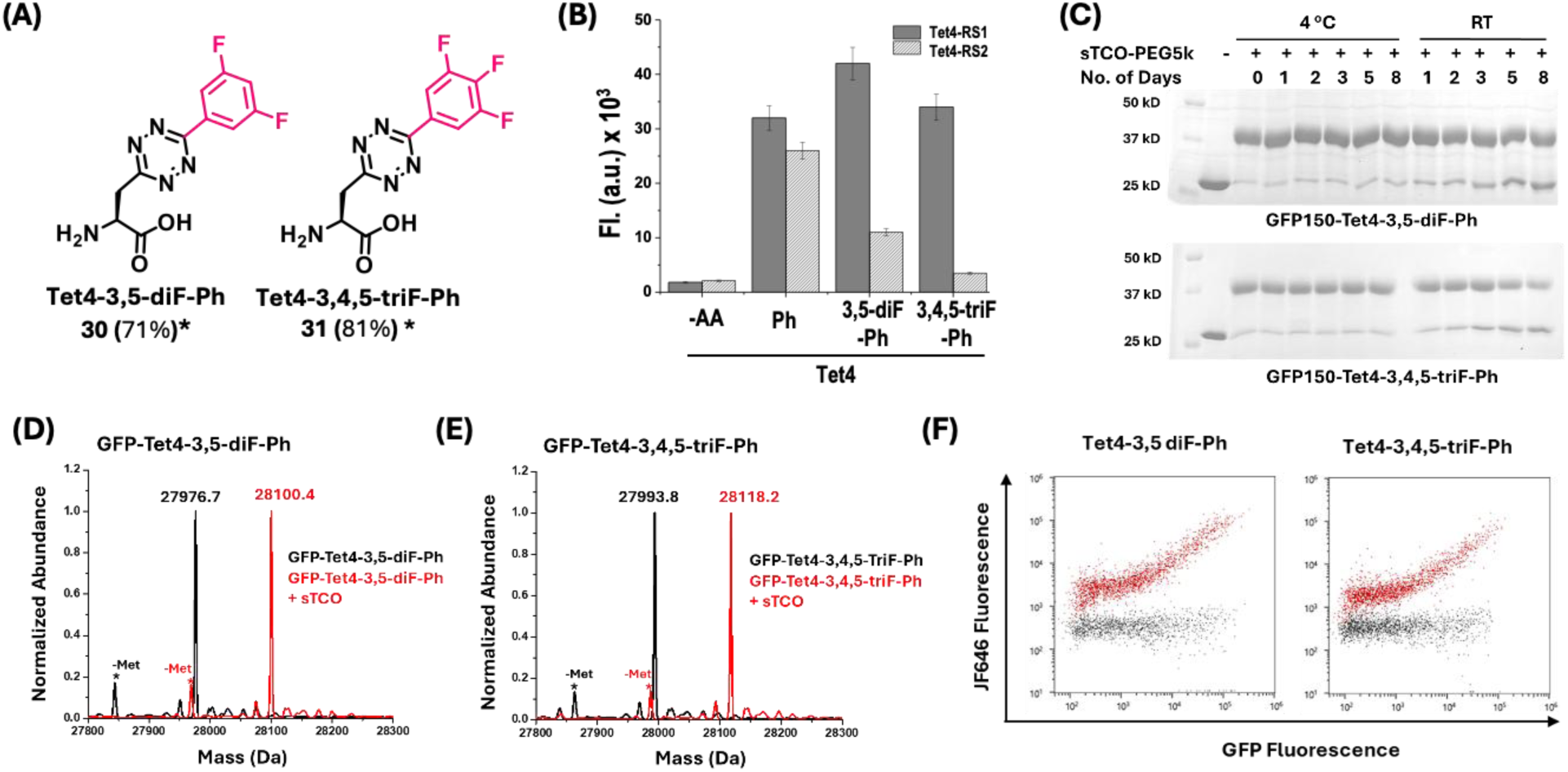
(A) Chemical structure of di-F and tri-F-Tet4-Ph derivatives. Asterisks (*) indicate synthesis performed following reported protocol ^[46,47]^ (Synthetic yields in parenthesis). (B) Permissibility and efficiency of Tet4-synthetases measured by florescence of expressed sfGFP-TAG150 in absence and presence of 0.3 mM Tet4-ncAAs. (C) SDS-PAGE mobility shift assay by reacting sTCO-PEG5k verified the stability and reaction ability of encoded Tet4-3,5-diF-Ph (**30**) and Tet4-3,4,5-triF-Ph (**31**) stored at 4 °C and RT for eight days in PBS (pH∼7.1) at the presence of 100 mM imidazole. ESI mass spectrometry analysis of purified sfGFP-Tet4 derivatives 30, 31 (black) and the reaction with 5-fold molar excess of sTCO for 10 minutes (red) shows as expected 124 Da increase in mass corresponding to the addition of sTCO and loss of molecular nitrogen. No unreacted sfGFP-Tet4 were detected, verifying that reaction with sTCO was quantitative. (D) sfGFP-Tet4-3,5-di-F-Ph observed: 27976.7 Da avg, (expected: 27976.1 Da avg); sfGFP-Tet4-3,5-di-F-Ph + sTCO observed: 28100.4 Da avg, (expected: 28100.1 Da avg). (E) sfGFP-Tet4-3,4,5-tri-F-Ph observed: 27993.8 Da avg, (expected: 27994.1 Da avg); sfGFP-Tet4-3,4,5-tri-F-Ph + sTCO observed: 28118.2 Da avg, (expected: 28118.1 Da avg). The lower mass peak labeled with –Met is a loss of n-terminal methionine. (F) Live in cell labeling of the new Tet4 ncAAs encoded in sfGFP^150^. Cells were incubated with sTCO-JF646 (100 nM) for 60 min prior to analysis. Expressions were performed for 24 h before labeling assays and FACS measurements.

#### Characterization of new Tet4-ncAA labeling, stability and reaction kinetics

The quantitative labeling efficiency of **30** and **31** in a protein context was confirmed by SDS-PAGE gel mobility shift and MS analyses (Figure 9C-E). In terms of their on-protein stability (Figure 9C-E), no degradation was seen at 4 °C just as for all other derivatives, and importantly, at room temperature they were substantially more stable than the Tet4-Pyr derivative (Figure 6D), for which 50% was degraded after just one day. For **30** and **31,** respectively, the levels of reactive protein remained constant at greater than 95% reactivity after 2 days and 80 and 70% after 8 days of incubation. Reaction kinetics measurements demonstrated a substantial increase in reaction rate with sTCO, achieving *k*₂ values of 84 and 110 × 10⁴ M⁻¹ s⁻¹ (Figure S13). These kinetics values for the di- and tri-fluoro derivatives are, respectively, two and three times faster than the mono-3F-Ph derivative (**27**) and approaching the *k*₂ value of Tet4-Pyr (**21**) (Figure 7). Reactivity in live eukaryotic cells was also verified (Figure 9F and Figure S14C) confirming the utility of the **30** and **31** for fast in-cell protein labeling applications.

## Discussion

To date, the encoded Tet ncAAs have been shown to be the fastest and most quantitative site-specific bioorthogonal conjugation method for labeling proteins *in vitro* and *in vivo*.^[9,48–51]^ Here we provide simple tools to (1) compare the GCE efficiency and fidelity of encoding into proteins, (2) assess reactivity and stability of Tet-proteins and resulting conjugates, (3) determine reaction rates on protein, (4) and validate reactivity in live eukaryotic cells. Using these tools, we characterize the suitabilities of 29 Tet-ncAAs, 20 of which are novel, for bioorthogonal ligations using GCE (Figure 4 and Tables S1-3). These results, for Tet-ncAAs based on the Tet2, Tet3 and Tet4 scaffolds, demonstrate that the tetrazine functional group is remarkably versatile and can be synthetically converted into a diverse pallet of ncAAs many of which can be encoded by previously selected tRNA/RS pairs. While not all derivatives exhibit high permissivity, the incorporation efficiency can be enhanced by selecting new synthetases for the corresponding Tet-ncAAs through a selection process.

The Tet2 and Tet3 tRNA/RS pairs were selected using the Tet2-Me (**3**) and Tet3-Me (**8**) ncAAs,^[9,17]^ whereas the Tet4 tRNA/RS pairs were selected using the Tet4-Ph (**20**) since no functional pairs were obtained when selecting for Tet4-Me (**16**).^[18]^ The initial set of five Tet substituents – (methyl, ethyl, isopropyl, phenyl and pyridyl – surveyed for each scaffold were synthetically accessible in good yields but their encoding efficiency varied dramatically depending on the scaffold. The Tet2 system showed good efficiency for encoding the alkyl substituents but no encoding for the aromatic substituents, presumably because the Tet2 RS active site is too small to accommodate the bulker substituents. Conversely, the Tet4 scaffold was unable to encode the smaller alkyl Tet-ncAA, requiring the phenyl structure for good recognition (Figure 4). Many substituents on the Tet4 phenyl ring are tolerated, but larger more hydrophilic *para* substituents were not. The Tet3 scaffold shows the broadest substrate profile accepting alkyl substituents with good efficiency and aromatic substituents with moderate efficiency, but the addition of the larger and more hydrophilic 4-amino-phe substituent led to low encoding efficiency. This first mapping of the Tet RSs substrate preference shows that Tet-ncAA can be diversly encoded enabling studies on-protein Tet reactivity and stability.

The kind of quantitative labeling to generate a homogeneous modified protein that is desirable for many biochemical studies requires both high efficiency GCE encoding tools and an ncAA labeling system having high reactivity and high stability. There were clear pros and cons for different encoded Tet-ncAAs which were linked to their encoding efficiency and wide reaction rates range spanning over two orders of magnitude from 0.9 × 10^4^ M^-1^s^-1^ for Tet2-Ip (**5**) to 120 × 10^4^ M^-1^s^-1^ for Tet4-Pyr (**21**). The Tet-ncAAs in the “good efficiency” category (Figure 4D) all showed perfect encoding and reactivity as assessed by both mobility shift assay and full protein MS analysis, with Tet4-3,5-diF-Ph (**30**) and Tet4-3,4,5 triF-Ph (**31**) having the fastest on protein reaction rate of 84-110 × 10^4^ M^-1^s^-1^. Those Tet ncAAs encoded with “moderate efficiency” mostly showed 10-20% misencoding due to NCS by MS and unreacted bands in the mobility shift assay at levels 3 – 15% higher than was indicated by MS. The large quantities of protein that can be used in mobility shift assays enable a higher level of sensitivity than standard full protein MS, because low levels of unreactive byproduct can be lost with the background signals from salt peaks and the protein missing the N-terminal methionine (Figure S8). If a Tet-ncAA in the moderate or low efficiency category has reactivity attributes that are desired for quantitative labeling, we expect it will be possible to use de novo selections for that ncAA to generate new high efficiency and fidelity tRNA/RS pairs, as was done with Tet2-ethyl (**4**).^[22]^

Identifying a bioorthogonal chemistry with the right balance of reactivity and stability for quantitative labeling of proteins is not trivial as this requires that (i) all of the encoded Tet-ncAA is sufficiently stable on proteins during translation to be available for reaction, and (ii) that the Tet-protein has a sufficiently high reaction rate as to complete labeling in a reasonable timeframe even at low concentrations of the protein and labeling reagent. Here, we show that it is possible to make multiple Tet-ncAAs with on protein reaction rates of >10^6^ M^-1^s^-1^ that do not have on protein stability issues and that are capable of quantitative labeling. Furthermore, we confirm our earlier findings that when Tet-ncAAs are encoded, their on-protein context increases their reaction rates and the extent of the increase is scaffold dependent. Tet2 and Tet3 scaffolds show a 3-4 fold rate increase when protein encoded, whereas Tet4 scaffolds show a 10-20 fold increase in reactivity with zero cost in stability. Substituent effects on the Tet4 scaffold also make a larger difference than on the Tet3 scaffold. For instance, the addition of a single fluorine to Tet3-Ph (**12**) making Tet3-4F-Ph (**14**) provided just a 10% increase in rate whereas that same addition to the Tet4-Ph (**20**) making Tet4-4F-Ph (**25**) or Tet4-3F-Ph (**27**) provided a 2-fold increase in rate. Furthermore, introducing two and three fluorine on Tet4-3,5-diF-Ph (**30**) and Tet4-3,4,5 triF-Ph (**31**) resulted 4 to 5-fold enhancement in reaction rate.

Since every incremental increase in a bioorthogonal reaction rate advances our ability to monitor short-lived cellular events or to study proteins at lower native-like concentrations, we strive to identify the fastest encodable Tet-ncAA. Some such applications may not need quantitative labeling. These include monitoring membrane receptor recycling^[29,35]^ and monitoring protein conformational changes in cells using double electron-electron spin resonance (DEER) with electron paramagnetic resonance (EPR) labels that have a short lifetime in cells.^[18]^ For such applications, fast-reacting Tet-ncAAs will be perfectly acceptable even if they are not be suitable for achieving quantitative reactions due to exhibiting some degradation in cells such as Tet4-Pyr (**21**) or having low levels of NCS during encoding such as Tet4-3CF_3_-Ph (**29**). In contrast, other labeling applications have a stringent requirement for quantitative labeling and highly efficient protein expression. Examples are producing antibody drug conjugates or quantifying protein quaternary structure on the single molecule level. For these applications, the good efficiency Tet-ncAA encoding systems (Figure 4D) with greater stability and still relatively high reaction rates of 10^4^-10^5^ M^-1^s^-1^ will be preferred.^[52,53]^ Finally, some labeling applications benefit from an ability to control the distance of the label from the protein surface and its flexibility. For those needing longer flexible, linkers for polymer or ligand attachment the Tet3 scaffold is optimal while those needing shorter, less flexible linkers for modern biophysical techniques such as Förster resonance energy transfer (FRET), EPR, NMR and super-resolution microscopy, the Tet4 scaffold will be desired.

Here, we demonstrate that unlike strained alkenes, the tetrazine functionality is a unique bioorthogonal functional group in terms of its remarkable tunability, proving that there exists a diverse palate of Tet-ncAAs that are readily synthesized, can be genetically encoded into proteins, and can react at high rates with strained alkenes. Additionally, we provide a clear road map for furthering explorations of this chemical space, starting with the chemical synthesis of diverse Tet-ncAAs, and progressing through the genetic encoding and evaluation of on-protein reactivity both in vitro and in live eukaryotic cells. In comparing the three Tet scaffolds thus far developed, Tet2, Tet3, and Tet4, we show that the Tet4 scaffold provides an impressive 10 or more-fold reaction rate increase when encoded on proteins, opening the question to what additional Tet scaffolds might provide similar or even higher rate enhancements. Here, we uncovered that the addition of fluorine to the Tet4 scaffold leads to a significant rate enhancement without compromising stability. To generate the smallest, fastest Tet ncAA that maintains high stability for quantitative labeling we used the roadmap to generate ncAAs, Tet4-3,5-diF-Ph (**30**) and Tet4-3,4,5-triF-Ph (**31**), achieving reaction rates at the 10⁶ M⁻¹s⁻¹ level without the rapid degradation observed for Tet4-Pyr (**21**). Others have started to demonstrate that these tRNA/RS scaffolds can be used creatively to encode new ncAA structures with faster reactions or dual functionality,^[10,23]^ here we provide structural guidelines for how these encoding systems can be further exploited. This work significantly enhances the toolbox of bioorthogonal reagents for Tet-ncAA-based, site-specific protein labeling reactions, offering greater flexibility in designing bioconjugate pairs for a wide range of applications.

## Supporting Information

Materials and Methods, Figures S1 to S14, Tables S1-4, ^1^H and ^13^C NMR spectra. This material is available in the Supporting Information.

## Acknowledgements

This research was funded in part by the GCE4All Biomedical Technology Development and Dissemination Center supported by the National Institute of General Medical Science grant RM1-GM144227 as well as National Institutes of Health grant 1R01GM131168-01 and National Science Foundation grant NSF-2054824. Protein mass spectrometry was carried out at the Oregon State University Mass Spectrometry Center on the Waters Ion Mobility ToF Mass Spectrometer supported by NIH grant 1S10RR025628-01.

