## supplemental information for "Tuning Encodable Tetrazine Chemistry for Site-Specific Protein Bioorthogonal Ligations"

**This PDF file includes:**

Materials and Methods
Tables S1–S4
Figures S1 to S14
$^{1}H$ and $^{13}C$ NMR spectra
References 1–7

### Materials and Methods

#### General Synthetic Methods:

All chemicals purchased were utilized without additional purification. Boc-4-cyano-L-phenylalanine, Boc-3-cyano-L-phenylalanine and Boc-β-cyano-L-alanine were purchased from Chem-Impex International. Thin-layer chromatography was performed on silica-coated aluminum plates from Merck pre-coated with $F_{254}$. Flash chromatographic purification was performed using silica gel 60 (230-400 mesh size). $^{1}$H NMR spectra were recorded on a Bruker spectrometer at 400MHz and 700 MHz and $^{13}$C NMR spectra were recorded at 175 MHz. The chemical shifts were shown in ppm and are referenced to the residual non-deuterated solvent peak $CDCl_3$ ($\delta$ =7.26 in $^{1}$H NMR, $\delta$ = 77.23 in $^{13}$C NMR), $CD_3OD$ ($\delta$ =3.31 in $^{1}$H NMR, $\delta$ = 49.0 in $^{13}$C NMR), $d_6$-DMSO ($\delta$ =2.5 in $^{1}$H NMR, $\delta$ = 39.5 in $^{13}$C NMR) as an internal standard. Splitting patterns of protons are designated as follows: s-singlet, d-doublet, t-triplet, q-quartet, m-multiplet, bs- broad singlet, dd-doublet of doublets. ESI-MS data were recorded on Waters, SynaptG2, Q-TOF mass spectrometer for small molecules.

#### Synthetic Procedure of Tet2, Tet3 and Tet4-ncAAs:

#### General Synthetic Procedure for Tet2-derivatives.[1]

Boc-protected Tet2. In a flame-dried, 48 mL heavy-wall reaction tube equipped with a stir bar, Boc-protected 4-cyano phenylalanine (2.75 mmol) was charged with $Ni(OTf)_2$ (1.38 mmol) and corresponding nitriles (27.5 mmol for alkyl nitrile and 19 mmol for aryl nitrile) under an argon atmosphere. Subsequently, anhydrous hydrazine (137.5 mmol) was slowly added to the reaction tube. The reaction mixture was then purged with argon for 10 minutes under constant stirring, sealed immediately then, and heated to 50-55 $^{o}$C for 24 hours (50 $^{o}$C for Tet2-alkyl and 55 $^{o}$C for Tet2-Aryl). Upon completion, the reaction mixture was cooled to room temperature, the sealed reaction vessel was gradually opened, and the mixture was treated with 20 equivalents of 2 M sodium nitrite aqueous solution and 10 mL of water. Ethyl acetate (2x 20 mL) washings were applied to remove the homo-coupling product from the aqueous phase. The aqueous layer was then acidified to pH ~2 with 4 M HCl under ice-cold conditions and extracted with ethyl acetate (3x 30 mL). The combined organic extracts were washed with brine, dried over anhydrous sodium sulfate, and concentrated under reduced pressure. Flash column chromatography on silica gel (eluted with 30-35% ethyl acetate containing 1% acetic acid in hexanes) afforded Boc-protected Tet2- amino acids as a pinkish-red gummy material with a yield ranging from 30 to 80%.

Chloride salt of Tet2. In a dry round-bottom flask, Boc-protected Tet2 (2.23 mmol) in 5 mL ethyl acetate was charged with 3 mL 1,4-dioxane saturated with HCl gas under an argon atmosphere. The reaction proceeded at room temperature with continuous stirring until the starting materials were fully consumed, as determined by thin-layer chromatography (typically 3 to 4 h). Then it was concentrated under reduced pressure, re-dissolved in ethyl acetate (2x 10 mL), and similarly concentrated to remove excess HCl gas, yielding Tet2 as a pink solid material with quantitative yield.

**General Synthetic Procedure for Tet3-derivatives.**[2]

Following the above procedure for making Tet2, Boc-protected 3-cyano phenylalanine was used to synthesize and purify Tet3-derivatives.

**General Synthetic Procedure for Tet4-derivatives.**[3]

*Boc-protected Tet4*. In a flame-dried, 15 mL heavy-walled reaction tube, Boc-protected β-cyano L-alanine (0.560 mmol), Nickel(II) trifluoromethanesulfonate ($Ni(OTf)_2$) (0.28 mmol), and corresponding nitrile (5.6 mmol for alkyl nitrile and 3.9 mmol for aryl nitrile) were taken under an argon atmosphere. Then, anhydrous hydrazine (28 mmol) and 0.3 mL of ethanol (EtOH) were added to the reaction mixture. After purging with argon for 10 min and immediate sealing of the tube, the reaction mixture was heated to 37- 42 °C for 30 h (37 $^{o}$C for Tet4-alkyl and 42 $^{o}$C for Tet4-Aryl). The reaction mixture was cooled to room temperature, the mixture was slowly opened, and 10 equivalents of 2 M sodium nitrite aqueous solution and 10 mL of water were added. Subsequently, the reaction mixture was washed with ethyl acetate to eliminate the homocoupled product. The aqueous phase was then acidified with 4 M HCl (pH ~2) under ice-cold conditions and extracted with ethyl acetate (3 times). The combined organic layer was dried with anhydrous sodium sulfate and concentrated under reduced pressure. Flash column chromatography on silica gel (eluted with 15-35 % ethyl acetate in hexanes with 1% acetic acid) afforded Boc-protected Tet4- amino acids.

Trifluoroacetate salt of Tet4-alkyl (Me, Et, Isp, Bu) derivatives. In a dry round-bottom flask, Boc-protected Tet4-alkyl derivatives (0.5 mmol) were dissolved in a 1:1 mixture of dry dichloromethane and trifluoroacetic acid (TFA) by volume (total volume 1.5-2 mL) under argon. The reaction mixture was stirred at room temperature for 1 h. After completion of the reaction, it was concentrated under reduced pressure and dissolved in dichloromethane to remove TFA. This process was repeated twice before drying completely under high vacuum, resulting in a pinkish red TFA salt of Tet4, obtained in quantitative yield.

Chloride salt of Tet4-aryl derivatives. Following the Boc-deprotection procedure for making Tet2, Boc-Tet4-aryl derivatives were converted to chloride salt in quantitative yields.

**Characterization of Tet2, Tet3, and Tet4-ncAAs.**

**Hydrochloride salt of (S)-2-amino-3-(4-(6-methyl-1,2,4,5-tetrazin-3-yl)phenyl) propanoic acid**:[x]

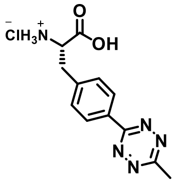

**Tet2-Me**: $^{1}H$ NMR (400MHz, $CD_3OD$) $\delta$ 8.56 (d, 2H), 7.60 (d, 2H), 4.39 (t, 1H), 3.48- 3.33 (dd, 2H), 3.1 (s, 3H). $^{13}C$ NMR (175MHz, $CD_3OD$) $\delta$ 171.2, 169.1, 165.3, 140.7, 133.3, 131.7, 129.6, 55.1, 37.4, 21.3. ESI-MS calculated for $C_{12}H_{14}N_5O_2$ ([M + H]$^+$) 260.114, found 260.10.

**Hydrochloride salt of (S)-2-amino-3-(4-(6-ethyl-1,2,4,5-tetrazin-3-yl)phenyl) propanoic acid**:[x]

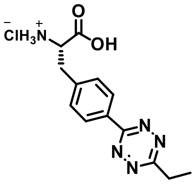

**Tet2-Et**: $^{1}H$ NMR (700MHz, $d_6$-DMSO) $\delta$ 8.61 (bs, 1H), 8.42 (d, 2H), 7.59 (d, 2H), 4.26 (bs, 1H), 3.32 (q, 2H), 3.30-3.26 (dd, 2H), 1.46 (t, 3H). $^{13}C$ NMR (175MHz, $d_6$-DMSO) $\delta$ 170.8, 170.7, 163.8, 140.3, 131.3, 131.1, 128.1, 53.3, 36.1, 28.1, 12.2. ESI-MS calcd for $C_{13}H_{16}N_5O_2$ ([M + H]$^+$) 274.1299, found 274.12.

**Hydrochloride salt of (S)-2-amino-3-(4-(6-isopropyl-1,2,4,5-tetrazin-3-yl)phenyl) propanoic acid**:

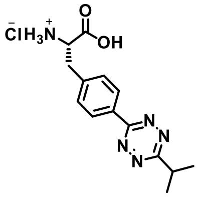

**Tet2-Ip**: $^{1}H$ NMR (700MHz, $d_6$-DMSO) $\delta$ 8.53 (bs, 1H), 8.43 (d, 2H), 7.59 (d, 2H), 4.26 (t, 1H), 3.60 (sept, 1H), 3.27 (d, 2H), 1.48 (d, 6H). $^{13}C$ NMR (175MHz, $d_6$-DMSO) $\delta$ 172.6, 170.1, 163.2, 139.6, 130.6, 130.3, 127.4, 52.6, 35.4, 33.3, 20.7. ESI-MS calcd for $C_{14}H_{18}N_5O_2$ ([M + H]$^+$) 288.1455, found 288.14.

**Hydrochloride salt of (S)-2-amino-3-(4-(6-phenyl-1,2,4,5-tetrazin-3-yl)phenyl) propanoic acid**:

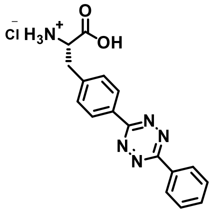

**Tet2-Ph**: $^{1}$H NMR (700MHz, $d_6$-DMSO) $\delta$ 8.55 (dd, 2H), 8.50 (d, 2H), 7.75-7.70 (m, 3H), 7.61 (d, 2H), 4.30 (t, 1H), 3.29 (d, 2H). $^{13}$C NMR (175MHz, $d_6$-DMSO) δ 170.3, 163.2, 163.2, 140.1, 132.7, 131.9, 130.9, 130.7, 129.5, 127.8, 127.5, 52.9, 35.6. ESI-MS calcd for $C_{17}H_{16}N_5O_2$ ([M + H]$^+$) 322.1299, found 322.10.

**Hydrochloride salt of (S)-2-amino-3-(4-(6-(pyridin-2-yl)-1,2,4,5-tetrazin-3-yl)phenyl)-propanoic acid*:***

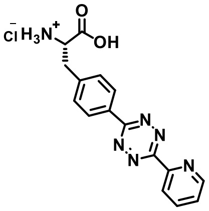

**Tet2-Py**: $^{1}$H NMR (700MHz, $d_6$-DMSO) $\delta$ 8.94 (d, 1H), 8.60 (d, 1H), 8.54 (d, 2H), 8.46 (bs, 1H), 8.17 (td, 1H), ), 7.74 (dd, 1H), 7.63 (d, 2H), 7.52 (d, 1H), 4.31 (t, 1H), 3.29 (d, 2H). $^{13}$C NMR (175MHz, $d_6$-DMSO) δ 170.3, 163.4, 163.1, 150.5, 150.1, 137.9, 130.7, 130.5, 128.1, 127.7, 126.6, 124.1, 52.8, 35.7. ESI-MS calcd for $C_{16}H_{15}N_6O_2$ ([M + H]$^+$) 323.1251, found 323.12.

**Hydrochloride salt of Tet3-Me, Tet3-Et, Tet3-Isp, Tet3-Bu:** See previous published paper Jang et al.[2]

**Hydrochloride salt of (S)-2-amino-3-(3-(6-phenyl-1,2,4,5-tetrazin-3-yl)phenyl)propanoic acid :**

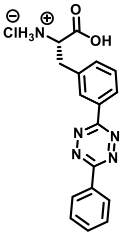

**Tet3-Ph**: $^{1}$H NMR (700MHz, $d_6$-DMSO) $\delta$ 8.55 (d, 2H), 8.49 (s, 1H), 8.46 (d, 1H), 7.73-7.68 (m, 3H), 7.67 (t, 1H), 7.64 (d, 1H), 4.32 (bs, 1H), 3.32 (dd, 2H). $^{13}$C NMR (175MHz, $d_6$-DMSO) $\delta$ 170.3, 163.5, 163.4, 136.5, 134.1, 132.8, 132.2, 131.9, 129.9, 129.6, 128.8, 127.7, 126.7, 53.1, 35.7. ESI-MS calcd for $C_{17}H_{16}N_5O_2$ ([M + H]$^+$) 322.1299, found 322.13.

**Hydrochloride salt of (S)-2-amino-3-(3-(6-(pyridin-2-yl)-1,2,4,5-tetrazin-3-yl)phenyl)propanoic acid:**

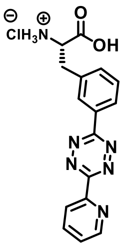

**Tet3-Py**: $^{1}H$ NMR (700MHz, $CD_3OD$) $\delta$ 9.20 (d, 1H), 8.82 (s, 1H), 8.73-8.70 (m, 2H), 8.29 (s, 1H), 7.75 (d, 2H), 7.61 (dd, 1H), 4.45 (t, 1H), 3.35 (d, 1H). 3.41 (d, 1H). $^{13}C$ NMR (175MHz, $CD_3OD$) δ 171.2, 166.3, 161.5, 147.2, 146.7, 137.7, 136.1, 133.6, 131.6, 130.9, 129.4, 127.7, 127.2, 55.1, 37.4. ESI-MS calcd for $C_{16}H_{15}N_6O_2$ ([M + H]$^+$) 323.1251, found 323.12.

**Hydrochloride salt of (S)-2-amino-3-(3-(6-(4-fluorophenyl)-1,2,4,5-tetrazin-3-yl)phenyl)propanoic acid:**

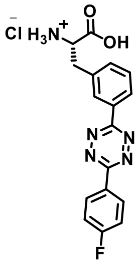

**Tet3-4F-Ph**: $^{1}H$ NMR (700MHz, $CD_3OD$) $\delta$ 8.70 (d, 2H), 8.60 (td, 2H), 7.69 (t, 1H), 7.64 (d, 1H), 7.43 (t, 2H), 4.40 (q, 1H), 3.48 (dd, 1H). 3.37 (dd, 1H). $^{13}C$ NMR (175MHz, $CD_3OD$) δ 171.3, 166.5, 165.2, 164.8, 137.3, 134.9, 131.6, 130.6, 128.5, 117.7, 117.6, 55.1, 37.3. ESI-MS calcd for $C_{17}H_{15}FN_5O_2$ ([M + H]$^+$) 340.1204, found 340.11.

**Hydrochloride salt of (S)-2-amino-3-(3-(6-(4-aminophenyl)-1,2,4,5-tetrazin-3-yl)phenyl)propanoic acid:**

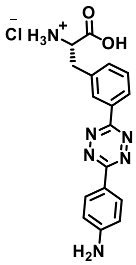

**Tet3-4NH$_2$-Ph**: $^{1}H$ NMR (700MHz, $CD_3OD$) $\delta$ 8.64 (dd, 3H), 8.57 (d, 1H), 7.71-7.63 (m, 2H), 7.42 (d, 1H), 7.35 (d, 1H), 4.41 (t, 1H), 3.50 (dd, 1H). 3.39 (dd, 1H). $^{13}C$ NMR (175MHz, $d_6$-

DMSO) δ 170.3, 163.3, 162.1, 142.2, 136.3, 132.4, 132.1, 129.7, 129.4, 128.7, 128.1, 125.9, 114.8, 53.1, 35.7. ESI-MS calcd for $C_{17}H_{17}N_6O_2$ ([M + H]$^+$) 337.1408, found 337.14.

**Trifluoro acetate salt of (S)-2-amino-3-(6-methyl-1,2,4,5-tetrazin-3-yl)propanoic acid:**[3]

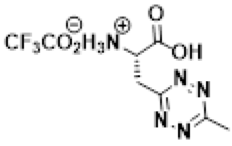

**Tet4-Me**: $^1$H NMR (700MHz, $CD_3OD$) δ 4.48 (bs, 1H), 3.98 (dd, 1H), 3.83 (dd, 1H), 3.04 (s, 3H). $^{13}$C NMR (175MHz, $CD_3OD$) δ 169.7, 167.7, 163.5, 51.9, 36.7, 21.3. ESI-MS calculated for $C_6H_{10}N_5O_2$ ([M + H]$^+$) 184.0829, found 184.08.

**(S)-2-((tert-butoxycarbonyl)amino)-3-(6-ethyl-1,2,4,5-tetrazin-3-yl)propanoic acid:**

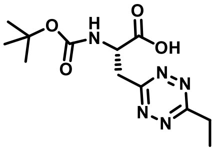

**Boc-Tet4-Et**: $^1$H NMR (400MHz, $CDCl_3$) δ 5.71 (d, 1H), 4.91 (bs, 1H), 3.94 (dd, 1H), 3.79 (dd, 1H), 3.35 (q, 2H), 1.51 (t, 3H), 1.36 (s, 9H). $^{13}$C NMR (175MHz, $CDCl_3$) δ 174.6, 171.2, 166.7, 155.4, 80.5, 51.8, 37.3, 28.2, 20.8, 12.2. ESI-MS calculated for Boc-deprotected Tet4-Et, $C_7H_{12}N_5O_2$ ([M + H]$^+$) 198.0986, found 198.09.

**(S)-2-((tert-butoxycarbonyl)amino)-3-(6-isopropyl-1,2,4,5-tetrazin-3-yl)propanoic acid:**

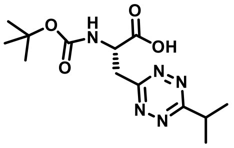

**Boc-Tet4-Isp**: $^1$H NMR (400MHz, $CDCl_3$) δ 5.67 (d, 1H), 4.97 (bs, 1H), 3.95 (dd, 1H), 3.81 (dd, 1H), 3.65 (sept, 1H), 1.53 (d, 6H), 1.39 (s, 9H). $^{13}$C NMR (175MHz, $CDCl_3$) δ 174.9, 174.1, 166.6, 155.4, 80.6, 51.6, 37.2, 34.2, 28.2, 21.2. ESI-MS calculated for Boc-deprotected Tet4-Isp, $C_8H_{14}N_5O_2$ ([M + H]$^+$) 212.1142, found 212.11.

**(*S*)-2-((*tert*-butoxycarbonyl)amino)-3-(6-butyl-1,2,4,5-tetrazin-3-yl)propanoic acid:**

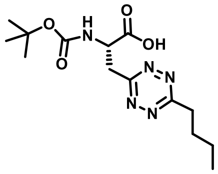

**Boc-Tet4-Bu**: $^1$H NMR (400MHz, $CDCl_3$) δ 5.60 (d, 1H), 4.91 (q, 1H), 3.90 (dd, 1H), 3.78 (dd, 1H), 3.28 (t, 2H), 1.90 (quin, 2H), 1.44 (sext, 2H), 1.35 (s, 9H), 0.95 (t, 3H). $^{13}$C NMR (175MHz, $CDCl_3$) δ 174.6, 170.8, 166.7, 155.6, 80.8, 51.8, 37.3, 34.6, 30.4, 28.3, 22.4, 13.9.

**Trifluoro acetate salt of (S)-2-amino-3-(6-butyl-1,2,4,5-tetrazin-3-yl)propanoic acid:**

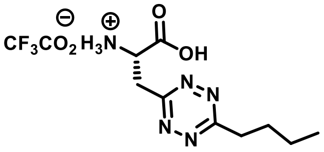

**Tet4-Bu**: $^1$H NMR (400MHz, $CD_3OD$) δ 4.46 (t, 1H), 4.40 (dd, 1H), 3.89 (dd, 1H), 3.33 (t, 2H), 1.95 (quin, 2H), 1.49 (sext, 2H), 1.03 (t, 3H). $^{13}$C NMR (175MHz, $CD_3OD$) δ 172.3, 167.5, 163.1, 52.8, 36.3, 35.4, 31.3, 23.3, 14.1. ESI-MS calculated for $C_9H_{16}N_5O_2$ ([M + H]$^+$) 226.1299, found 226.12.

**Hydrochloride salt of (S)-2-amino-3-(6-phenyl-1,2,4,5-tetrazin-3-yl)propanoic acid:**[3]

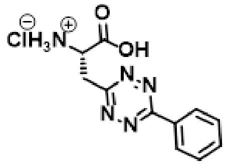

**Tet4-Ph**: $^1$H NMR (700MHz, $CD_3OD$) δ 8.59 (d, 2H), 7.69 (t, 1H), 7.65 (t, 2H), 4.81 (dd, 1H), 4.03 (ddd, 2H). $^{13}$C NMR (175MHz, $CD_3OD$) δ 170.6, 166.9, 166.3, 134.2, 133.4, 130.6, 129.2, 52.2, 36.22. ESI-MS calculated for $C_{11}H_{12}N_5O_2$ ([M + H]$^+$) 246.0986, found 246.09.

**Hydrochloride salt of (S)-2-amino-3-(6-(pyridin-2-yl)-1,2,4,5-tetrazin-3-yl) propanoic acid:**[3]

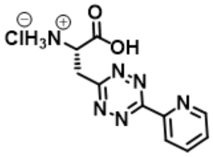

**Tet4-Py (21)**: $^1$H NMR (700MHz, $CD_3OD$) $\delta$ 9.1 (d, 1H), 9.06 (d, 1H), 8.71 (t, 1H), 8.21 (t, 1H), 4.23(dd, 1H), 4.13(dd, 1H). $^{13}$C NMR (175MHz, $CD_3OD$) $\delta$ 170.3, 169.1, 162.5, 147.6, 147.1, 146.3, 130.7, 127.2, 52.1, 36.5. ESI-MS calculated for $C_{10}H_{11}N_6O_2$ ([M + H]$^+$) 247.0938, found 247.09.

**Hydrochloride salt of (S)-2-amino-3-(6-(p-tolyl)-1,2,4,5-tetrazin-3-yl)propanoic acid:**

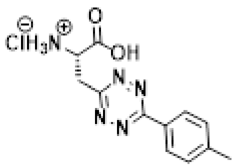

**Tet4-4Me-Ph**: $^1$H NMR (700MHz, $CD_3OD$) δ 8.48 (d, 2H), 7.47 (d, 2H), 4.79 (q, 1H), 4.05 (dd, 1H), 3.96 (dd, 1H), 2.48 (s, 3H). $^{13}$C NMR (175MHz, $CD_3OD$) δ 170.6, 166.9, 166.3, 145.3, 131.3, 130.5, 129.2, 52.3, 36.2, 21.8. ESI-MS calculated for $C_{12}H_{14}N_5O_2$ ([M + H]$^+$) 260.1142, found 260.11.

**Hydrochloride salt of (S)-2-amino-3-(6-(m-tolyl)-1,2,4,5-tetrazin-3-yl)propanoic acid:**

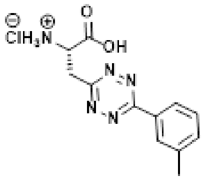

**Tet4-3Me-Ph**: $^1$H NMR (700MHz, $CD_3OD$) δ 8.44 (s, 1H), 8.40 (d, 1H), 7.54 (m, 2H), 4.83 (t, 1H), 4.09 (dd, 1H), 3.99 (dd, 1H), 2.51 (s, 3H). $^{13}$C NMR (175MHz, $CD_3OD$) δ 170.6, 167.1, 166.3, 140.7, 134.8, 133.2, 130.5, 129.6, 126.4, 52.2, 36.1, 21.6. ESI-MS calculated for $C_{12}H_{14}N_5O_2$ ([M + H]$^+$) 260.1142, found 260.11.

**Hydrochloride salt of (S)-2-amino-3-(6-(4-fluorophenyl)-1,2,4,5-tetrazin-3-yl)propanoic acid**:

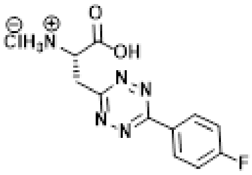

**Tet4-4F-Ph**: $^1$H NMR (700MHz, $CD_3OD$) δ 8.68 (dd, 1H), 8.62 (dd, 1H), 7.68 (dt, 1H), 7.42 (t, 1H), 4.82 (dd, 1H), 4.01 (dd, 1H), 3.99 (m, 1H). $^{13}$C NMR (175MHz, $CD_3OD$) δ 170.6, 167.2, 166.3, 165.5, 131.1, 129.2, 117.7, 52.3, 36.2. ESI-MS calculated for $C_{11}H_{11}FN_5O_2$ ([M + H]$^+$) 264.0891, found 264.08.

**Hydrochloride salt of (S)-2-amino-3-(6-(3-fluorophenyl)-1,2,4,5-tetrazin-3-yl)propanoic acid**:

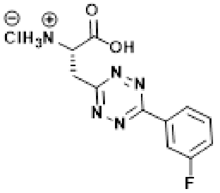

**Tet4-3F-Ph**: $^1$H NMR (700MHz, $CD_3OD$) δ 8.47 (d, 1H), 8.33 (d, 1H), 7.71 (q, 1H), 7.47 (td, 1H), 4.83 (m, 1H), 4.11 (dd, 1H), 4.03 (dd, 1H). $^{13}$C NMR (175MHz, $CD_3OD$) δ 170.5, 167.5, 165.4, 164.1 135.6, 132.6, 125.1, 120.9, 115.5, 52.1, 36.2. ESI-MS calculated for $C_{11}H_{11}FN_5O_2$ ([M + H]$^+$) 264.0891, found 264.08.

**Hydrochloride salt of (S)-3-(6-(3-acetylphenyl)-1,2,4,5-tetrazin-3-yl)-2-aminopropanoic acid:**

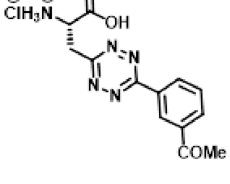

**Tet4-3COMe-Ph**: $^1$H NMR (700MHz, $CD_3OD$) δ 9.21 (s, 1H), 8.84 (d, 1H), 8.35 (d, 1H), 7.84 (t, 1H), 4.84 (t, 1H), 4.12 (dd, 1H), 4.02 (dd, 1H), 2.74 (s, 3H). $^{13}$C NMR (175MHz, $CD_3OD$) δ 199.4,

170.5, 167.5, 165.7, 139.5, 133.9, 133.7, 133.4, 131.2, 128.8, 52.2, 36.2, 26.9. ESI-MS calculated for $C_{13}H_{14}N_5O_3$ ([M + H]$^+$) 288.1091, found 288.10.

**Hydrochloride salt of (S)-2-amino-3-(6-(3-(trifluoromethyl)phenyl)-1,2,4,5-tetrazin-3-yl)propanoic acid:**

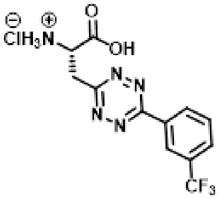

**Tet4-3CF$_3$-Ph**: $^1$H NMR (700MHz, $CD_3OD$) $\delta$ 9.89 (bs, 2H), 8.06 (d, 1H), 7.92 (t, 1H), 4.43 (m, 1H), 4.13 (dd, 1H), 4.03 (dd, 1H). $^{13}$C NMR (175MHz, $CD_3OD$) δ 170.7, 167.7, 165.2, 134.5, 132.6, 131.8, 130.5, 126.3, 125.7, 125.6, 52.3, 36.3. ESI-MS calculated for $C_{12}H_{11}F_3N_5O_2$ ([M + H]$^+$) 314.0859, found 314.09.

**Hydrochloride salt of (S)-2-amino-3-(6-(4-aminophenyl)-1,2,4,5-tetrazin-3-yl)propanoic acid:**

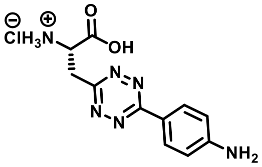

**Tet4-4NH$_2$-Ph**: $^1$H NMR (700MHz, $CD_3OD$) δ 8.65 (d, 2H), 7.37 (d, 2H), 4.81 (q, 1H), 4.08 (dd, 1H), 3.98 (dd, 1H). $^{13}$C NMR (175MHz, $CD_3OD$) δ 170.6, 167.1, 165.7, 146.6, 131.1, 130.7, 121.2, 52.3, 36.2. ESI-MS calculated for $C_{11}H_{13}N_6O_2$ ([M - H]$^-$) 259.0946, found 259.09.

**Hydrochloride salt of (S)-2-amino-3-(6-(4-hydroxyphenyl)-1,2,4,5-tetrazin-3-yl)propanoic acid:**

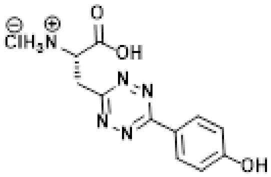

**Tet4-4OH-Ph**: $^1$H NMR (700MHz, $CD_3OD$) δ 8.47 (d, 2H), 7.02 (d, 2H), 4.78 (q, 1H), 4.03 (dd, 1H), 3.93 (dd, 1H). $^{13}$C NMR (175MHz, $CD_3OD$) δ 170.5, 166.2, 166.1, 163.8, 131.2, 124.1, 117.4, 52.2, 36.1. ESI-MS calculated for $C_{12}H_{16}N_5O_3$ ([M + H]$^+$) 262.0940, found 262.09.

**Scheme 1: Synthesis of Tet4-3,5-difluorophenyl derivative**

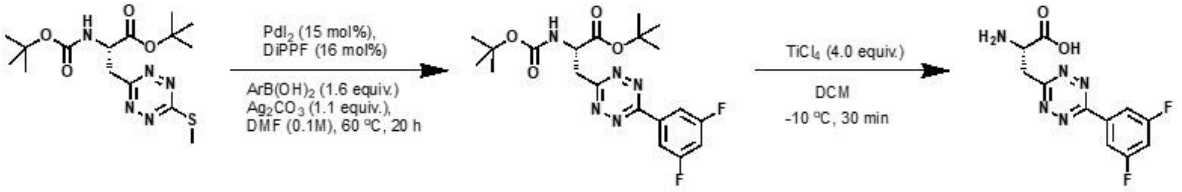

***tert*-butyl (S)-2-((*tert*-butoxycarbonyl)amino)-3-(6-(3,5-difluorophenyl)-1,2,4,5-tetrazin-3-yl)propanoate (Protected Tet4-3,5-diFphenyl)**: The compound was synthesized according to the literature procedure with some modifications.[4,5] In a two-neck round bottom flask equipped with a magnetic stir bar anhydrous DMF (0.1 M, w.r.t. tetrazine starting material) was added and degassed by purging with argon, followed by the addition of 1,1' bis(diisopropylphosphino)-ferrocene (DiPPF) (11 mol%) and Palladium(II) iodide ($PdI_2$) (10 mol%) under an inert atmosphere. The reaction mixture was heated at 60 °C for 1 h. To this reaction mixture was added *tert*-butyl (S)-2-((*tert*-butoxycarbonyl)amino)-3-(6-(methylthio)-1,2,4,5-tetrazin-3-yl)propanoate (0.25 g, 0.67 mmol, 1.0 equiv.), (3,5-difluorophenyl)boronic acid (0.17 g, 1.07 mmol, 1.6 equiv.) followed by silver carbonate (0.203 g, 0.74 mmol, 1.1 equiv.) and continued stirring at 60 °C till 20 h. Upon completion, the reaction mixture was filtered through a celite pad, and the filtrate was diluted with water and extracted with ethyl acetate (20 mL x 3). The organic fraction was collected, the solvent was evaporated under vacuum and the residue was purified by column chromatography using 5 -10 % ethyl acetate in hexane as an eluent as a bright red solid Yield: 0.21 g, 71%.

**N-Boc-C-$^t$Bu-Tet4-3,5-diF-Ph**: $^1$H NMR (700 MHz, $CDCl_3$) δ 8.19 – 8.10 (m, 2H), 7.13 – 7.03 (m, 1H), 5.47 (d, *J* = 7.1 Hz, 1H), 4.83 – 4.73 (m, *J* = 12.5, 7.0 Hz, 1H), 3.91 (dd, *J* = 20.2, 10.2 Hz, 1H), 3.76 (dd, *J* = 14.5, 7.2 Hz, 1H), 1.44 (s, 9H), 1.34 (s, 9H). $^{13}$C NMR (176 MHz, $CDCl_3$) δ 169.69, 167.68, 164.47, 164.40, 163.05, 162.98, 155.21, 135.10, 111.24, 111.21, 111.11, 111.08, 108.31, 108.17, 108.03, 99.55, 99.52, 99.39, 96.25, 83.30, 80.31, 52.99, 38.47, 28.35, 28.05. (splitting in $^{13}$C resonances observed due to $^{13}$C-$^{19}$F coupling). ESI-MS Calculated for $C_{20}H_{25}F_2N_5O_4$ $[M+Na]^+$: 460.1767; Found: 460.17. (A major fragment ion at 282.08 was observed due to loss of N-Boc and *tert*-butyl ester group).

**(S)-2-amino-3-(6-(3,5-difluorophenyl)-1,2,4,5-tetrazin-3-yl)propanoic acid (Tet4.0-3,5-diF phenyl.HCl)**: In a round bottom flask equipped with a magnetic stir bar was charged with *tert*-butyl 2-((*tert*-butoxycarbonyl)amino)-3-(6-(3,5-difluorophenyl)-1,2,4,5-tetrazin-3-yl)propanoate (0.147 g, 0.34 mmol, 1.0 equiv.) was dissolved in anhydrous DCM (0.4 M). The reaction flask was sealed and purged with nitrogen. A solution of $TiCl_4$ (0.128 g, 0.67 mmol, 2.0 equiv.) in 1 M DCM was added to the reaction mixture dropwise at -20 °C. The reaction mixture was allowed to stir at -10 to -15 °C for 1 – 2 h. Upon completion, the reaction mixture was filtered through a

Whatman filter paper. The residue was washed with excess DCM followed by EtOAc. The residue was dissolved in methanol and the solvent was evaporated to yield the product as a pink solid. Yield: 0.1 g, 94 %.

**Tet4-3,5-diF-Ph**: $^{1}$H NMR (700 MHz, MeOD) δ 8.21 – 8.10 (m, 2H), 7.38 – 7.27 (m, 1H), 4.82 (s, 1H), 4.31 – 4.02 (m, 2H). $^{13}$C NMR (176 MHz, MeOD) δ 169.42, 167.29, 165.72, 165.65, 164.38, 164.31, 164.23, 136.68, 111.94, 111.90, 111.81, 111.78, 111.73, 108.99, 108.84, 108.80, 108.70, 54.03, 51.93, 35.82. (splitting in $^{13}$C resonances observed due to $^{13}$C-$^{19}$F coupling). ESI-MS: Calc. for $C_{11}H_9F_2N_5O_2$ [M+H]$^+$: 282.0797; Found: 282.07.

**Scheme 2: Synthesis of Tet4-3,4,5-triFluorophenyl derivative**

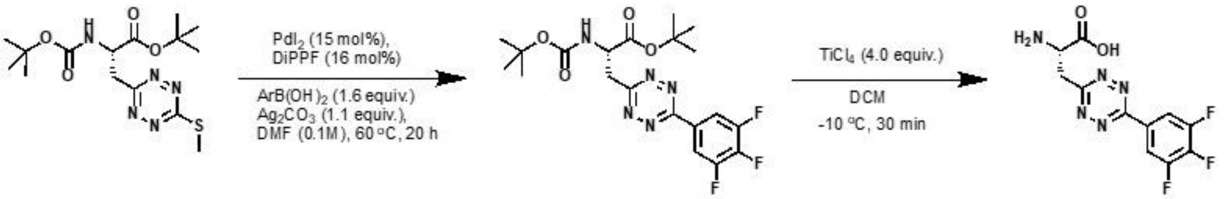

***tert*-butyl (S)-2-((*tert*-butoxycarbonyl)amino)-3-(6-(3,4,5-trifluorophenyl)-1,2,4,5-tetrazin-3-yl)propanoate (protected Tet4-3,4,5triFphenyl)**: Following the synthetic procedure of Tet4-3,5-diF-Ph and using (3,4,5-difluorophenyl)boronic acid, the title compound was obtained as a bright red solid (Yield: 81%).

**N-Boc-C-$^t$Bu-Tet4-3,4,5-triF-Ph**: $^{1}$H NMR (700 MHz, $CDCl_3$) δ 8.34 – 8.23 (m, 2H), 5.46 (d, *J* = 7.2 Hz, 1H), 4.75 (m, 1H), 3.91 (m, 1H), 3.74 (m, 1H), 1.44 (s, 9H), 1.33 (s, 9H). $^{13}$C NMR (176 MHz, $CDCl_3$) δ 169.65, 167.57, 162.30, 155.21, 152.79, 152.75, 151.38, 151.32, 112.69, 112.66, 112.58, 112.56, 83.36, 80.33, 53.03, 38.51, 28.33, 28.05.(splitting in $^{13}$C resonances observed due to $^{13}$C-$^{19}$F coupling). ESI-MS Calculated for $C_{20}H_{24}F_3N_5O_4$ [M+Na]$^+$: 478.1673; Found: 478.16, (a major fragment ion at 300.0698 was observed due to loss of N-Boc and *tert*-butyl ester group).

**(S)-2-amino-3-(6-(3,4,5-trifluorophenyl)-1,2,4,5-tetrazin-3-yl)propanoic acid (Tet4.0-3,4,5-TriF phenyl)**: N-Boc-C-$^t$Bu was deprotected following the similar procedure of N, C-terminal deprotection of Tet4-3,5-diF-Ph, yielding the product as a pink solid (Yield: 90%).

**Tet4-3,4,5-triF-Ph**: $^{1}$H NMR (700 MHz, MeOD) δ 8.43 – 8.30 (m, 2H), 4.81 (s, 1H), 4.14 – 3.98 (m, 2H). $^{13}$C NMR (176 MHz, MeOD) δ 169.47, 167.20, 163.92, 153.83, 152.41, 113.70, 113.67, 113.56, 54.07, 51.98, 35.85. ESI-MS Calculated for $C_{11}H_8F_3N_5O_2$ [M+H]$^+$: 300.0703; Found: 300.06.

**Free Tet-ncAAs rate constant measurement.** Following the published protocol Jang et al.[2]

**Toxicity study of Tet-ncAAs**. A stable and nontoxic Tet-ncAAs is essential for cell-based Tet-protein expression. In our efforts to evolve tRNA synthetases (RS), we typically screened for viability using 1 mM concentrations. Thus, to assess the cell viability of Tet-ncAAs in *E. coli*, we monitored cell growth at concentrations of 0.25 mM, 0.5 mM, 0.75 mM, and 1 mM. Fresh and saturated DH10B *E. coli* cell stocks were inoculated into 5 mL auto-inducing media (AIM) with and without Tet-ncAAs, allowing growth for 40 h at 37°C. Optical density ($OD_{600}$) measurements taken at 24 and 40 h revealed that Tet2-alkyl and Tet3-alkyl derivatives at 1 mM exhibited minimal effects on cell growth, while pyridyl substituents significantly reduced the growth. On the other hand, Tet4-Ph impeded the cell growth and exhibited toxicity at 1 mM, whereas Tet4-alkyl substituents showed favorable behavior (Figure S1). Additionally, we observed that the cells tolerated both Tet3-Py and Tet4-Ph in their 1,4 dihydro form at 1 mM (reduced form of tetrazine), indicating lower toxicity compared to oxidized tetrazines (Figures S2 and S3).

Thus, to mitigate Tet-ncAAs' toxicity, reducing the concentration of Tet-ncAA can be achieved by enhancing the efficiency of the GCE machinery. At a lower concentration of 0.25 mM, Tet4-Ph proved capable of yielding substantial amounts of Tet-proteins. Additionally, we observed that cell toxicity was partially circumvented by supplementing Tet-ncAA into the media after 4 to 6 h of cell inoculation, when the cells reached a healthy growth phase ($OD_{600}$ ~ 0.5 to 0.7) (Figure S4). This approach significantly reduces Tet-ncAA toxicity for Tet-protein expression in *E. coli* cells.

**Permissivity screen for Tet2, Tet3, and Tet4-ncAAs with selected synthetases.** To determine whether our previously chosen amino acyl tRNA-synthetases (aaRS/tRNA pairs) for Tet2, Tet3 and Tet4 could accommodate a structurally similar set of Tet-ncAA derivatives, we assessed the expressions of sfGFP150TAG with and without 0.5 mM Tet-ncAAs in 25 mL auto-inducing media. This assessment was carried out using RSs D12-Tet2, R2-84-Tet3, and Tet4-1 (E1), Tet4-2 (D4) for each set respectively. Cultures were grown for 30-32 h at 37 °C and 250 rpm in the presence of respective antibiotics. The sfGFP expression of the of the culture was measured by fluorescence using a Turner Biosystems Picofluor fluorimeter diluting 100 µL cell culture in 1.9 mL water. (Figure 4).

**Expression and purification of sfGFP150-TAG-Tet-ncAAs.** Following the expression conditions outlined earlier for efficiency and fidelity measurements, sfGFP150-Tet was expressed in 50 mL AIM. After 32 h, all cells were harvested by centrifugation at 5000 rcf for 10 minutes. The media was then removed, and the cell pellets were stored at -80 °C. Cells were resuspended in wash buffer (NaCl 300 mM, $NaH_2PO_4$ 15.5 mM, $Na_2HPO_4$ 34.5 mM, imidazole 5 mM, pH 7.1). Cells were lysed using a Microfluidics M-110P microfluidizer (18,000 psi) and the lysate was collected in wash buffer. Subsequently, the lysate was clarified by centrifugation (at 21000 rcf for 30 minutes), and TALON resin (300 µL bed volume) was added to the clarified supernatant. The lysate was incubated with the resin for 1-2 h, gently rocking at 4°C. Afterward, the resin and lysate were applied to a column, and the flow-through was discarded. The resin was washed 5 times with 10 mL wash buffer. Protein was eluted with 250 µL elution buffer (NaCl 300 mM, $NaH_2PO_4$ 15.5 mM, $Na_2HPO_4$ 34.5 mM, imidazole 250 mM, pH 7.0). Protein concentration was determined by measuring absorbance at 280 nm. Protein purity was assessed using SDS-PAGE.

**Mobility Shift Assay.** To assess the purity and extent of Tet-reactivity, purified sfGFP-Wt and sfGFP150-Tet-ncAA variants (50 µM) were exposed to excess sTCO-PEG5k (250 µM) and allowed to react for 10 minutes in PBS at room temperature. Then, the protein was denatured through the addition of Laemmli buffer and heated at 95 °C for 10 minutes. Samples were then analyzed using a 12% SDS-PAGE gel (Figures 5 and S5-7).

**Mass spectra of GFP-Tet-ncAA.** Purified sfGFP-TAG150-Tet-ncAA was diluted to 50 µM and desalted using Zeba™ spin desalting column and analyzed using an FT LTQ mass spectrometer at the Oregon State University mass spectrometry facility. Waters SYNAPT G2 HDMS with a Waters Acquity I class UPLC mass spectrometer was used to verify the reaction of purified sfGFP-TAG150-Tet4 with sTCO. Samples were run 45 min gradient with $H_2O$:ACN: 0.1% formic acid using a Thermo Scientific- MAbPac™RP column. 2.1x100 mm and a 0.2ml/min flow rate. Spectra were deconvoluted using the Maximum Entropy deconvolution algorithm (MaxEnt3) in Waters MassLynx software.

**Measuring reaction rates of Tet-ncAA with sTCO-OH on protein.** The fluorescence of sfGFP undergoes quenching by 4-6 times when Tet-ncAA is encoded at the 150 position. Therefore, fluorescence dequenching of the pure sfGFP-Tet protein upon reaction with sTCO-reagents served as a method to measure the rates of Tet-reaction on proteins. The fluorescence of sfGFP-Tet in 3 mL of PBS (15 nmol) was measured (488 nm excitation, 510 nm emission, 5 points/second) for 50 seconds prior to the addition of 10 µL sTCO -OH (0.03 -3 µmol sTCO reagent in PBS). The sTCO reagents were prepared in methanol at stock concentrations ranging from 10 to 900 µmol. Monitoring of the reactions continued until the fluorescence returned to stability. Subsequently, curves were fitted to a single exponential equation using the curve-fitting program OriginPro 8.5 to determine the pseudo-first-order kinetic constants. Next, the pseudo-first-order rate constant was plotted against the concentration of sTCO-OH to determine second-order rate constants of tetrazine reaction on sfGFP, as illustrated in Figures S12 and S13.

**Methods of eukaryotic expression and labeling of Tet-proteins.** Following published protocol Jang et al.[2]

**Table S1.** Summary table of the genetic encoding of Tet2-ncAAs and their reactivity inside the protein.

| Tet2-R | | | | | | | | | | |
|---|---|---|---|---|---|---|---|---|---|---|
| | Yield (mg/L) | Fidelity ratio | Host cells | | Fidelity (% of NCS in gel) | Fidelity (% of NCS in MS) | Stability on protein at RT | $k_2$ (x $10^4$ $M^{-1}s^{-1}$) | | Ratio[a] |
| | | | *E.coli* | HEK | | | | Free ncAA $k_2$ (1) | on protein $k_2$ (2) | |
| Me | 90 | 6.1 | ✓ | X | 6 | n.d. | --- | 2.95 ± 0.07 | 8.7 ± 0.3 | 2.9 |
| Et | 117 | 10.2 | ✓ | X | 4 | n.d. | --- | --- | 5.9 ± 0.4 | --- |
| Ip | 95 | 7 | ✓ | X | 8 | --- | --- | --- | 0.9 ± 0.05 | --- |
| Ph | --- | 1.1 | X | --- | --- | | | ---[b] | --- | --- |
| Pyr | --- | 1.3 | X | --- | --- | | | 20.4 ± 1.2 | --- | --- |

'a' ratio of second-order reaction kinetics ($k_2$) on protein and free ncAA. 'b' kinetics was not measured due to solubility issues. n.d. (not detectable).

**Table S2.** Summary table of the genetic encoding of Tet3-ncAAs and their reactivity inside the protein.

| Tet3-R | | | | | | | | | | |
|---|---|---|---|---|---|---|---|---|---|---|
| | Yield (mg/L) | Fidelity ratio | Host cells | | Fidelity (% of NCS in gel) | Fidelity (% of NCS in MS) | Stability at RT (% of reactive protein remaining after 8 d) | $k_2$ (x $10^4$ $M^{-1}s^{-1}$) | | Ratio[a] |
| | | | *E. coli* | *HEK* | | | | Free ncAA $k_2$ (1) | on protein $k_2$ (2) | |
| Me | 74 | 7.4 | ✓ | ✓ | 7 | n.d. | --- | 2.48 ± 0.08 | 7.8 ± 0.4 | 3.1 |
| Et | 74 | 7 | ✓ | ✓ | 4 | n.d. | --- | 2.22 ± 0.13 | 8.1 ± 0.20 | 3.6 |
| Ip | 67 | 6.1 | ✓ | ✓ | --- | --- | --- | 0.5 ± 0.07 | 2.3 ± 0.07 | 4.6 |
| Bu | 80 | 8 | ✓ | ✓ | 6 | n.d. | --- | 2.05 ± 0.06 | 7.8 ± 0.15 | 3.1 |
| Ph | 45 | 4 | ✓ | ✓ | 16 | 13 | >75% | ---[b] | 7.4 ± 0.38 | --- |
| 4-F-Ph | 42 | 3.6 | ✓ | ✓ | 22 | 10 | --- | ---[b] | 8.6 ± 0.60 | --- |
| 4-$NH_2$-Ph | 12 | 1.7 | ✓ | --- | 57 | 49 | --- | 1.81 ± 0.10 | 5.1 ± 0.30 | 2.8 |
| Pyr | 36 | 2.9 | ✓ | ✓ | 34 | 24 | <5% | 9.64 ± 0.98 | 23 ± 1.41 | 2.3 |

'a' ratio of second-order reaction kinetics ($k_2$) on protein and free ncAA. 'b' kinetics was not measured due to solubility issues. n.d. (not detectable).

**Table S3.** Summary table of the genetic encoding of Tet4-ncAAs and their reactivity inside the protein.

| Tet4-R | | | | | | | | | | | |
|---|---|---|---|---|---|---|---|---|---|---|---|
| | Yield (mg/L) | Fidelity ratio | | Host cells | | Fidelity (% of NCS in gel) | Fidelity (% of NCS in MS) | Stability at RT (% of reactive protein remaining after 3 d) | $k_2$ (x $10^4$ $M^{-1}s^{-1}$) | | Ratio[a] |
| | | RS-1 | RS-2 | *E. coli* | HEK | | | | Free ncAA | On protein | |
| Me | --- | 1.1 | 1.2 | X | --- | --- | --- | --- | 1.09 ± 0.14 | --- | --- |
| Et | --- | 0.9 | 1.1 | X | --- | --- | --- | --- | --- | --- | --- |
| Ip | --- | 0.9 | 0.9 | X | --- | --- | --- | --- | --- | --- | --- |
| Bu | 5 | 3.5 | 2.7 | ✓ | --- | 30 | --- | --- | --- | 21 ± 1.5 | --- |
| Ph | 55 | 16 | 13 | ✓ | ✓ | 5 | n.d. | > 90% | 2.42 ± 0.19 | 22 ± 1.2 | 9.1 |
| 3-Me-Ph | 40 | 13 | 11 | ✓ | ✓ | 4 | n.d. | > 90% | --- | 34 ± 2.3 | --- |
| 4-Me-Ph | 15 | 5 | 4.6 | ✓ | --- | 26 | --- | >80% | --- | 44 ± 3.1 | --- |
| 3-F-Ph | 44 | 13 | 12 | ✓ | ✓ | 3 | n.d. | >80% | --- | 46 ± 2.3 | --- |
| 4-F-Ph | 18 | 5.5 | 5 | ✓ | ✓ | 3 | n.d. | --- | 2.52 ± 0.26 | 45 ± 1.5 | 18 |
| 4-OH-Ph | --- | 0.8 | 0.8 | X | --- | --- | --- | --- | 0.7 ± 0.1 | --- | --- |
| 4-$NH_2$-Ph | --- | 1.3 | 1.1 | X | --- | --- | --- | --- | 2.26 ± 0.22 | --- | --- |
| 4-COMe -Ph | --- | 1.2 | 0.9 | X | --- | --- | --- | --- | --- | --- | --- |
| 3-$CF_3$-Ph | 6 | 3.5 | 3.2 | ✓ | --- | 28 | 15 | >80% | --- | 62 ± 2.2 | --- |
| Pyr | 35 | 2 | 7.5 | ✓ | ✓ | 32 | 18 | <10% | 8.59 ± 1.19 | 120 ± 7.3 | 13.9 |
| 3,5-diF-Ph | 67 | 14 | 4.5 | ✓ | ✓ | 4 | n.d. | >85% | --- | 84 ± 3.9 | --- |
| 3,4,5-triF-Ph | 42 | 12 | 1.5 | ✓ | ✓ | 5 | n.d. | >80% | --- | 110 ± 9.4 | --- |

'a' ratio of second-order reaction kinetics ($k_2$) on protein and free ncAA. n.d. (not detectable).

**Toxicity screen of synthesized Tet-ncAAs.**

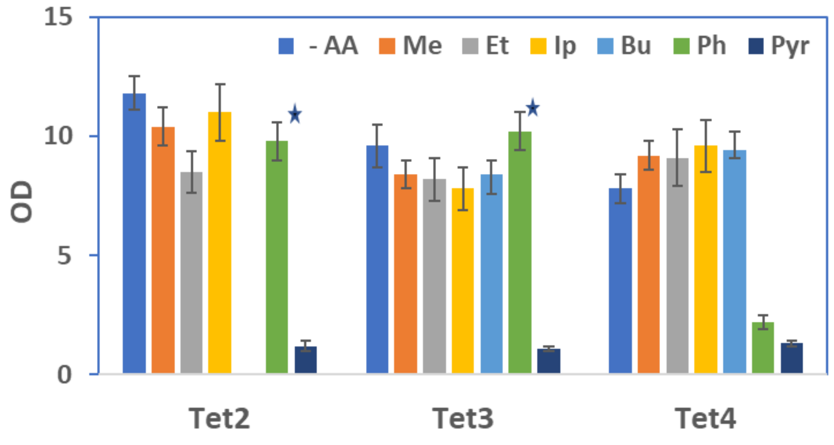

**Figure S1**: Measured optical density ($OD_{600}$) of fresh cultured DH10B *E. coli* cells in 5 mL auto-inducing media (AIM) in the presence and absence of 1 mM Tet2, Tet3, and Tet4 (-Me, -Et, -Ip, Bu, -Ph, and -Py) derivatives after 40 h. The cell's growth temperature was 37 $^{o}$C. Asterisks (*) indicate that the Tet2-Ph and Tet3-Ph concentrations were 0.25 mM due to low solubility in dimethylformamide (DMF).

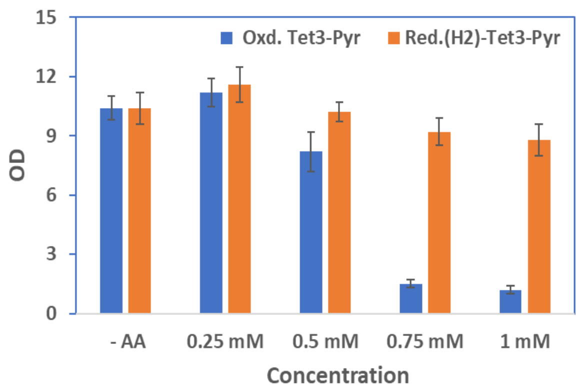

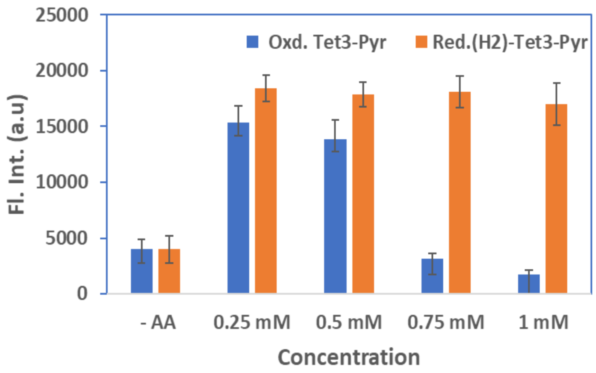

**Figure S2**: Measured cell density ($OD_{600}$) and fluorescence intensity of expressed GFP150-Tet3-Py using Tet3-Py (oxidized tetrazine, blue) and 1,4 Dihydro ($H_2$)-Tet3-Py (reduced tetrazine, orange) side by side. Cells were inoculated with and without Tet3-Py at different concentrations: 0.25 mM, 0.5 mM, 0.75 mM and 1 mM (50 mL AIM, 32 h. at 37 $^{o}$C).

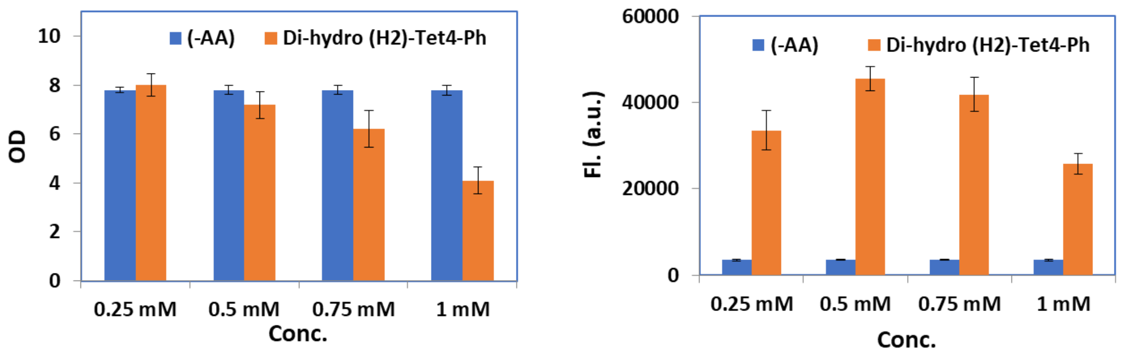

**Figure S3**: Measured cell density ($OD_{600}$) and fluorescence intensity of expressed sfGFP150-Tet4-Ph in the absence (blue) and presence (orange) of 1,4 Dihydro ($H_2$)-Tet4-Ph (reduced tetrazine) with varying concentrations 0.25 mM, 0.5 mM, 0.75 mM and 1 mM (50 mL AIM, 32 hrs. at 37$^{o}$C). 1,4 Dihydro ($H_2$)-Tet4-Ph supplemented into the AIM media after 4 – 6 h. of cell inoculation when the cells are healthy growth phase ($OD_{600}$ ~ 0.5 to 0.7).

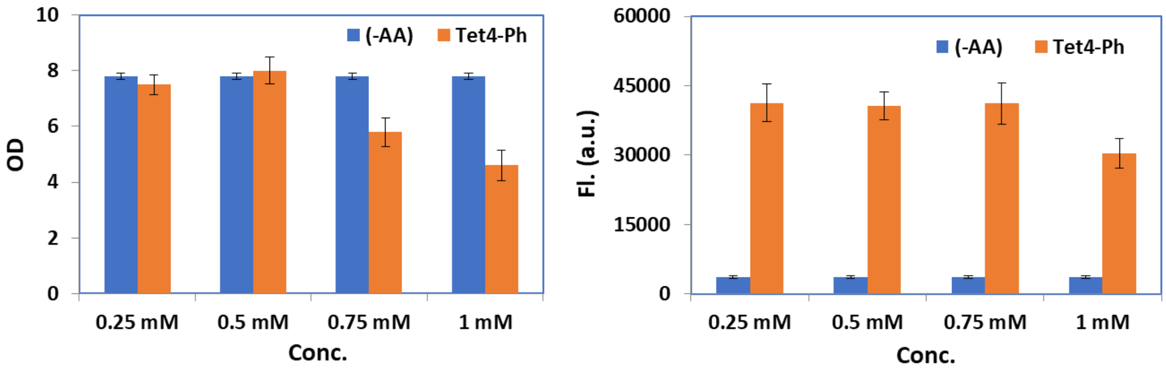

**Figure S4**: Measured cell density ($OD_{600}$) and fluorescence intensity of expressed sfGFP150-Tet4-Ph in absence (blue) and presence (orange) of Tet4-Ph (oxidized tetrazine) with varying concentrations 0.25 mM, 0.5 mM, 0.75 mM and 1 mM (50 mL AIM, 32 hrs. at 37$^{o}$C). Tet4-Ph supplemented into the AIM media after 4 – 6 hrs. of cell inoculation when the cells are healthy growth phase ($OD_{600}$ ~ 0.5 to 0.7).

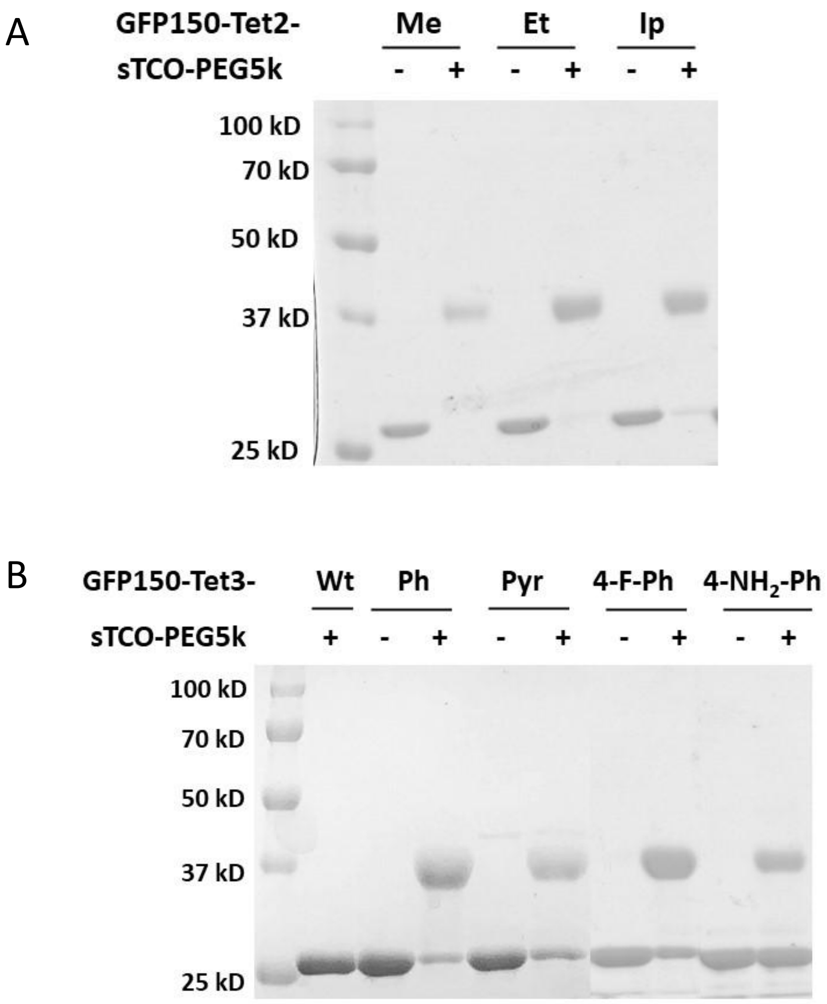

**Figure S5. SDS-PAGE mobility shift assay**. Labeling efficiency of (A) sfGFP-Tet2-(Me, Et, Ip) and (B) sfGFP-Tet3-(Ph, Pyr, 4-F-Ph, 4-$NH_2$-Ph) verified by SDS-PAGE mobility shift upon reaction with sTCO-PEG5k in PBS (pH~7.1).

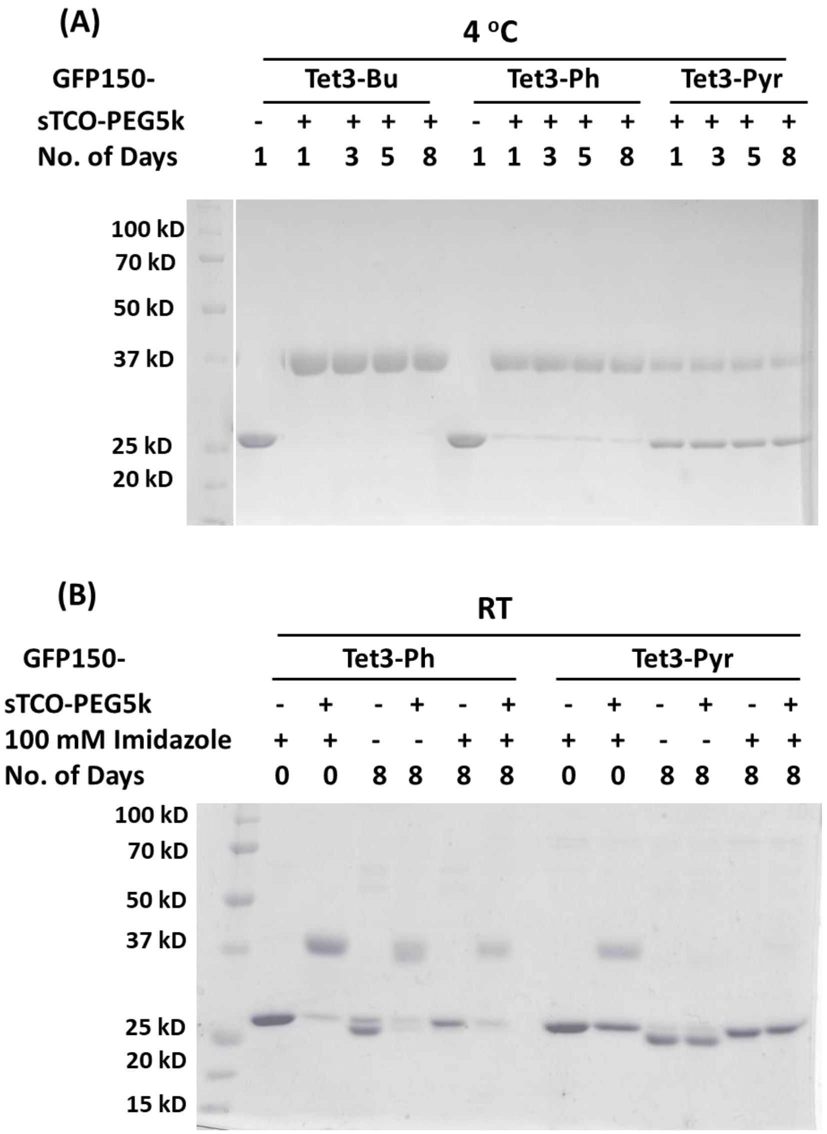

**Figure S6. Verification of stability and labeling efficiency of Tet3 on protein for 8 days incubation in PBS (pH~7.1) at 4 °C and room temperature (RT)**. SDS-PAGE mobility shift assay by reacting sTCO-PEG5k, verified stability and reaction ability of encoded Tet3-(Ph, and Pyr) stored at (A) 4 °C and (B) RT for eight days in the presence of 100 mM imidazole.

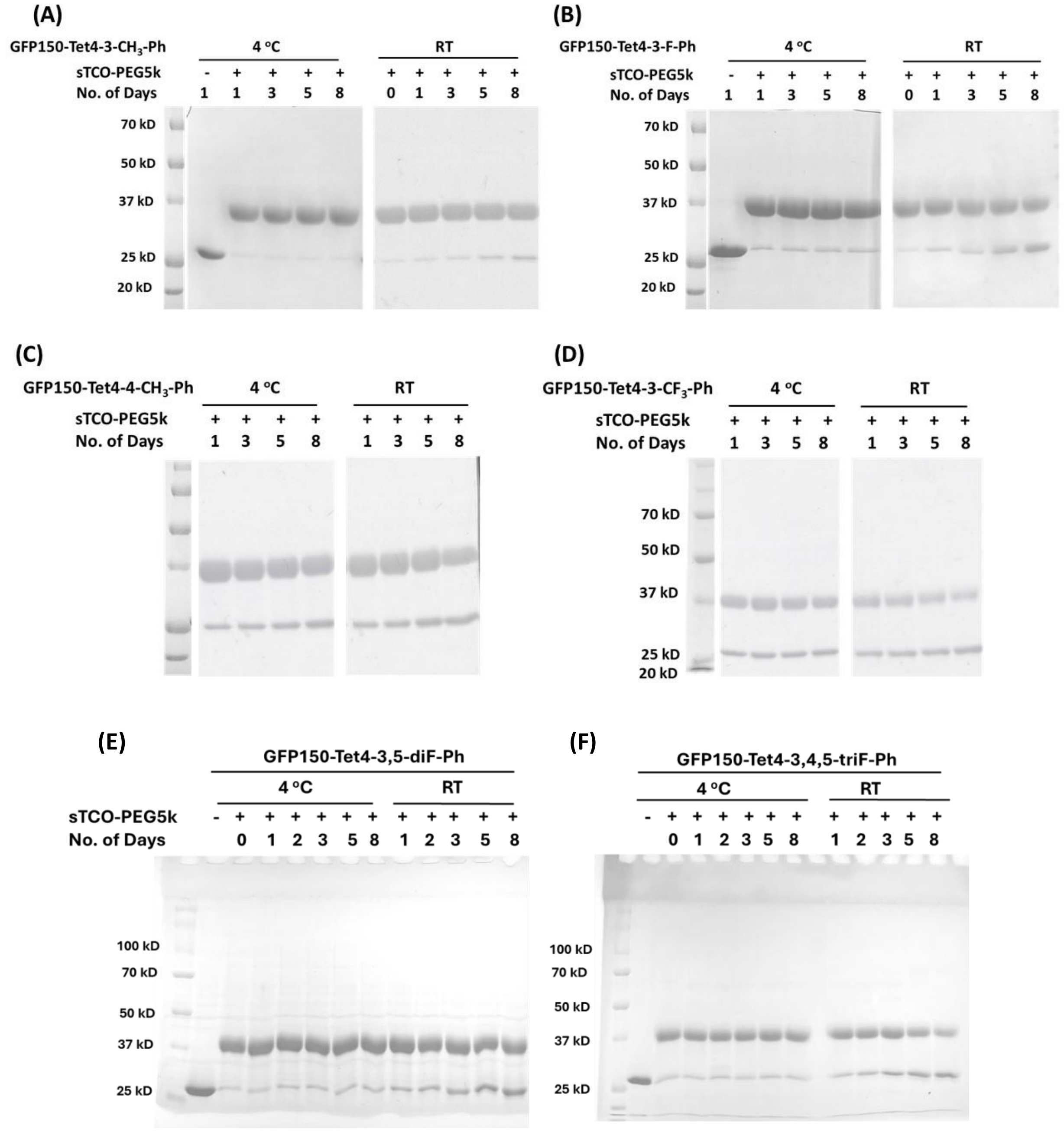

**Figure S7.** Verification of stability and labeling efficiency of Tet4 on protein for 8 days incubation in PBS (pH~7.1) at 4 $^{o}$C and room temperature (RT). SDS-PAGE mobility shift assay by reacting sTCO-PEG5k, verified stability and reactivity of (A) sfGFP-Tet4-3$CH_3$-Ph, (B) sfGFP-Tet4-3F-Ph, (C) sfGFP-Tet4-4$CH_3$-Ph, (D) sfGFP-Tet4-3$CF_3$-Ph, (E) sfGFP-Tet4-3,5-diF-Ph, and (F) sfGFP-Tet4-3,4,5-triF-Ph for 8 days at 4 $^{o}$C and RT in presence of 100 mM imidazole.

**Mass spectra analysis of genetically incorporated Tet3-ncAA derivatives into sfGFP**

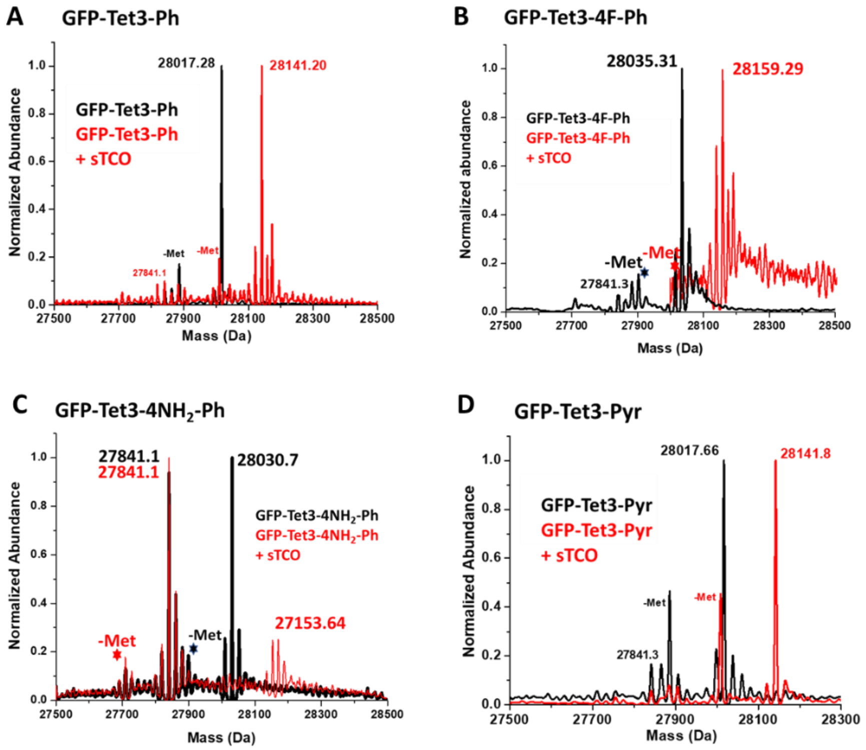

**Figure S8. ESI-Q-TOF mass spectrometry analyzed the incorporation efficiency of Tet3-Ph derivatives and amplitude of labeling reaction with sTCO**. ESI mass spectrometry analysis of GFP-Tet3 (-Ph, -4F-Ph, -4$NH_2$-Ph, -Pyr) and reactions with sTCO-OH**.** The purified sfGFP-Tet3-ncAAs (black) and reacted with 5-fold molar excess of sTCO-OH for 10 minutes (red) in PBS (pH~7.1). The reacted sfGFP-Tet3 proteins showed as expected 124 Da increase in mass corresponding to the addition of sTCO-OH and loss of molecular nitrogen. No unreacted sfGFP-Tet3 (-Ph, -4F-Ph, -Pyr) was detected, verifying the reactivity of encoded Tet3-Ph/4F-Ph/Pyr with sTCO-OH was quantitative. The lower mass peak labeled with * is a loss of n-terminal methionine and the upper mass peaks are salt sodium and potassium adducts. Whereas, the unreacted lower peak at 27841 Da avg. observed due to near-cognate suppression of amber codon. Near-cognate suppression was predominantly observed for sfGFP-Tet3-4-$NH_2$-Ph due to lower incorporation efficiency of working synthetase R2-84. Cal. Mass of sfGFP-wt: 27827.02 Da avg; **(A)** sfGFP-Tet3-Ph observed: 28017.28 Da avg, (expected: 28016.17 Da avg); sfGFP-Tet3-Ph + sTCO-OH observed: 28141.20 Da avg, (expected: 28140.18 Da avg). **(B)** sfGFP-Tet3-4F-Ph observed: 28035.31 Da avg, (expected: 28034.16 Da avg); sfGFP-Tet3-4F-Ph + sTCO-OH observed: 28159.29 Da avg, (expected: 28158.17 Da avg). **(C)** sfGFP-Tet3-4$NH_2$-Ph observed: 27841.1 Da avg (near-cognate suppression); 28030.75 Da avg, (expected: 28031.18 Da avg); sfGFP-Tet4-4-$NH_2$-Ph + sTCO-OH observed: 27841.1 Da avg (NCS); 28153.64 Da avg, (expected: 28155.19 Da avg). **(D)** sfGFP150-Tet3-Pyr observed 28017.66 Da avg (expected: 28017.1 Da avg). sfGFP-Tet3-Pyr + sTCO-OH observed: 28141.8 Da avg. (expected: 28141.1 Da avg.)

### Stability assessment of Tet3-Ph and Tet3-Py inside protein by MS analysis

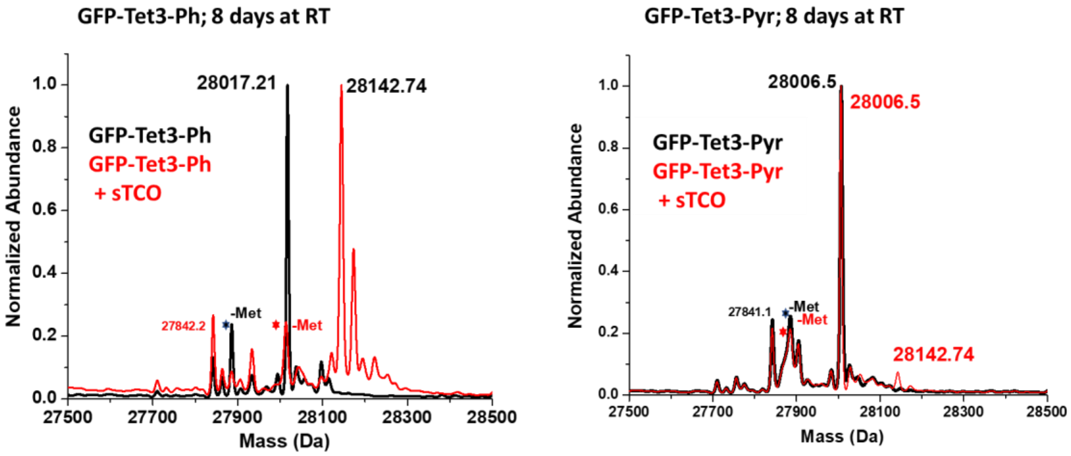

**Figure S9. Verification of stability and reactivity of Tet3-Ph and Tet3-Pyr inside protein by ESI-Q-TOF mass spectrometry after 8 days incubation at room temperature**. ESI mass spectrometry analysis of pure sfGFP-Tet3-Ph/Py in absence and presence of sTCO after 8 days incubation in PBS (pH~7.1) at room temperature under basic conditions (100 mM imidazole was added). Purified sfGFP-Tet3-ncAAs (black) and upon reaction with 5-fold molar excess of sTCO-OH for 10 minutes (red). The sfGFP-Tet3-Ph showed quantitative labeling by the addition of sTCO-OH. No unreacted sfGFP-Tet3-Ph was detected, which verified that the genetically encoded Tet3-Ph is stable enough for long time incubation. Whereas, no reaction was observed for GFP-Tet-Pyr. The lower mass peak labeled with * is a loss of n-terminal methionine and upper mass peaks are salt sodium and potassium adducts. The lower peak at 27842.2 Da avg. observed due to near-cognate suppression of amber codon. **(A)** sfGFP-Tet3-Ph observed: 28017.21 Da avg, (expected: 28016.17 Da avg); sfGFP-Tet3-Ph + sTCO-OH observed: 28142.74 Da avg, (expected: 28140.18 Da avg). **(B)** In presence and absence of sTCO, it shows an unreactive single major peak at 28006.5 Da avg. which is 11 Da avg. unit lower than expected. (sfGFP-Tet3-Py expected: 28017.1 Da avg, and sfGFP-Tet3-Pyr+sTCO expected: 28141.1 Da avg). The MS result indicated that the sfGFP-Tet3-Pyr degraded under the following conditions and converted to its oxadiazole derivative which is 12 Da lower molecular mass than Tet3-Pyr.[6]

### Tet3-Pyr and Tet4-Pyr degradation inside sfGFP[6]

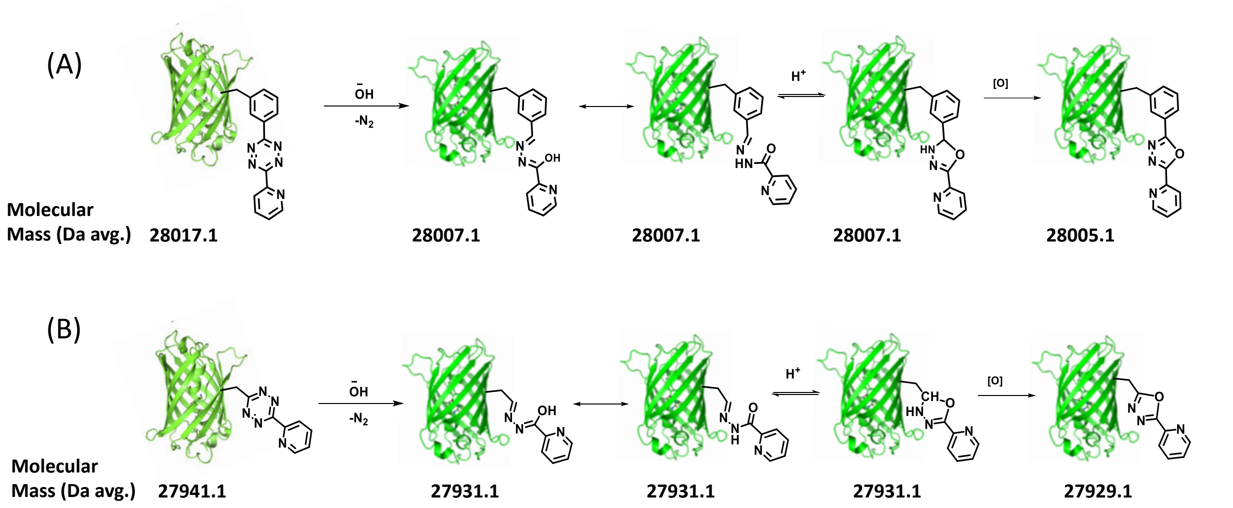

**Figure S10:** Proposed mechanism of (A) Tet3-Pyr (B) Tet4-Pyr side reactions and chemical structure of corresponding degraded product 1,3,4-oxadiazole derivatives.

**Table S4**: Mass Spectra analysis of *E. coli* expressed GFP150-Tet-ncAA variants with and without sTCO.

| ***E. coli* expressed sfGFP150-Tet-ncAA** | | | | | |
|---|---|---|---|---|---|
| Proteins | Residues at TAG sites | Theoretical Mass (Da avg.) | Observed Mass (Da avg.) | Difference (Da) | Modifications |
| wt-sfGFP | Asn150 | 27827.1 | 27826.8 | -0.3 | - |
| **(A) Analyzed incorporation and labeling of sfGFP150-Tet3** | | | | | |
| sfGFP-Tet3-Ph | Asn150/ Tet3-Ph | 28016.17 | 28017.28 | +1.11 | Tet3-Ph incorporation |
| | | | 27885.82 | -130.35 | -Met. |
| | | | 27841.10 (near-cognate supp.) | -175.08 | Glu. incorporation (minor) |
| sfGFP- Tet3-Ph + sTCO | Asn150/ Tet-Tet3-Ph | 28140.18 | 28141.2 | +1.02 | sTCO addition and $N_2$ loss |
| | | | 28011.1 | -129.1 | -Met. |
| | | | 27841.10 (unreactive) | -299.08 | Glu. incorporation (minor) |
| sfGFP-Tet3-4F-Ph | Asn150/ Tet3-4F-Ph | 28034.16 | 28035.31 | +1.15 | Tet3-4F-Ph incorporation |
| | | | 27903.9 | -130.26 | -Met. |
| | | | 27841.3 (near-cognate supp.) | -192.86 | Glu. incorporation (minor) |
| sfGFP-Tet3-4F + sTCO | Asn150/ Tet3-4F-Ph | 28158.17 | 28159.29 | +1.12 | sTCO addition and $N_2$ loss |
| | | | 28029.1 | -129.07 | -Met. |
| sfGFP-Tet3- $4NH_2$-Ph | Asn150/ Tet3- $4NH_2$-Ph | 28031.18 | 28030.75 | -0.42 | Tet3-$4NH_2$-Ph incorporation |
| | | | 27899.9 | -131.28 | -Met. |
| | | | 27841.1 (near-cognate supp.) | -190.08 | Glu. incorporation (major) |
| | | | 27708.9 | -322.28 | -Met. w.r.to GFP150-Glu |
| sfGFP- Tet3- $4NH_2$ + sTCO | Asn150/ Tet3- $4NH_2$-Ph | 28155.19 | 28153.64 | -1.55 | sTCO addition and $N_2$ loss (minor) |
| | | | 27841.1 (unreactive) | -314.09 | Glu. incorporation (major) |
| | | | 27709.5 | -445.69 | -Met. w.r.to GFP150-Glu |
| sfGFP-Tet3-Pyr | Asn150/ Tet3-Pyr | 28017.1 | 28017.66 | +0.56 | Tet3-Pyr incorporation |
| | | | 27886.41 | -130.69 | -Met. |
| | | | 27841.3 (near-cognate supp.) | 175.8 | Glu. incorporation (minor) |
| sfGFP-Tet3-Pyr + sTCO | Asn150/ Tet3-Pyr | 28141.1 | 28141.8 | +0.7 | sTCO addition and $N_2$ loss |
| | | | 28010.6 | -130.5 | -Met. |

| **(B) Analyzed incorporation and labeling of sfGFP150-Tet4** | | | | | |
|---|---|---|---|---|---|
| sfGFP-Tet4-Ph | Asn150/ Tet4-Ph | 27940.1 | 27941.5 | +1.4 | Tet4-Ph incorporation |
| | | | 27810.17 | -130 | -Met |
| sfGFP-Tet4-Ph +sTCO | Asn150/ Tet4-Ph | 28064.1 | 28065.5 | +1.4 | sTCO addition and $N_2$ loss |
| | | | 27934.12 | -130 | -Met |
| sfGFP-Tet4-3Me-Ph | Asn150/ Tet4-3Me-Ph | 27954.15 | 27955.1 | +0.95 | Tet4-3Me-Ph incorporation |
| | | | 27823.17 | -130.98 | -Met |
| sfGFP-Tet4-3Me-Ph + sTCO | Asn150 Tet4-3Me-Ph | 28078.16 | 28079.34 | +1.18 | sTCO addition and $N_2$ loss |
| | | | 27948.46 | -129.7 | -Met |
| sfGFP-Tet4-3F-Ph | Asn150/ Tet4-3F-Ph | 27958.13 | 27959.2 | +1.07 | Tet4-3F-Ph incorporation |
| | | | 27827.25 | -130.88 | -Met |
| sfGFP- Tet4-3F-Ph + sTCO | Asn150/ Tet4-3F-Ph | 28082.14 | 28082.84 | +0.7 | sTCO addition and $N_2$ loss |
| | | | 27953.13 | -129.01 | -Met |
| sfGFP-Tet4-3CF$_3$-Ph | Asn150/ Tet4-3CF$_3$-Ph | 28008.12 | 28008.22 | +0.1 | Tet4-3CF$_3$-Ph incorporation |
| | | | 27876.45 | -131.67 | -Met |
| | | | 27840.8 (near-cognate supp.) | -167.32 | Glu. incorporation (minor) |
| sfGFP-Tet4-3CF$_3$-Ph + sTCO | Asn150/ Tet4 3CF$_3$-Ph | 28132.13 | 28132.49 | +0.36 | sTCO addition and $N_2$ loss |
| | | | 27800.74 | -131.39 | -Met |
| | | | 27841.3 (unreactive) | -290.83 | Glu. incorporation (minor) |
| sfGFP-Tet4-Pyr | Asn150/ Tet4-Pyr | 27941.1 | 27941.3 | +0.2 | Tet4-Pyr incorporation |
| | | | 27810.6 | -130.5 | -Met |
| sfGFP-Tet4-Pyr+sTCO | Asn150/ Tet4-Pyr | 28065.11 | 28066.2 | +1.1 | sTCO addition and $N_2$ loss |
| | | | 27933.8 | -131.3 | -Met |
| sfGFP-Tet4-3,5-diF-Ph | Asn150/ Tet4-3,5-diF-Ph | 27976.1 | 27976.7 | +0.5 | Tet4-3,5-diF-Ph incorporation |
| | | | 27844.2 | -131.9 | -Met |
| sfGFP-Tet4-3,5-diF-Ph+ sTCO | Asn150/ Tet4-3,5-diF-Ph | 28100.1 | 28100.4 | +0.3 | sTCO addition and $N_2$ loss |
| | | | 27969.3 | -131.1 | -Met |
| sfGFP-Tet4-3,4,5-triF-Ph | Asn150/ Tet4-3,4,5-triF-Ph | 27994.1 | 27993.8 | -0.3 | Tet4-3,4,5-triF-Ph incorporation |
| | | | 27862.7 | -131.4 | -Met |
| sfGFP-Tet4-3,4,5-triF-Ph +sTCO | Asn150/ Tet4-3,4,5-triF-Ph | 28118.1 | 28118.2 | +0.1 | sTCO addition and $N_2$ loss |
| | | | 27987.1 | -131 | -Met |

| | | | | | |
|---|---|---|---|---|---|
| **(C) Stability assessment of Tet3-Ph and Tet3-Py inside protein**<br>(All samples were sited at room temperature for 8 days) | | | | | |
| sfGFP-Tet3-Ph | Asn150/<br>Tet3-Ph | 28016.17 | 28017.28 | +1.11 | Tet3-Ph incorporation |
| | | | 27885.82 | -130.35 | -Met. |
| | | | 27840.95<br>(near-cognate supp.) | -175.08 | Glu. incorporation (minor) |
| sfGFP- Tet3-Ph + sTCO | Asn150/<br>Tet3-Ph | 28140.18 | 28142.74 | +1.02 | sTCO addition and $N_2$ loss |
| | | | 28012.66 | -129.1 | -Met. |
| | | | 27842.29<br>(unreactive) | -299.08 | Glu. incorporation (minor) |
| sfGFP-Tet3-Pyr | Asn150/<br>Tet3-Pyr | 28017.1 | 28006.5 | -10.6 | Incorporated Tet3-Pyr degradation and oxadiazole formation |
| | | | 27885.82 | -131.28 | -Met. |
| | | | 27841.1<br>(near-cognate supp.) | -176.0 | Glu. incorporation (minor) |
| sfGFP-Tet3-Pyr + sTCO | Asn150/<br>Tet3-Pyr | 28141.1 | 28142.74 | +1.64 | sTCO addition and $N_2$ loss (minor) |
| | | | 28006.5<br>(unreactive) | -134.6 | Unreactive due to tetrazine functionality loses. |
| | | | 27885.82 | -255.28 | -Met. |
| | | | 27841.1<br>(unreactive) | -300.0 | Glu. incorporation (minor) |
| **(D) Stability assessment of Tet4-Ph and Tet4-Py inside protein**<br>(All samples were sited at room temperature for 8 days) | | | | | |
| sfGFP-Tet4-Ph | Asn150/<br>Tet4-Ph | 27940.1 | 27941.23 | +1.13 | Tet4-Ph incorporation |
| | | | 27810.17 | -129.93 | -Met. |
| sfGFP-Tet4-Ph +sTCO | Asn150/<br>Tet4-Ph | 28064.1 | 28065.2 | +1.1 | sTCO addition and $N_2$ loss |
| | | | 27935.10 | -129.0 | -Met. |
| sfGFP-Tet4-Pyr | Asn150/<br>Tet4-Pyr | 27941.1 | 27930.5 | -10.6 | Incorporated Tet4-Pyr degradation and oxadiazole formation |
| | | | 27810.1 | -131.0 | -Met. |
| sfGFP-Tet4-Pyr +sTCO | Asn150/<br>Tet4-Pyr | 28065.11 | 28066.8 | +1.69 | sTCO addition and $N_2$ loss (minor) |
| | | | 27929.9<br>(unreactive) | -135.21 | Unreactive due to tetrazine functionality loses. |
| | | | 27810.1 | -255.01 | -Met. |

### Kinetics of Tet-ncAA inside protein

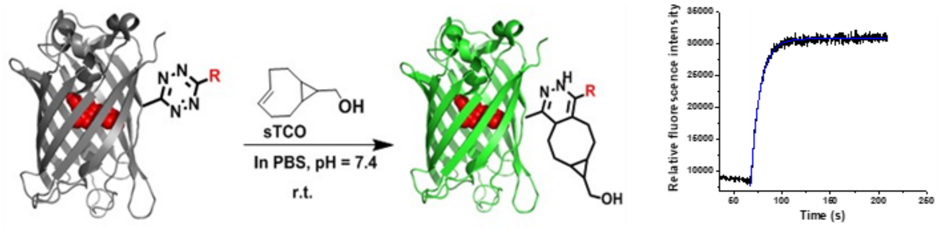

**Figure S11.** Tet-ncAAs reactivity inside protein was assessed by measuring the regenerated fluorescence of quenched sfGFP150-Tet with time when exposed to react with sTCO in PBS (pH ~7.1) at 25 $^oC$.

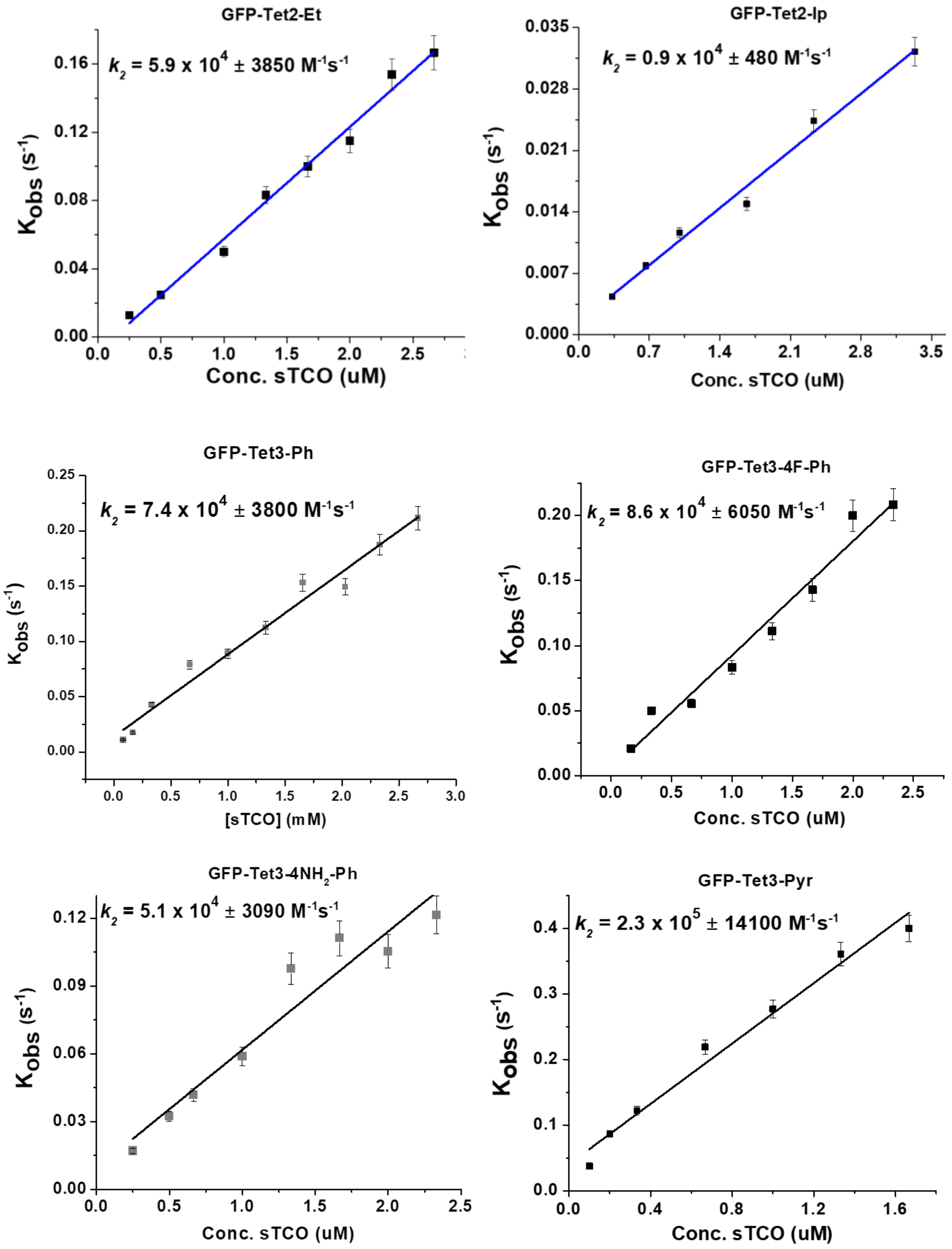

**Figure S12. Measured reaction rate for Tet2 and Tet3 derivatives**. Plot of pseudo first-order rate constant ($k'$) against concentration of sTCO to determine the second-order rate constant ($k_2$) for reaction of sfGFP-Tet2 and sfGFP-Tet3 with sTCO.

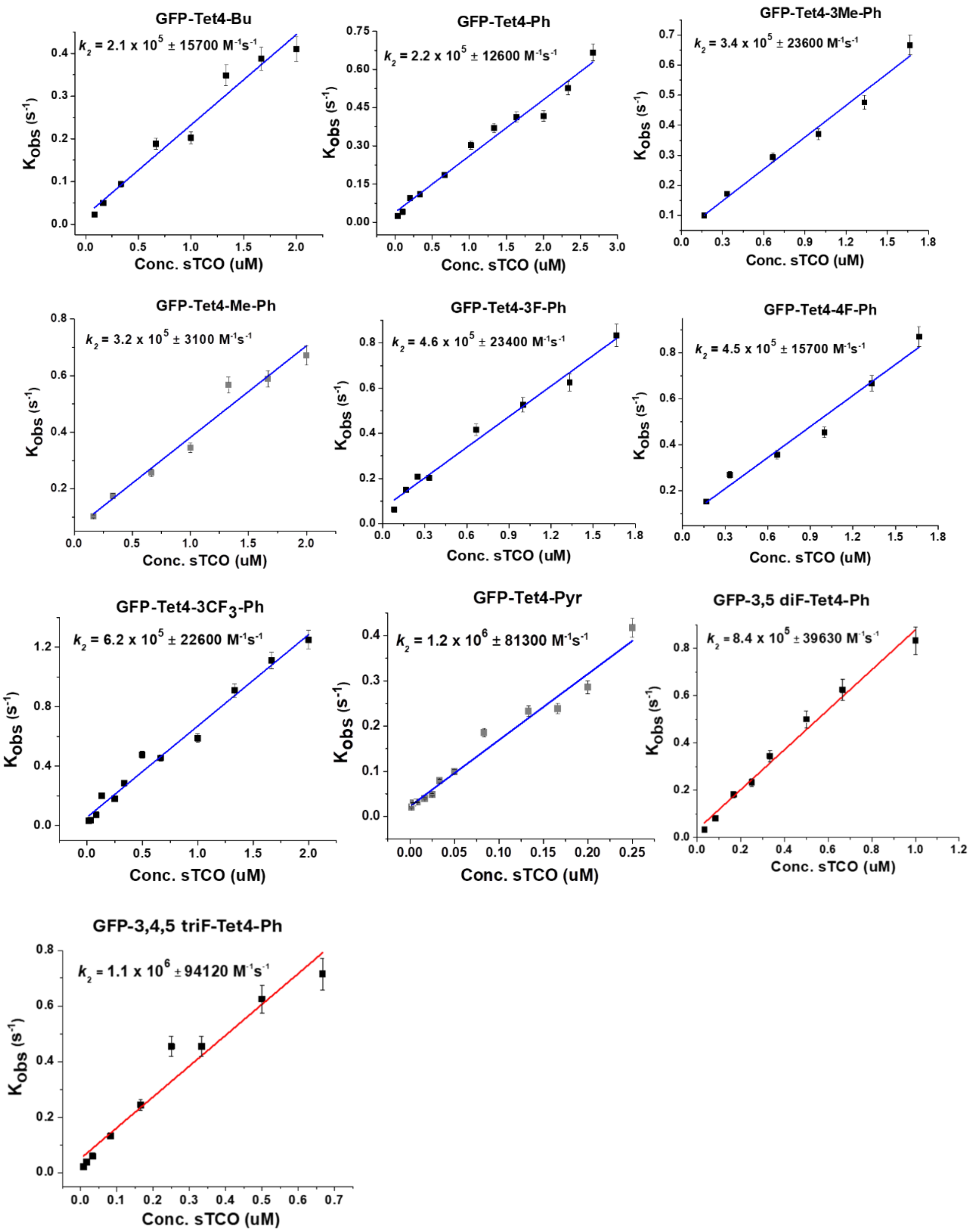

**Figure S13. Measured reaction rate for Tet4 derivatives**. Plot of pseudo-first-order rate constant ($k^{'}$) against concentration of sTCO to determine the second-order rate constant ($k_2$) for the reaction of sfGFP-Tet4 with sTCO.

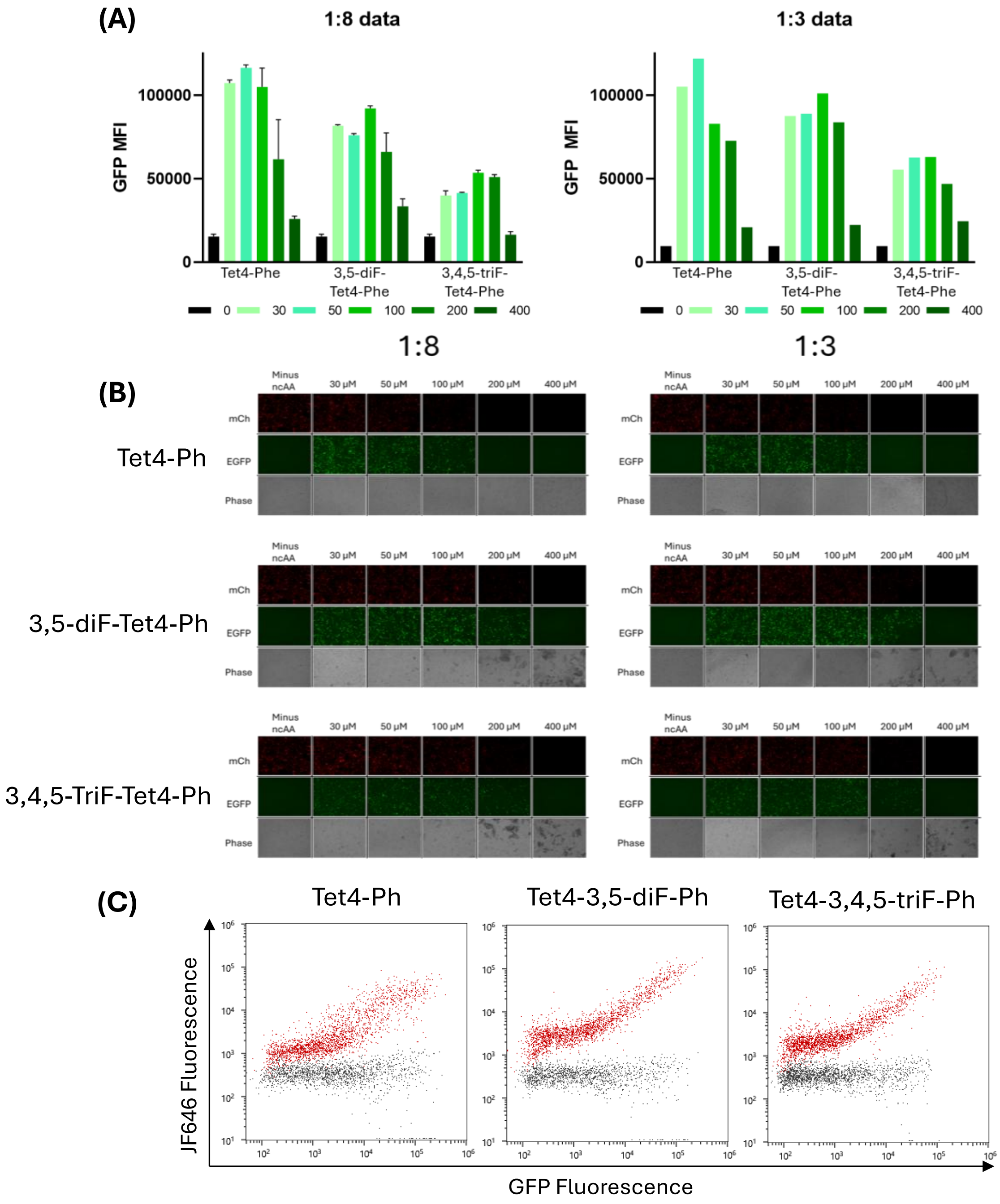

**Figure S14**. Evaluating the expression and labeling of fluorinated Tet4-Ph variants, Tet4-3,5-diF-Ph and Tet4-3,4,5-triF-Ph in mammalian cells (A) Expression of mCherry-linker$^{250}$-eGFP[7] using the three Tet4-Ph variants at 1:8 (left, N=2) and 1:3 (right, N=1) ratio of pAcBac1-E1RS-4x*MbPyl*-tRNA to pAcBac1-mCherry-linker$^{250}$-eGFP-4x*MbPyl*-tRNA machinery and expression plasmids. Different ncAA concentrations were tested for each ncAA, from 0 to 400 µM ncAA. (B) Fluorescent imagery shows that by 200 µM, significantly lower levels of cells are present due to toxicity. (C) Live in cell labeling of the Tet4-Ph and the new fluorinated Tet4 ncAAs encoded in sfGFP$^{150}$. Cells were incubated with sTCO-JF646 (100 nM) for 60 min prior to analysis. Expressions were performed for 24 h before labeling assays and FACS measurements.

**$^1$H and $^{13}$C NMR spectra of synthesized Tet-ncAAs:**

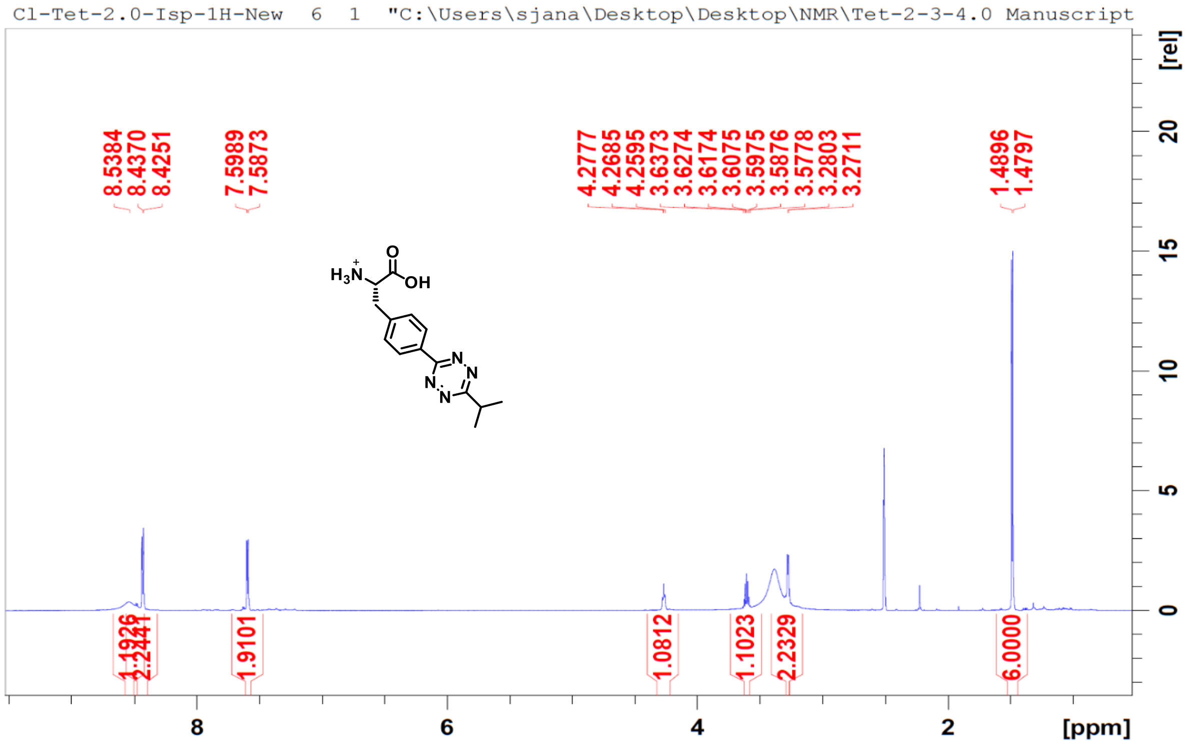

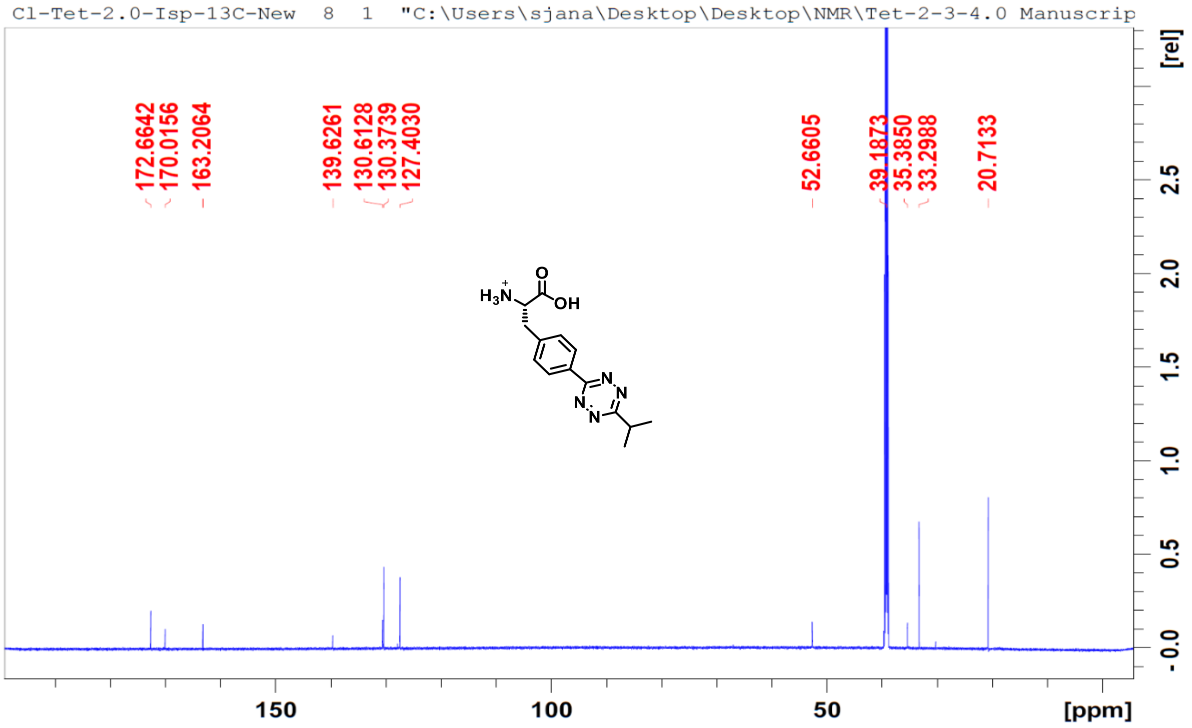

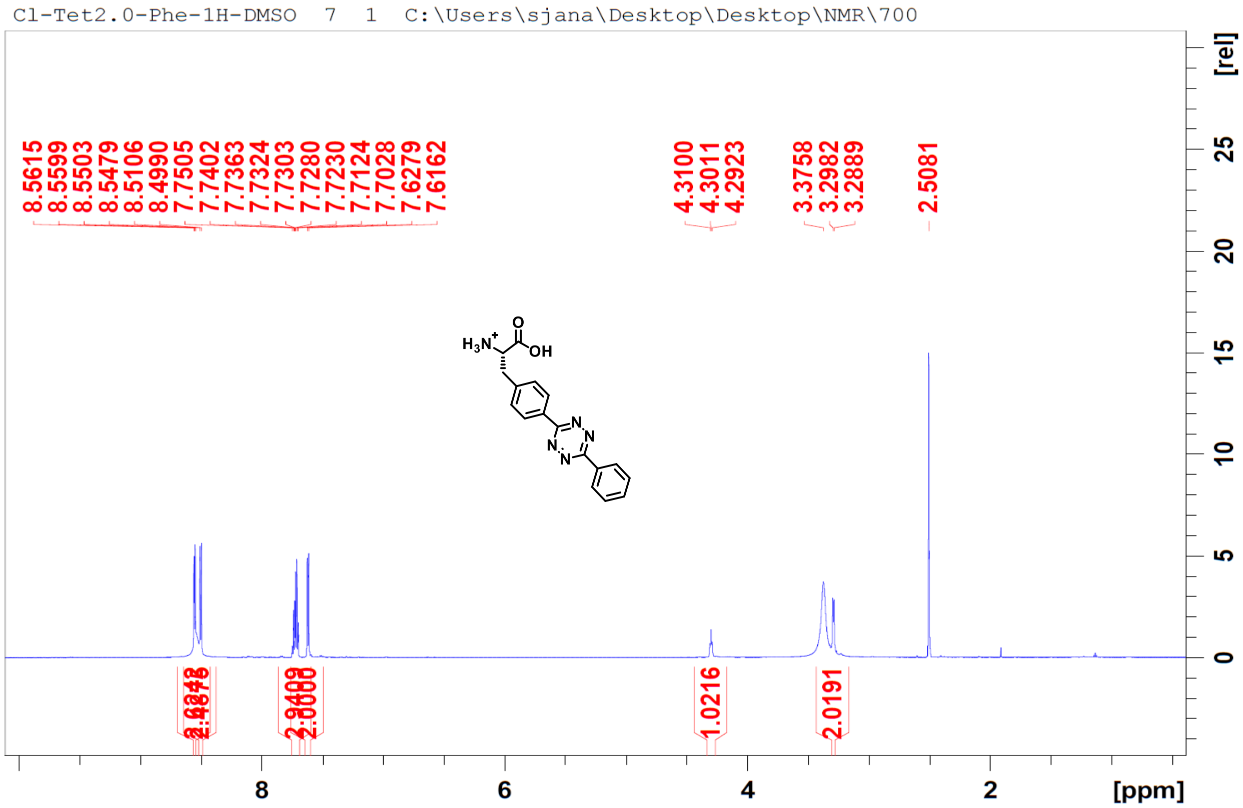
Cl-Tet2.0-Phe-1H-DMSO 7 1 C:\Users\sjana\Desktop\Desktop\NMR\700
8.5615
8.5599
8.5503
8.5479
8.5106
8.4990
7.7505
7.7402
7.7363
7.7324
7.7303
7.7280
7.7230
7.7124
7.7028
7.6279
7.6162
4.3100
4.3011
4.2923
3.3758
3.2982
3.2889
2.5081
1.0216
2.0191
[rel]
[ppm]

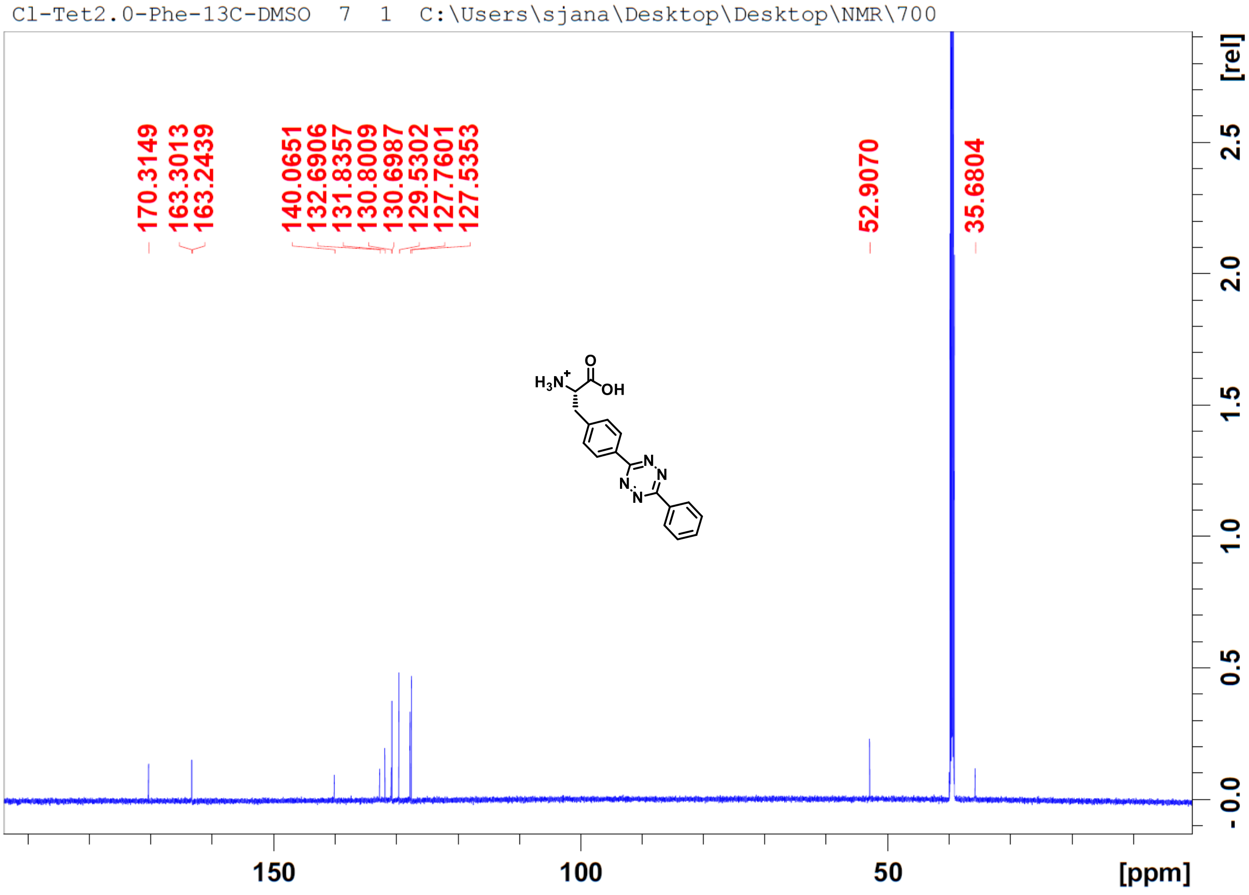
Cl-Tet2.0-Phe-13C-DMSO 7 1 C:\Users\sjana\Desktop\Desktop\NMR\700
170.3149
163.3013
163.2439
140.0651
132.6906
131.8357
130.8009
130.6987
129.5302
127.7601
127.5353
52.9070
35.6804
[rel]
[ppm]

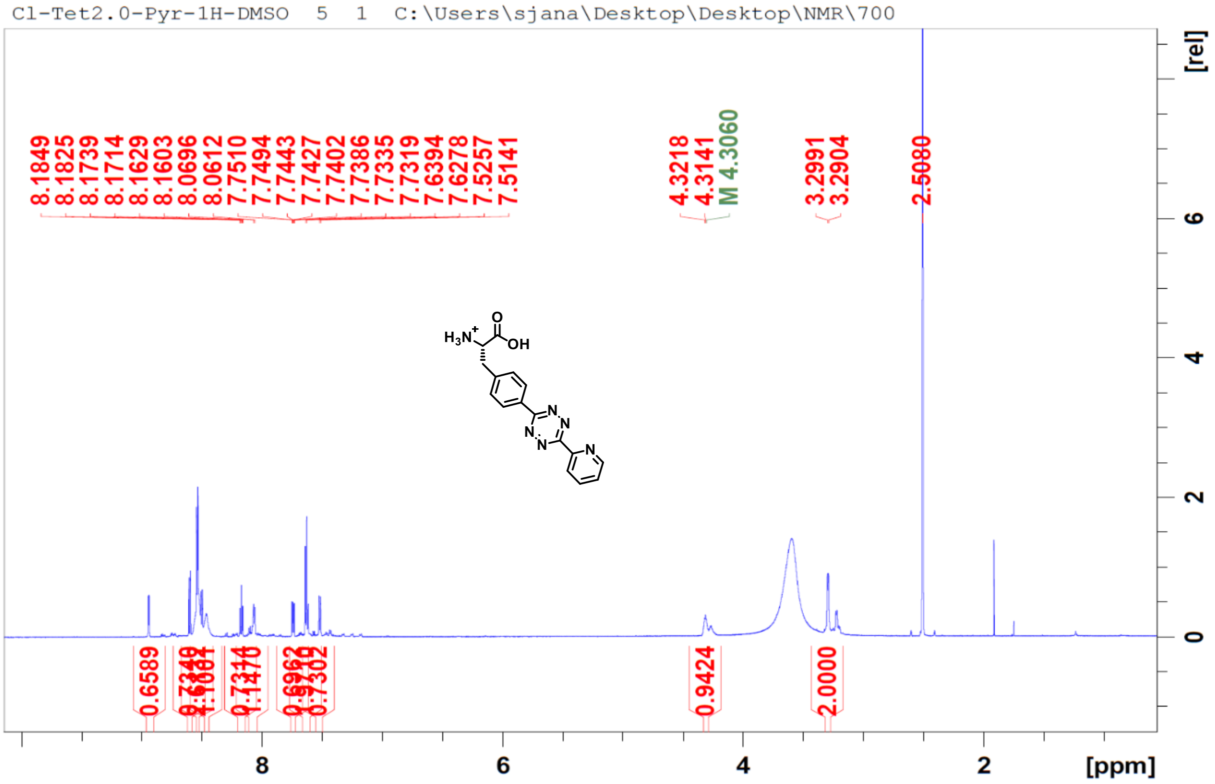
Cl-Tet2.0-Pyr-1H-DMSO 5 1 C:\Users\sjana\Desktop\Desktop\NMR\700
8.1849
8.1825
8.1739
8.1714
8.1629
8.1603
8.0696
8.0612
7.7510
7.7494
7.7443
7.7427
7.7402
7.7386
7.7335
7.7319
7.6394
7.6278
7.5257
7.5141
4.3218
4.3141
M 4.3060
3.2991
3.2904
2.5080
0.6589
0.9424
2.0000
[rel]
[ppm]

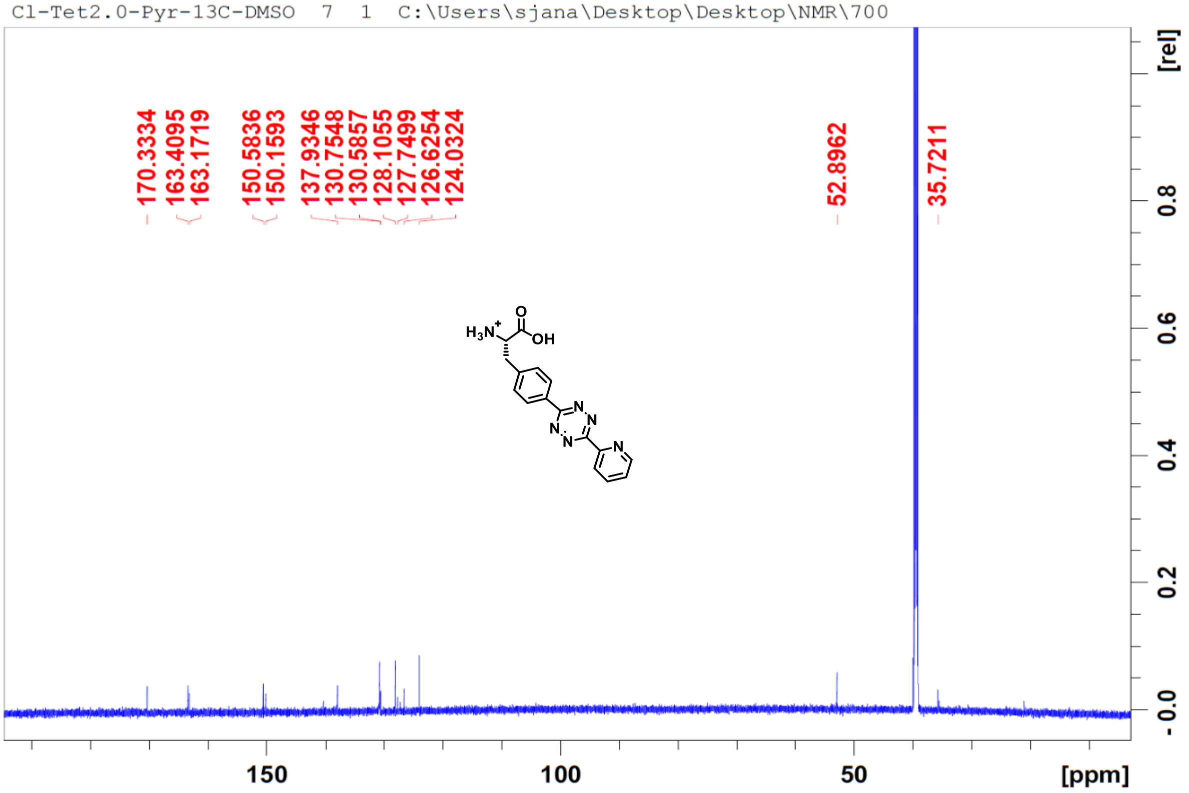
Cl-Tet2.0-Pyr-13C-DMSO 7 1 C:\Users\sjana\Desktop\Desktop\NMR\700
170.3334
163.4095
163.1719
150.5836
150.1593
137.9346
130.7548
130.5857
128.1055
127.7499
126.6254
124.0324
52.8962
35.7211
[rel]
[ppm]

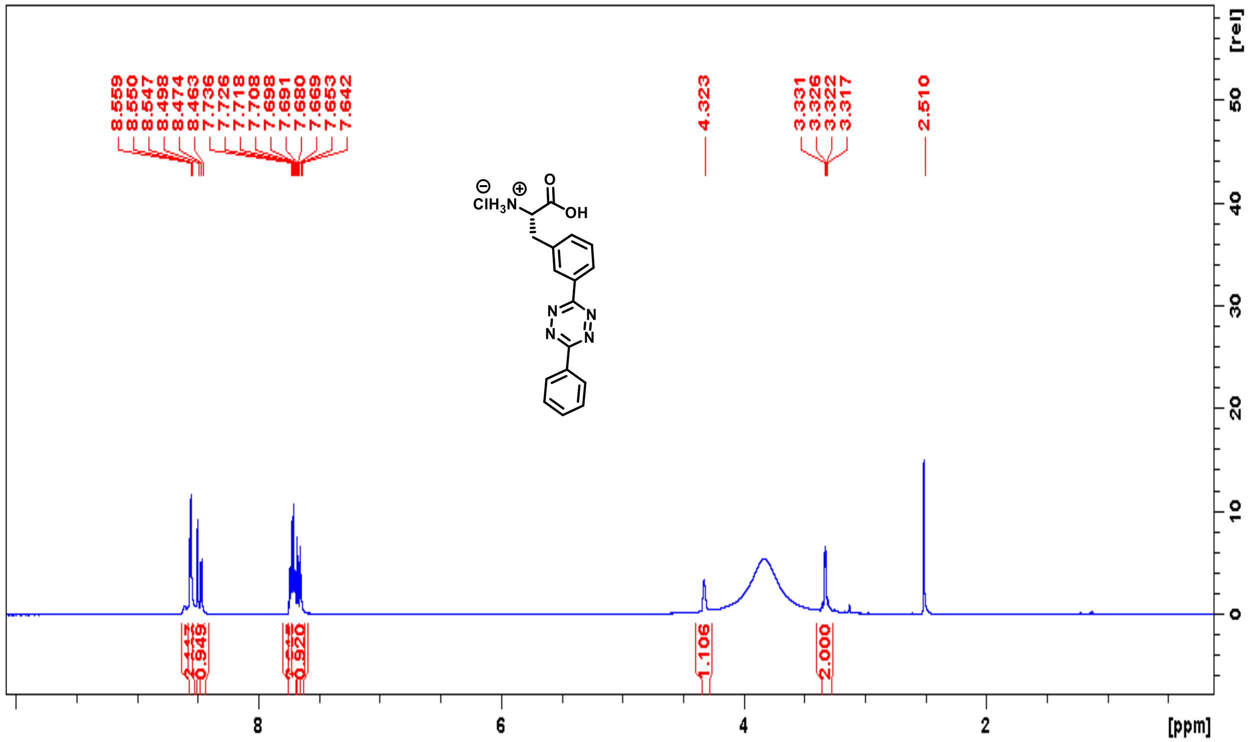
8.559
8.550
8.547
8.498
8.474
8.463
7.736
7.726
7.718
7.708
7.698
7.691
7.680
7.669
7.653
7.642
4.323
3.331
3.326
3.322
3.317
2.510
0.949
0.920
1.106
2.000
[ppm]
[rel]

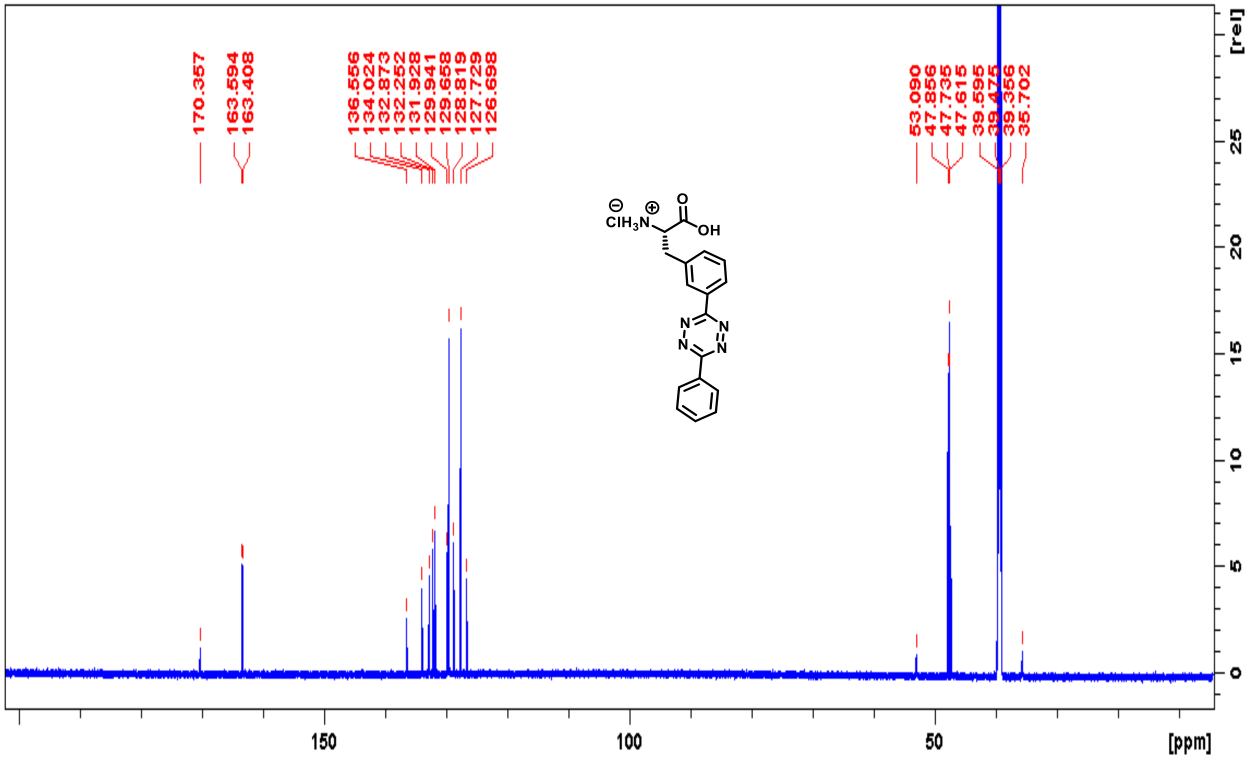
170.357
163.594
163.408
136.556
134.024
132.873
132.252
131.928
129.941
129.658
128.819
127.729
126.698
53.090
47.856
47.735
47.615
39.595
39.475
39.356
35.702
[ppm]
[rel]

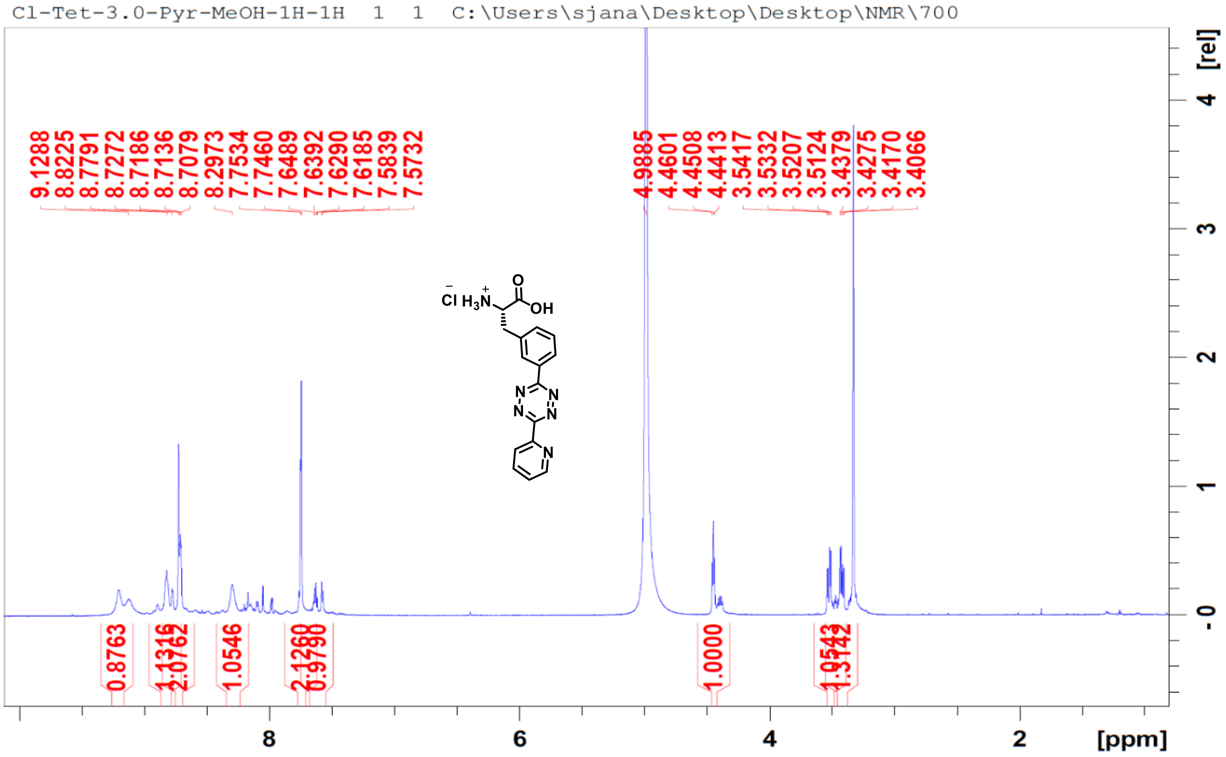
Cl-Tet-3.0-Pyr-MeOH-1H-1H 1 1 C:\Users\sjana\Desktop\Desktop\NMR\700
9.1288
8.8225
8.7791
8.7272
8.7186
8.7136
8.7079
8.2973
7.7534
7.7460
7.6489
7.6392
7.6290
7.6185
7.5839
7.5732
4.9885
4.4601
4.4508
4.4413
3.5417
3.5332
3.5207
3.5124
3.4379
3.4275
3.4170
3.4066
0.8763
1.0546
1.0000
8
6
4
2
[ppm]
[rel]
4
3
2
1
0

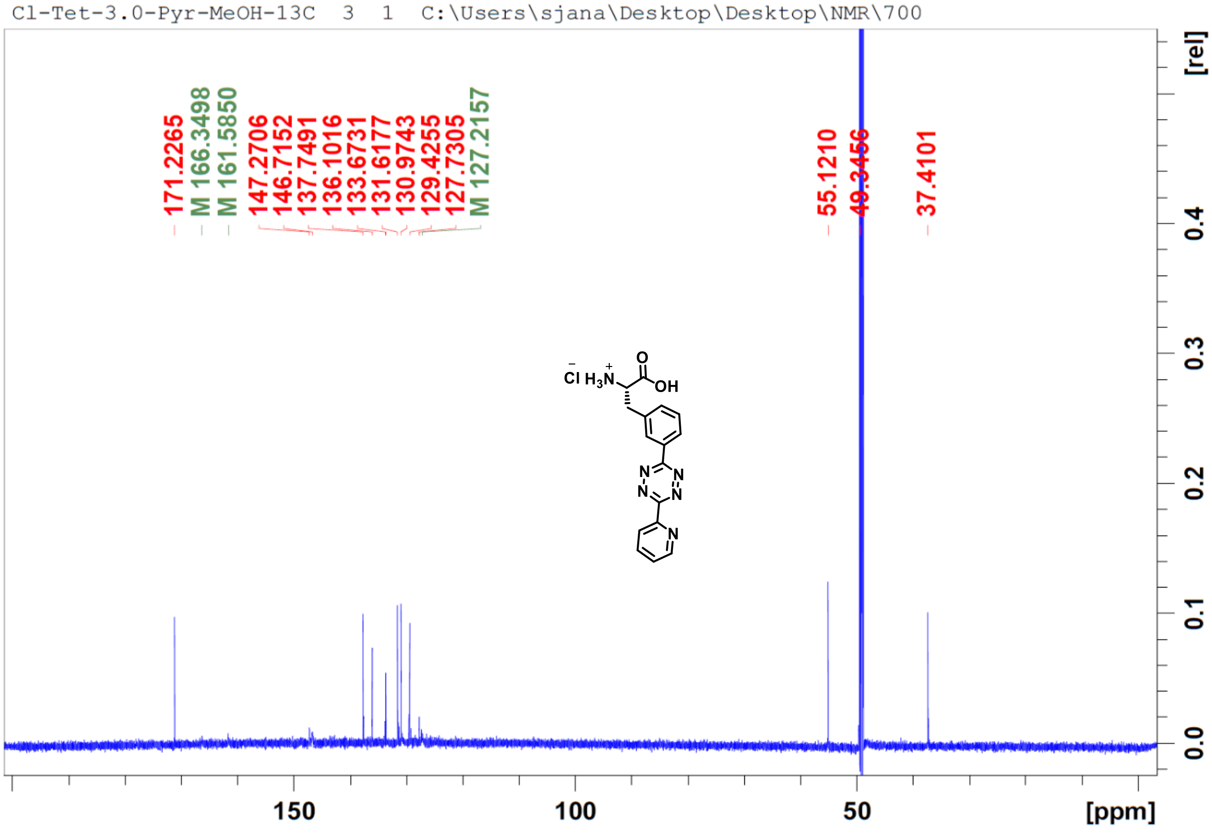
Cl-Tet-3.0-Pyr-MeOH-13C 3 1 C:\Users\sjana\Desktop\Desktop\NMR\700
171.2265
M 166.3498
M 161.5850
147.2706
146.7152
137.7491
136.1016
133.6731
131.6177
130.9743
129.4255
127.7305
M 127.2157
55.1210
49.3456
37.4101
150
100
50
[ppm]
[rel]
0.4
0.3
0.2
0.1
0.0

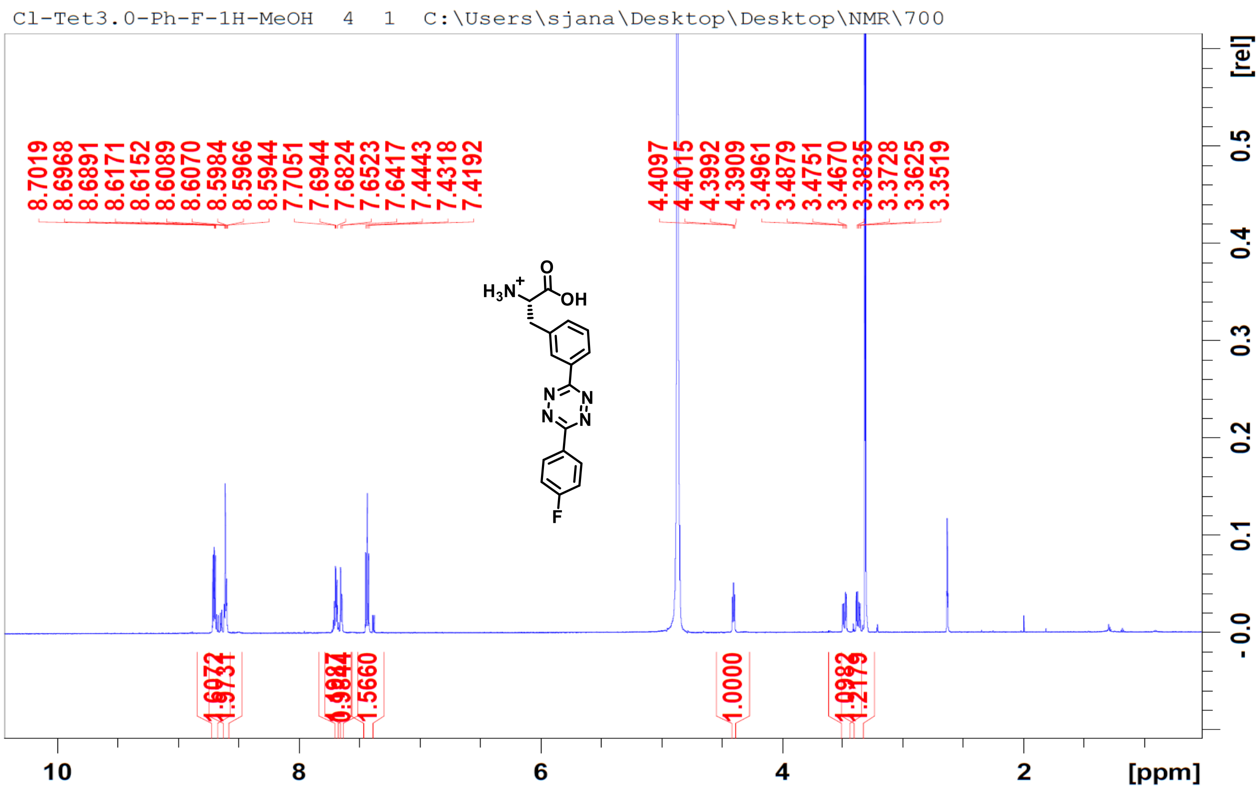
Cl-Tet3.0-Ph-F-1H-MeOH 4 1 C:\Users\sjana\Desktop\Desktop\NMR\700
8.7019
8.6968
8.6891
8.6171
8.6152
8.6089
8.6070
8.5984
8.5966
8.5944
7.7051
7.6944
7.6824
7.6523
7.6417
7.4443
7.4318
7.4192
4.4097
4.4015
4.3992
4.3909
3.4961
3.4879
3.4751
3.4670
3.3835
3.3728
3.3625
3.3519
1.0000
1.5660
[ppm]
[rel]
H3N+
O
OH
N
F

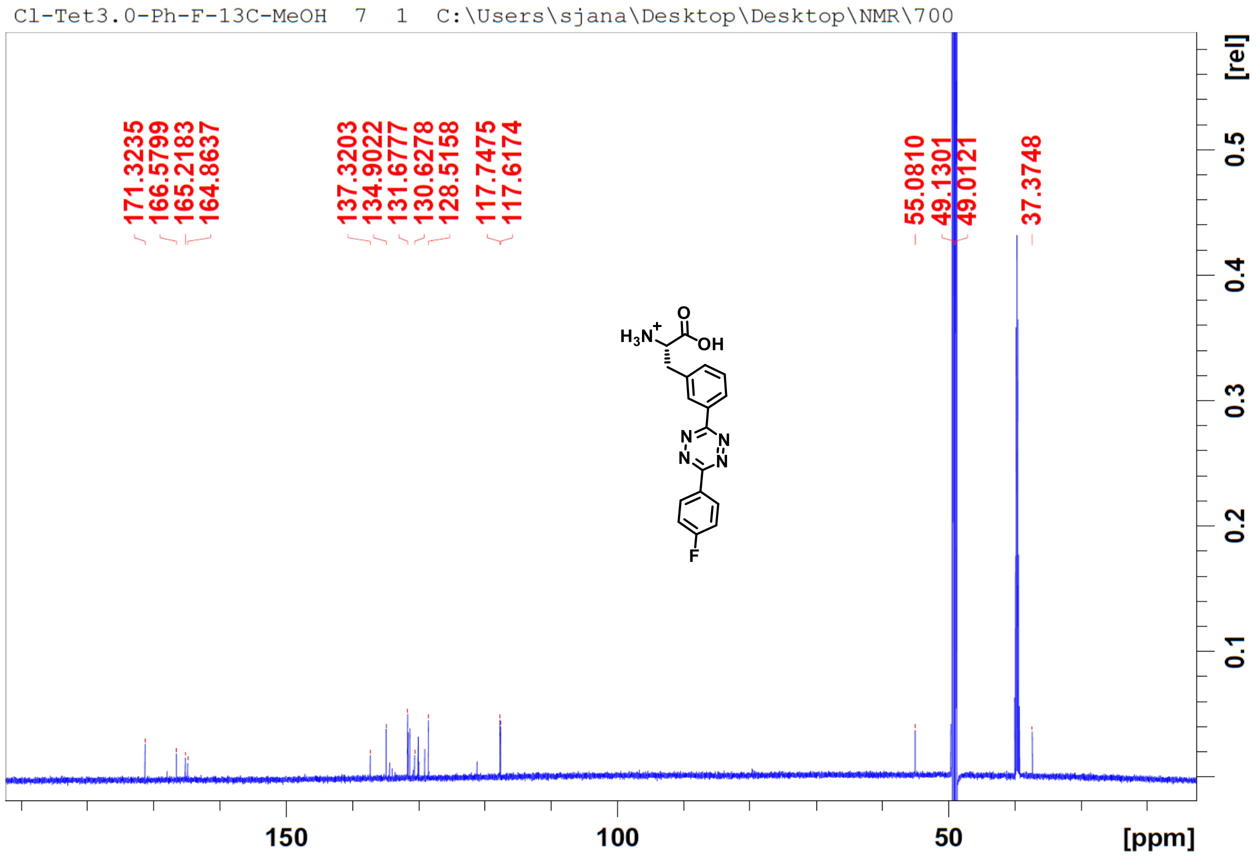
Cl-Tet3.0-Ph-F-13C-MeOH 7 1 C:\Users\sjana\Desktop\Desktop\NMR\700
171.3235
166.5799
165.2183
164.8637
137.3203
134.9022
131.6777
130.6278
128.5158
117.7475
117.6174
55.0810
49.1301
49.0121
37.3748
[ppm]
[rel]
H3N+
O
OH
N
F

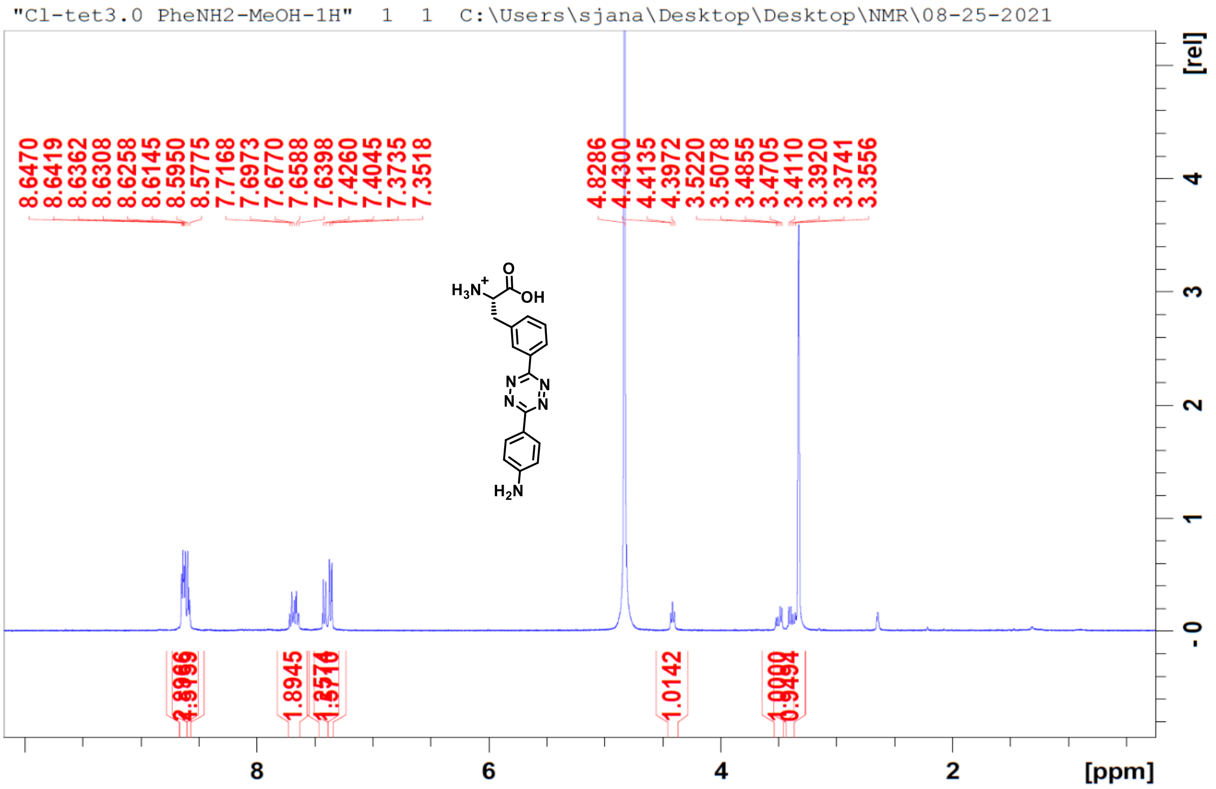
"Cl-tet3.0 PheNH2-MeOH-1H" 1 1 C:\Users\sjana\Desktop\Desktop\NMR\08-25-2021
8.6470
8.6419
8.6362
8.6308
8.6258
8.6145
8.5950
8.5775
7.7168
7.6973
7.6770
7.6588
7.6398
7.4260
7.4045
7.3735
7.3518
4.8286
4.4300
4.4135
4.3972
3.5220
3.5078
3.4855
3.4705
3.4110
3.3920
3.3741
3.3556
1.8945
1.0142
[rel]
[ppm]

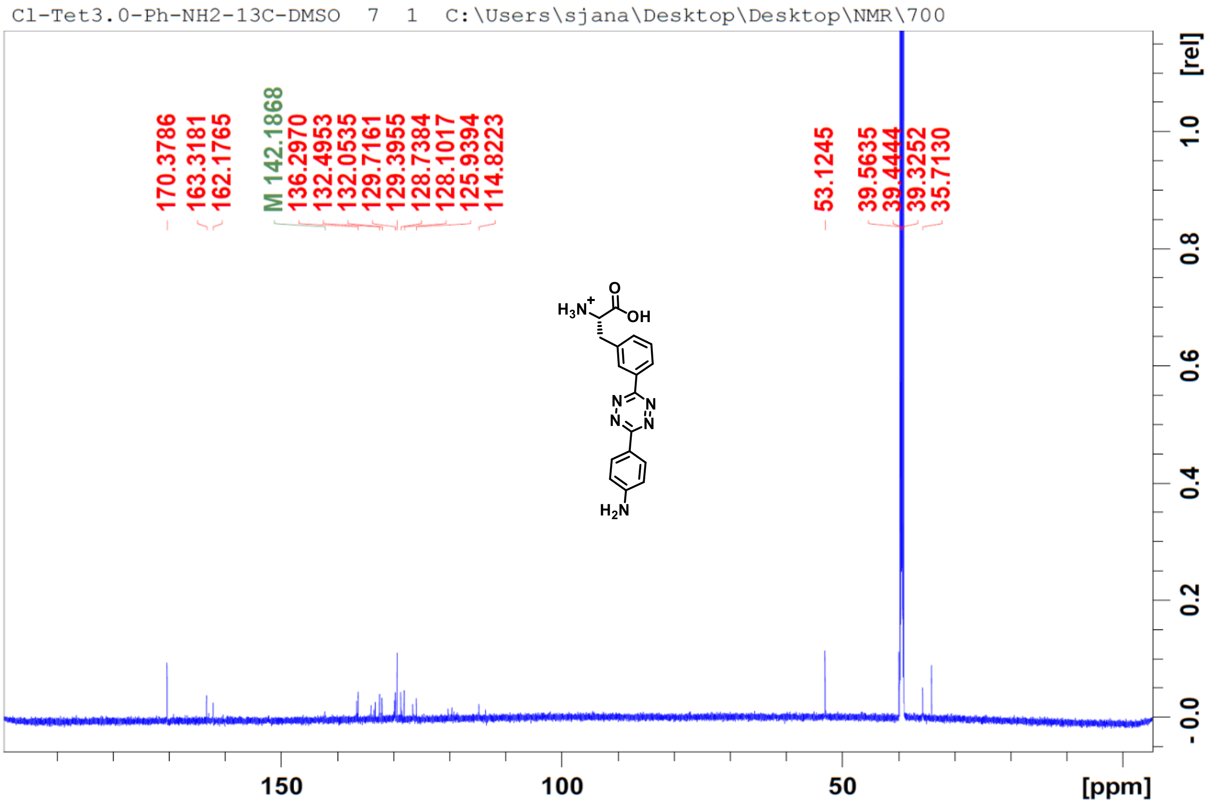
Cl-Tet3.0-Ph-NH2-13C-DMSO 7 1 C:\Users\sjana\Desktop\Desktop\NMR\700
170.3786
163.3181
162.1765
M 142.1868
136.2970
132.4953
132.0535
129.7161
129.3955
128.7384
128.1017
125.9394
114.8223
53.1245
39.5635
39.4444
39.3252
35.7130
[rel]
[ppm]

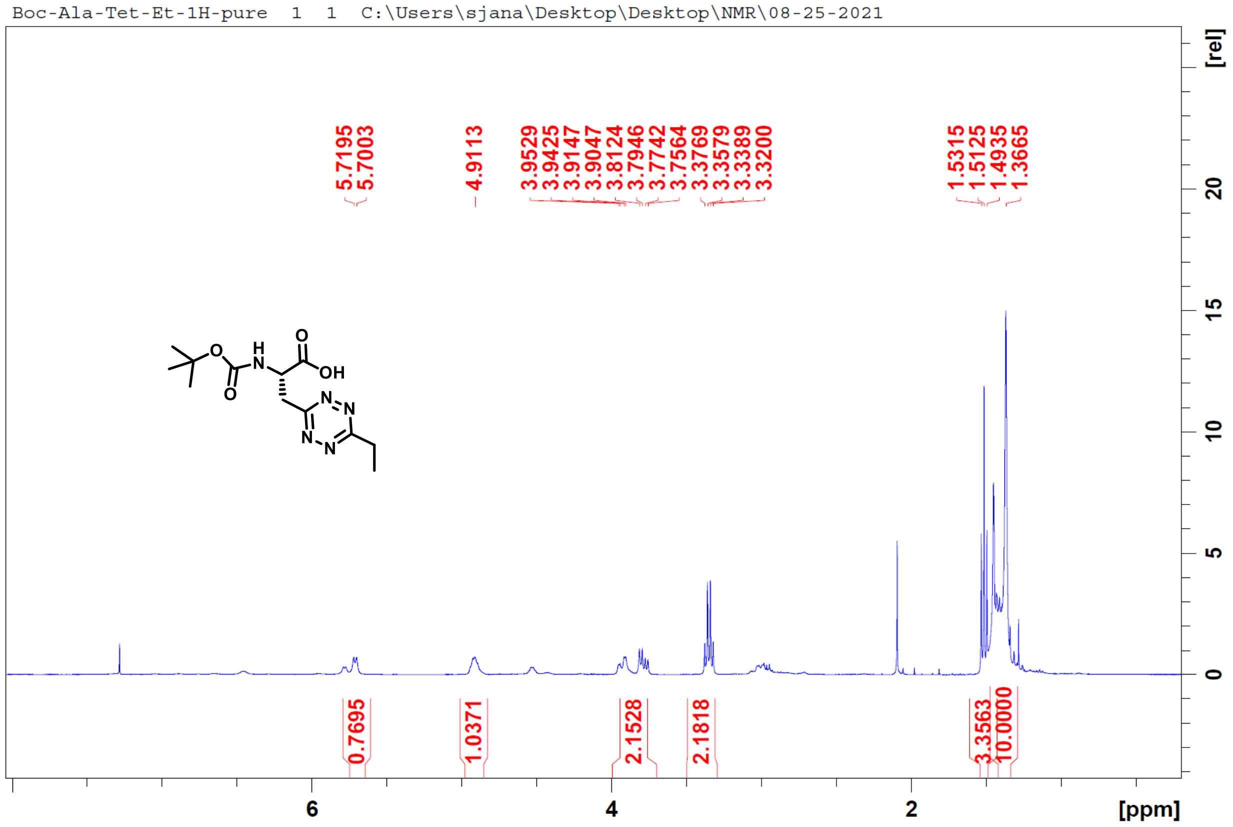
Boc-Ala-Tet-Et-1H-pure 1 1 C:\Users\sjana\Desktop\Desktop\NMR\08-25-2021
5.7195
5.7003
4.9113
3.9529
3.9425
3.9147
3.9047
3.8124
3.7946
3.7742
3.7564
3.3769
3.3579
3.3389
3.3200
1.5315
1.5125
1.4935
1.3665
0.7695
1.0371
2.1528
2.1818
3.3563
10.0000
[rel]
[ppm]

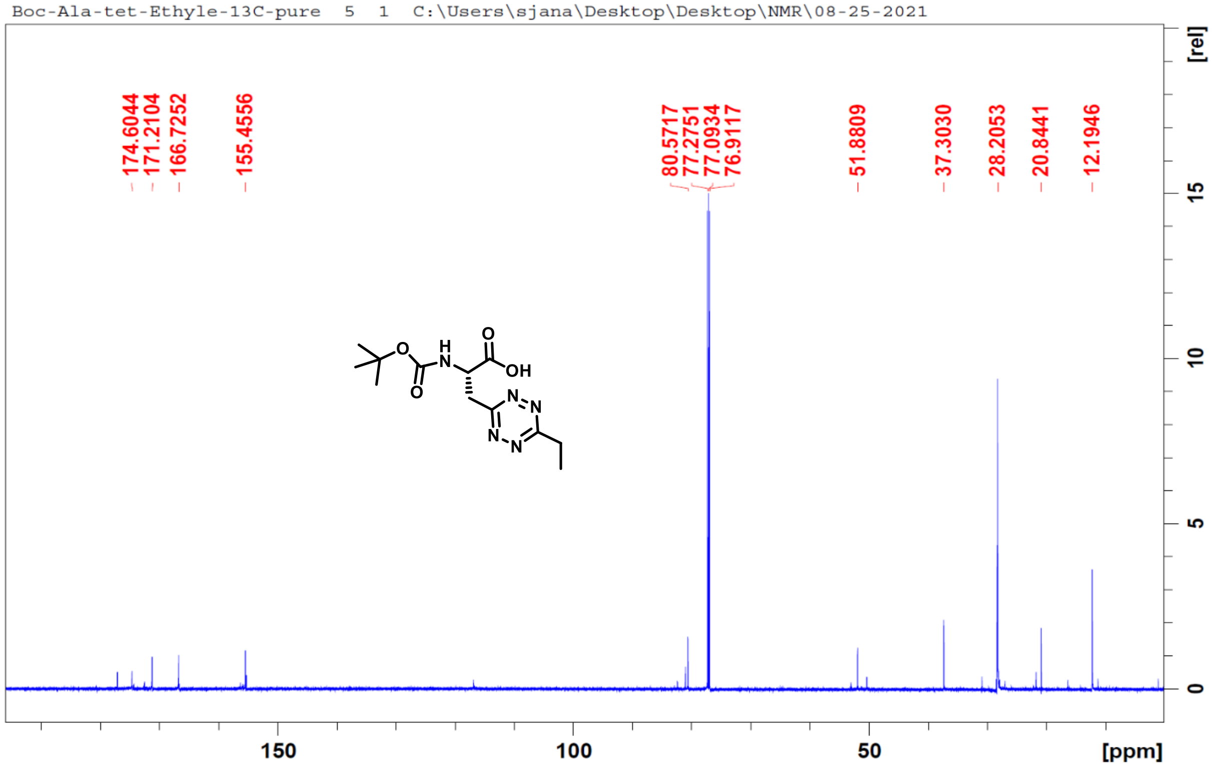
Boc-Ala-tet-Ethyle-13C-pure 5 1 C:\Users\sjana\Desktop\Desktop\NMR\08-25-2021
174.6044
171.2104
166.7252
155.4556
80.5717
77.2751
77.0934
76.9117
51.8809
37.3030
28.2053
20.8441
12.1946
[rel]
[ppm]

Boc-Ala-Tet-Isoprop-1H-pure 1 1 C:\Users\sjana\Desktop\Desktop\NMR\08-25-2021

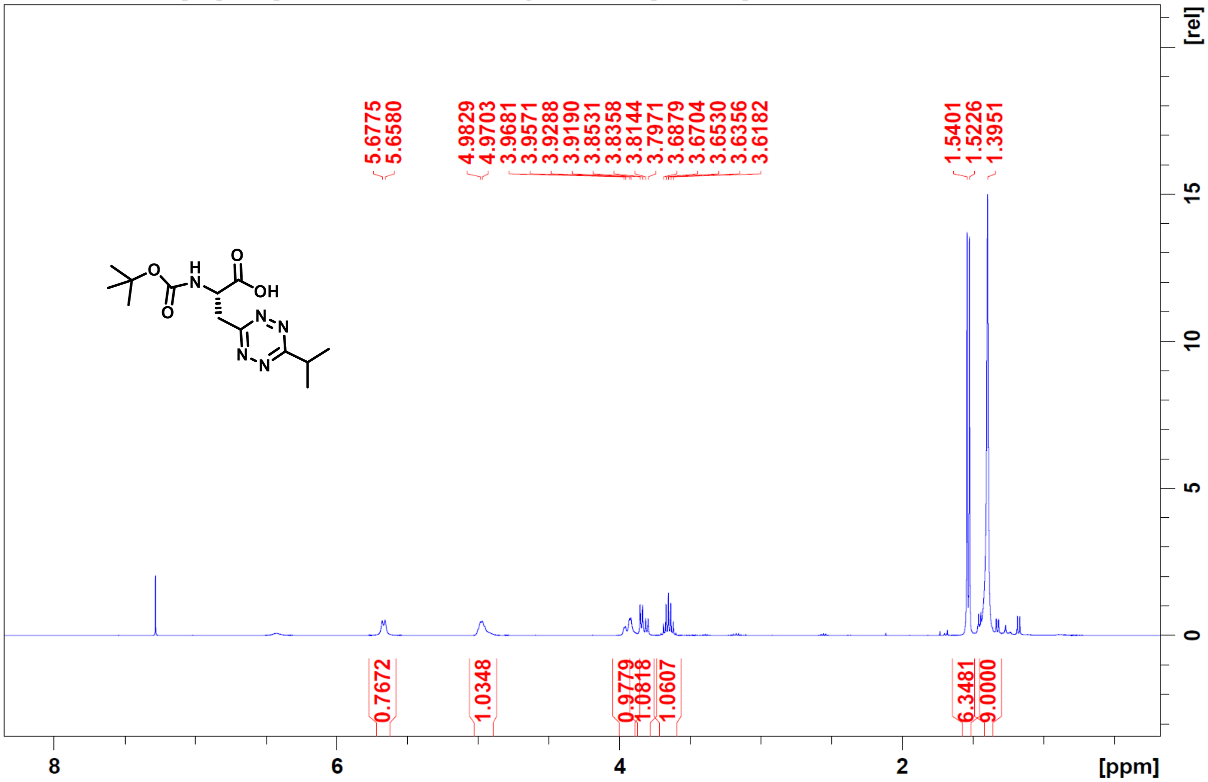

Boc-Ala-tet-Isoprop-13C-pure 4 1 C:\Users\sjana\Desktop\Desktop\NMR\08-25-2021

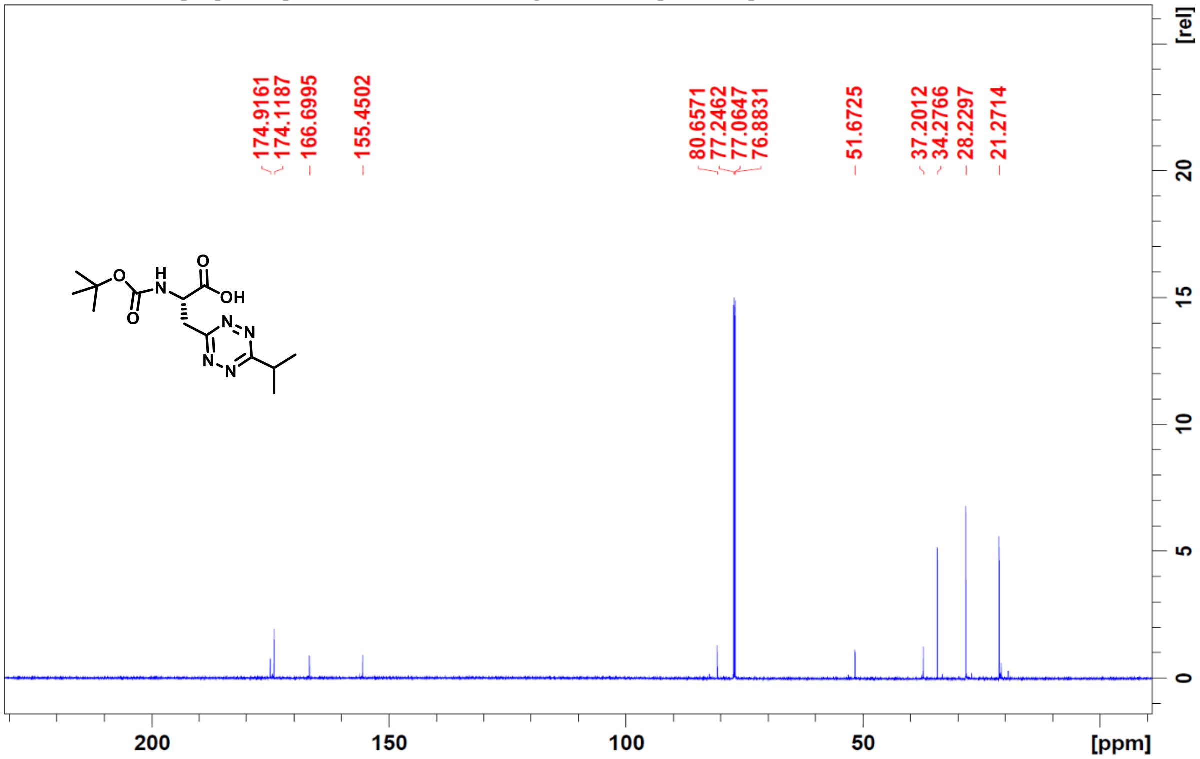

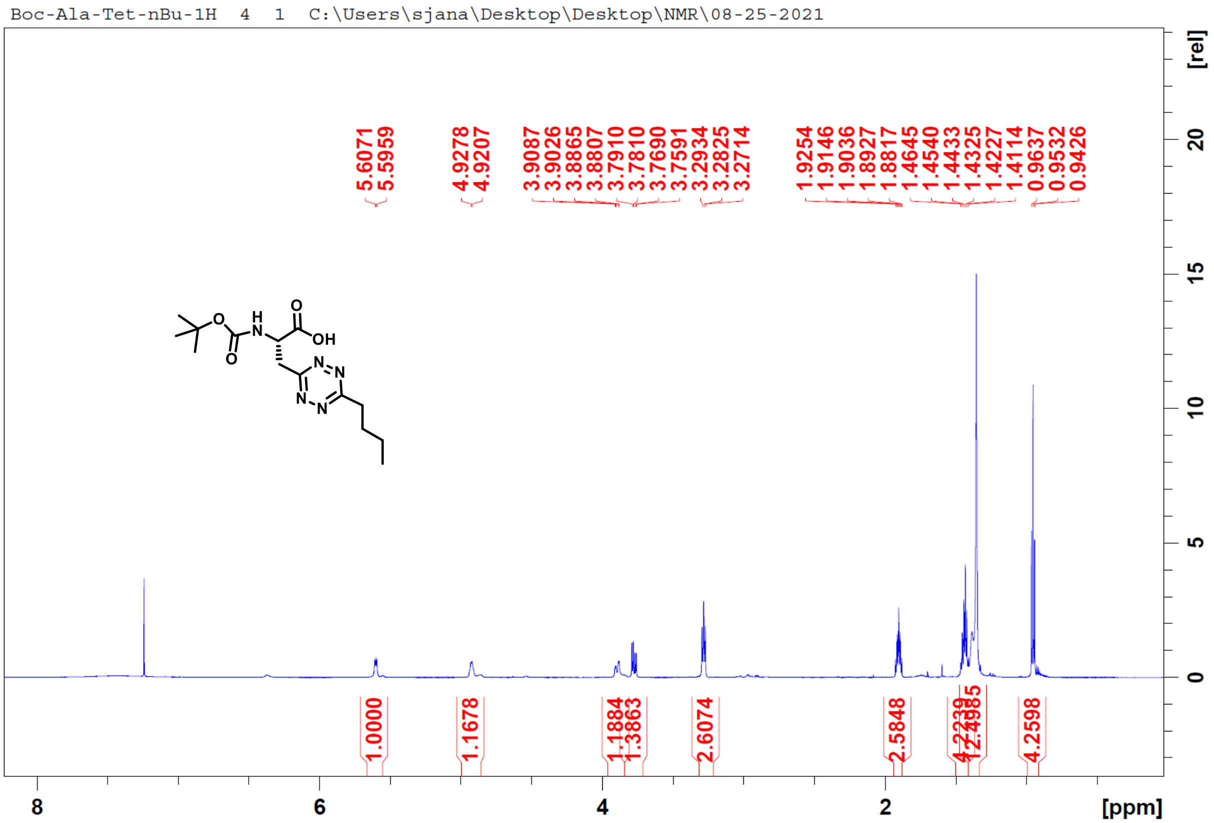
Boc-Ala-Tet-nBu-1H 4 1 C:\Users\sjana\Desktop\Desktop\NMR\08-25-2021
5.6071
5.5959
4.9278
4.9207
3.9087
3.9026
3.8865
3.8807
3.7910
3.7810
3.7690
3.7591
3.2934
3.2825
3.2714
1.9254
1.9146
1.9036
1.8927
1.8817
1.4645
1.4540
1.4433
1.4325
1.4227
1.4114
0.9637
0.9532
0.9426
1.0000
1.1678
1.1884
1.3863
2.6074
2.5848
4.2239
12.4985
4.2598
[ppm]
[rel]

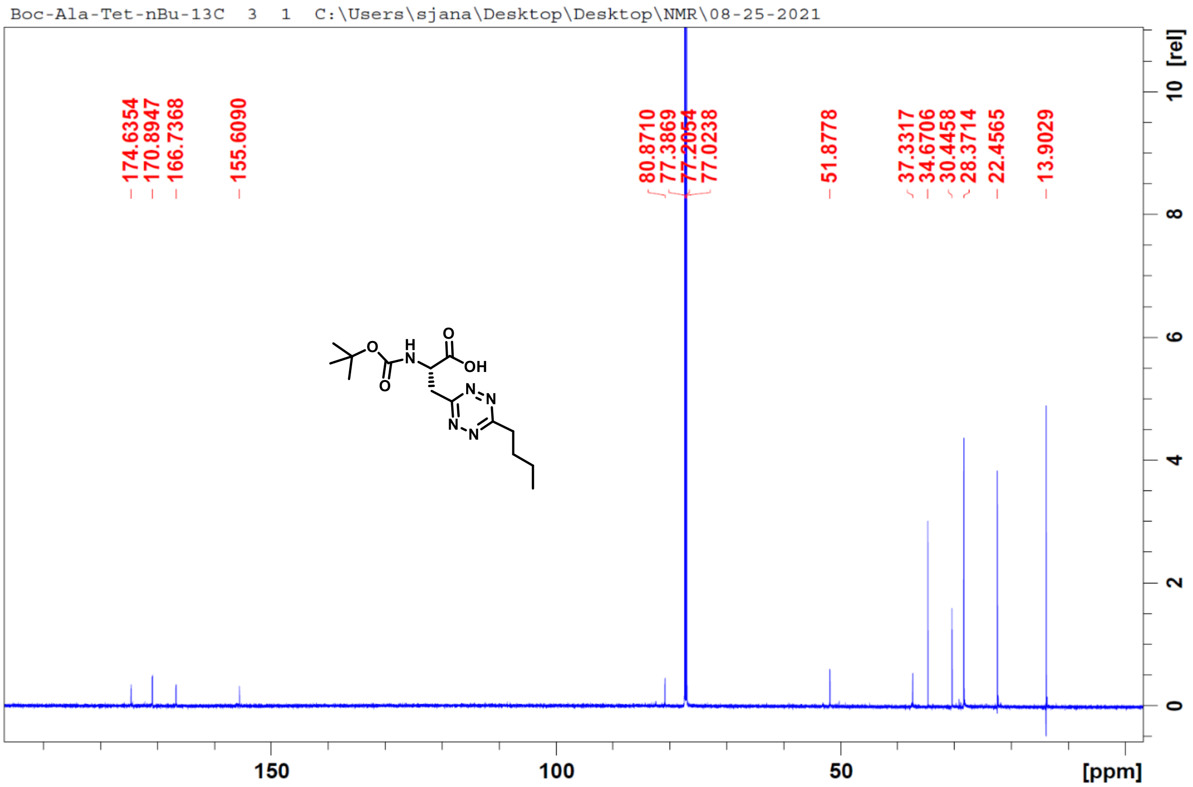
Boc-Ala-Tet-nBu-13C 3 1 C:\Users\sjana\Desktop\Desktop\NMR\08-25-2021
174.6354
170.8947
166.7368
155.6090
80.8710
77.3869
77.2054
77.0238
51.8778
37.3317
34.6706
30.4458
28.3714
22.4565
13.9029
[ppm]
[rel]

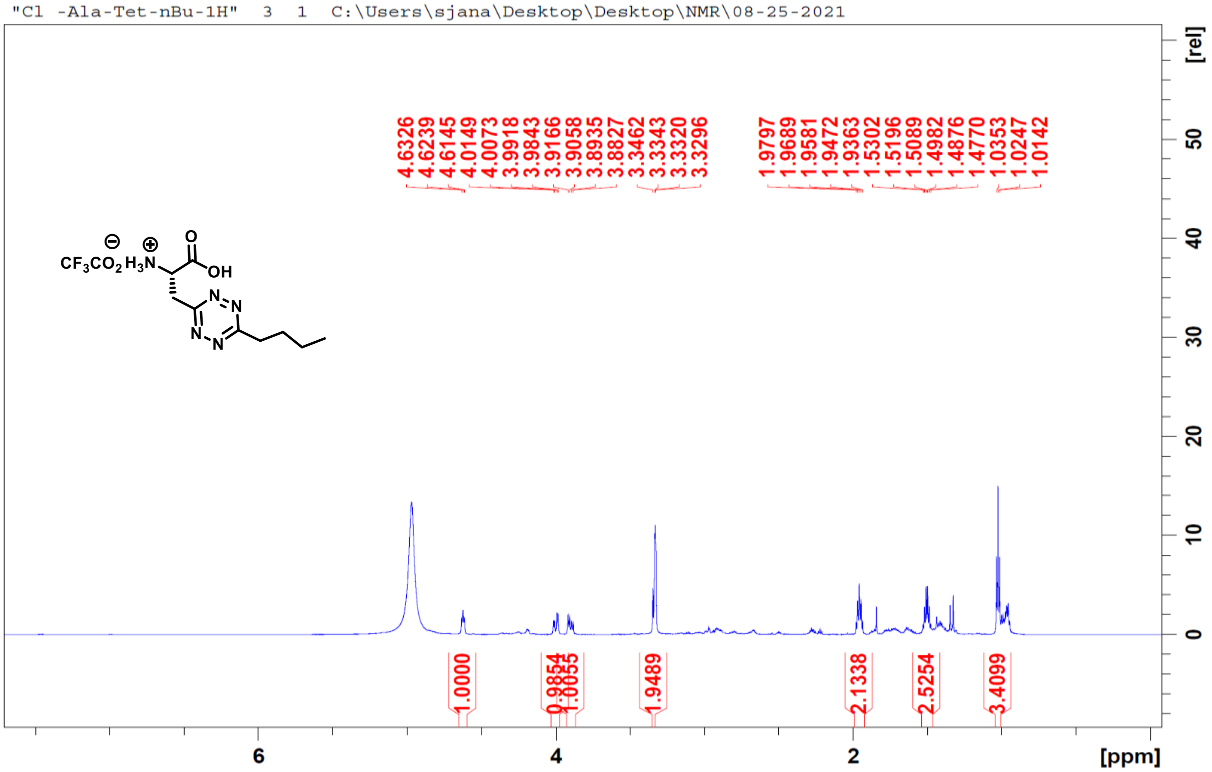
"Cl -Ala-Tet-nBu-1H" 3 1 C:\Users\sjana\Desktop\Desktop\NMR\08-25-2021
4.6326
4.6239
4.6145
4.0149
4.0073
3.9918
3.9843
3.9166
3.9058
3.8935
3.8827
3.3462
3.3343
3.3320
3.3296
1.9797
1.9689
1.9581
1.9472
1.9363
1.5302
1.5196
1.5089
1.4982
1.4876
1.4770
1.0353
1.0247
1.0142
CF3CO2 H3N
O
OH
N
N
N
N
1.0000
0.9854
1.0055
1.9489
2.1338
2.5254
3.4099
[rel]
[ppm]

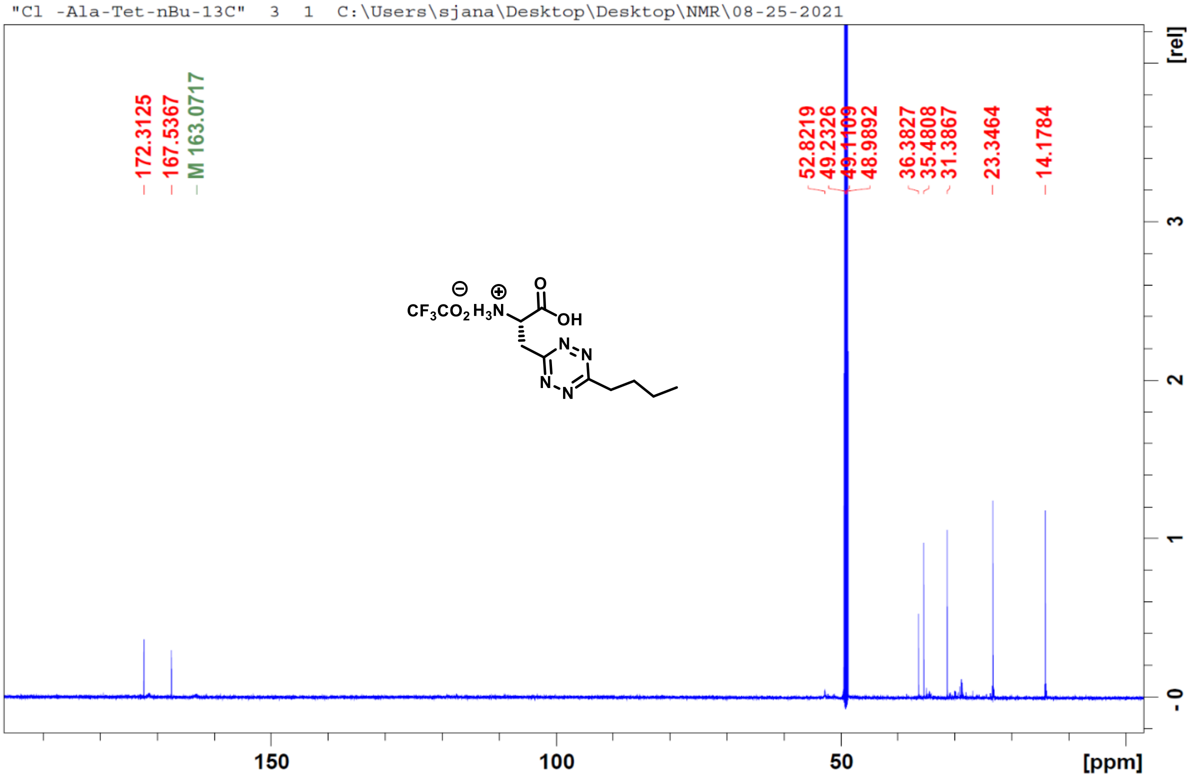
"Cl -Ala-Tet-nBu-13C" 3 1 C:\Users\sjana\Desktop\Desktop\NMR\08-25-2021
172.3125
167.5367
M 163.0717
52.8219
49.2326
49.1109
48.9892
36.3827
35.4808
31.3867
23.3464
14.1784
CF3CO2 H3N
O
OH
N
N
N
N
[rel]
[ppm]

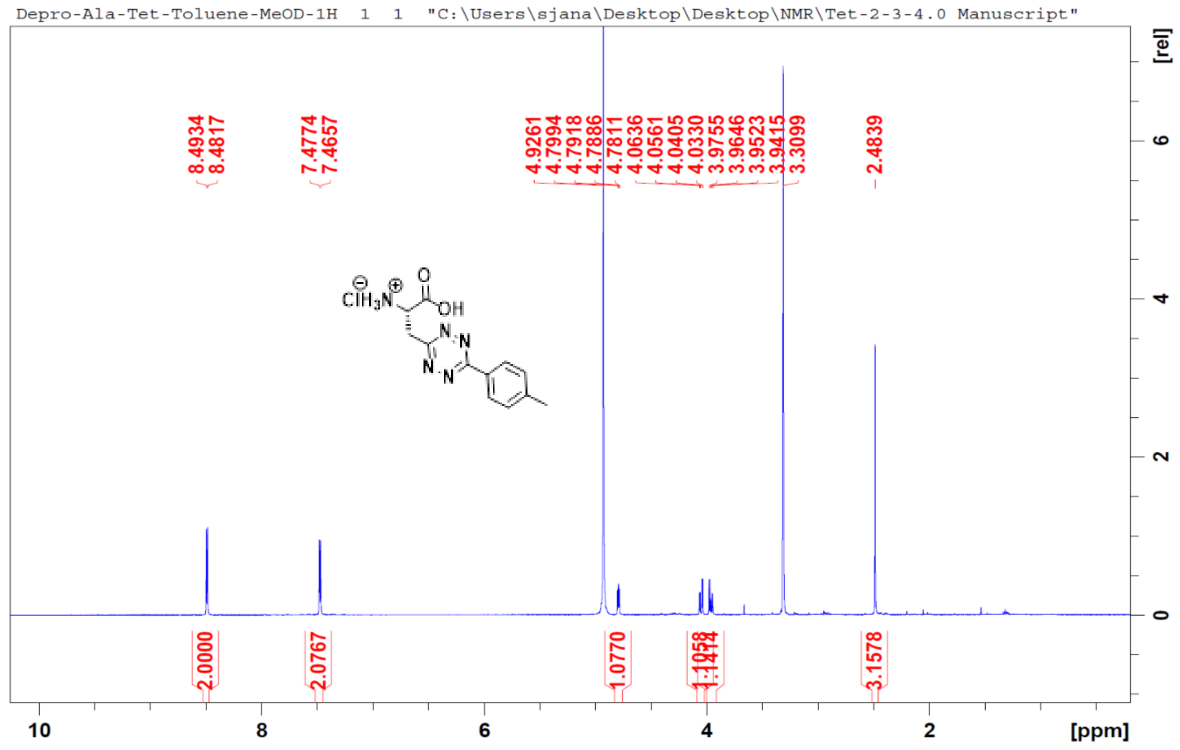
Depro-Ala-Tet-Toluene-MeOD-1H 1 1 "C:\Users\sjana\Desktop\Desktop\NMR\Tet-2-3-4.0 Manuscript"
8.4934
8.4817
7.4774
7.4657
4.9261
4.7994
4.7918
4.7886
4.7811
4.0636
4.0561
4.0405
4.0330
3.9755
3.9646
3.9523
3.9415
3.3099
2.4839
2.0000
2.0767
1.0770
3.1578
10
8
6
4
2
[ppm]
[rel]
0
2
4
6

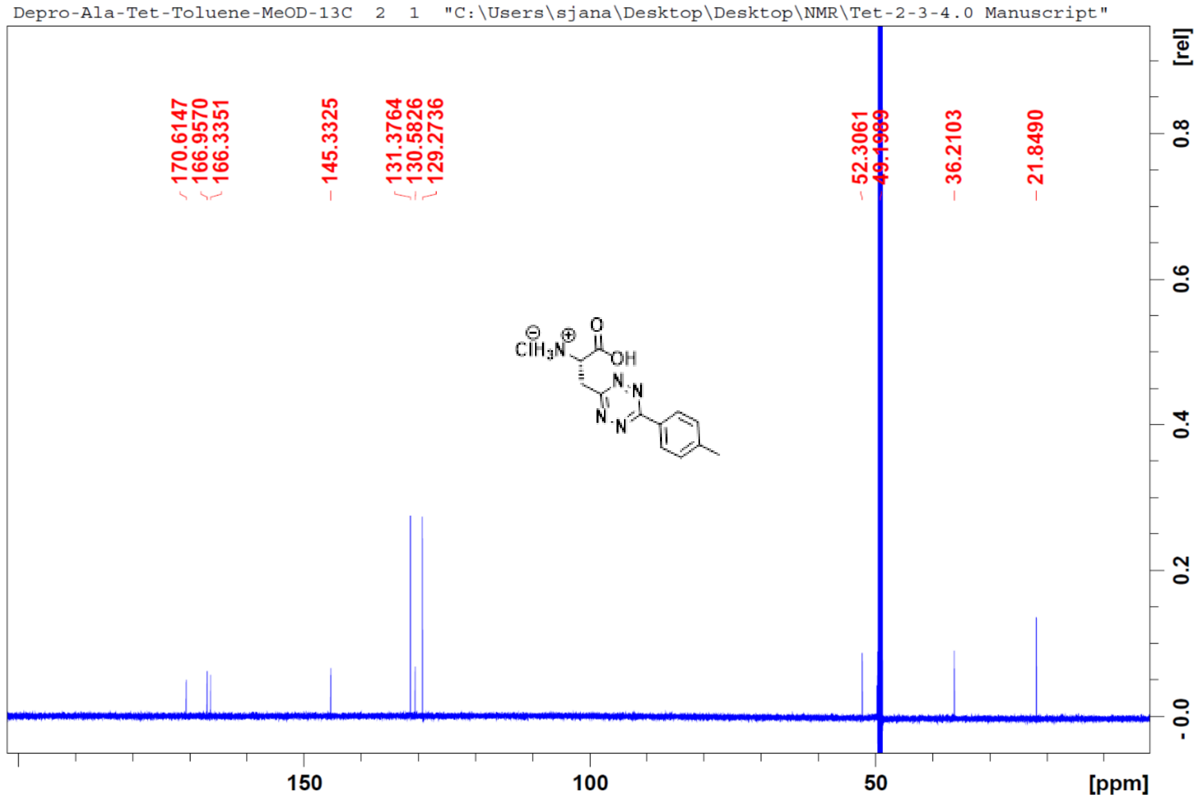
Depro-Ala-Tet-Toluene-MeOD-13C 2 1 "C:\Users\sjana\Desktop\Desktop\NMR\Tet-2-3-4.0 Manuscript"
170.6147
166.9570
166.3351
145.3325
131.3764
130.5826
129.2736
52.3061
36.2103
21.8490
150
100
50
[ppm]
[rel]
0.8
0.6
0.4
0.2
-0.0

Cl-Meta-Tet4.0-Phe-CH3-New-1H 4 1 "C:\Users\sjana\Desktop\Desktop\NMR\Tet-2-3-4.0 Manuscript"

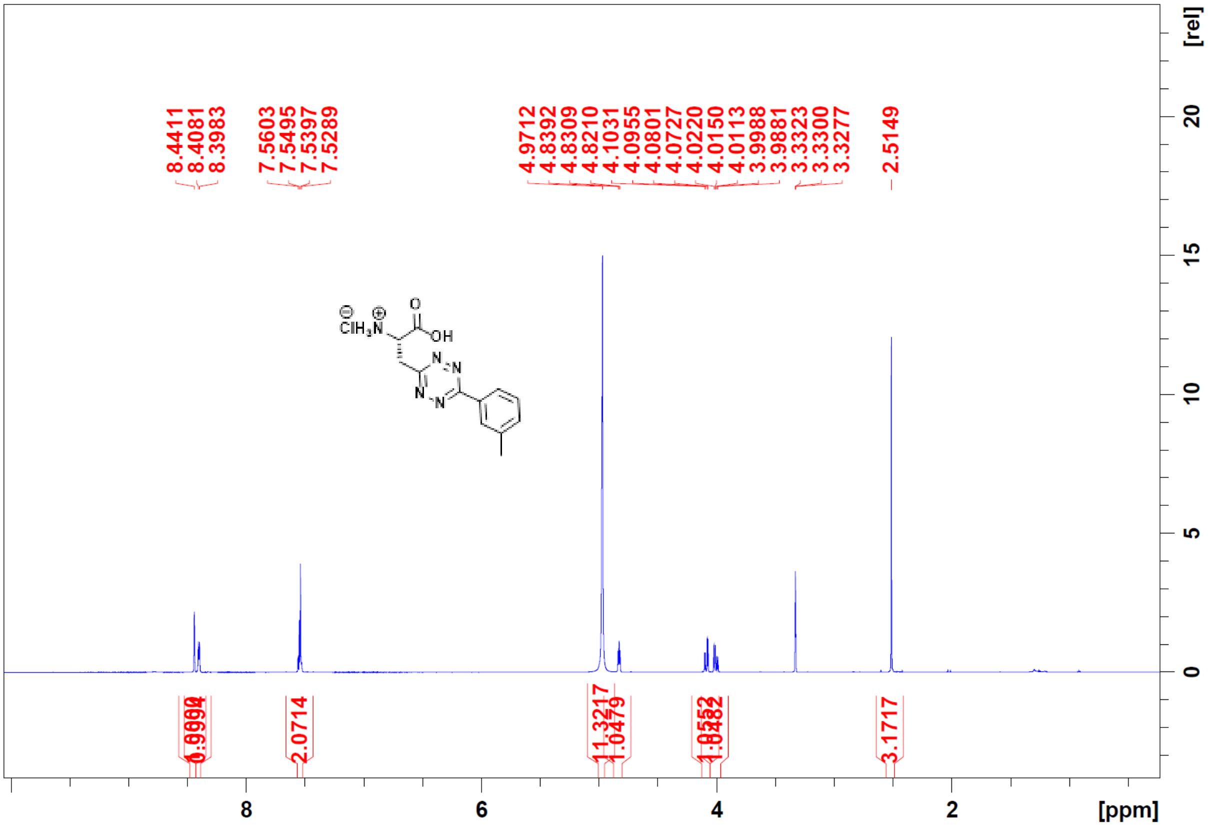

Boc-Meta-Tet4.0-Phe-CH3-New-13C 5 1 "C:\Users\sjana\Desktop\Desktop\NMR\Tet-2-3-4.0 Manuscript"

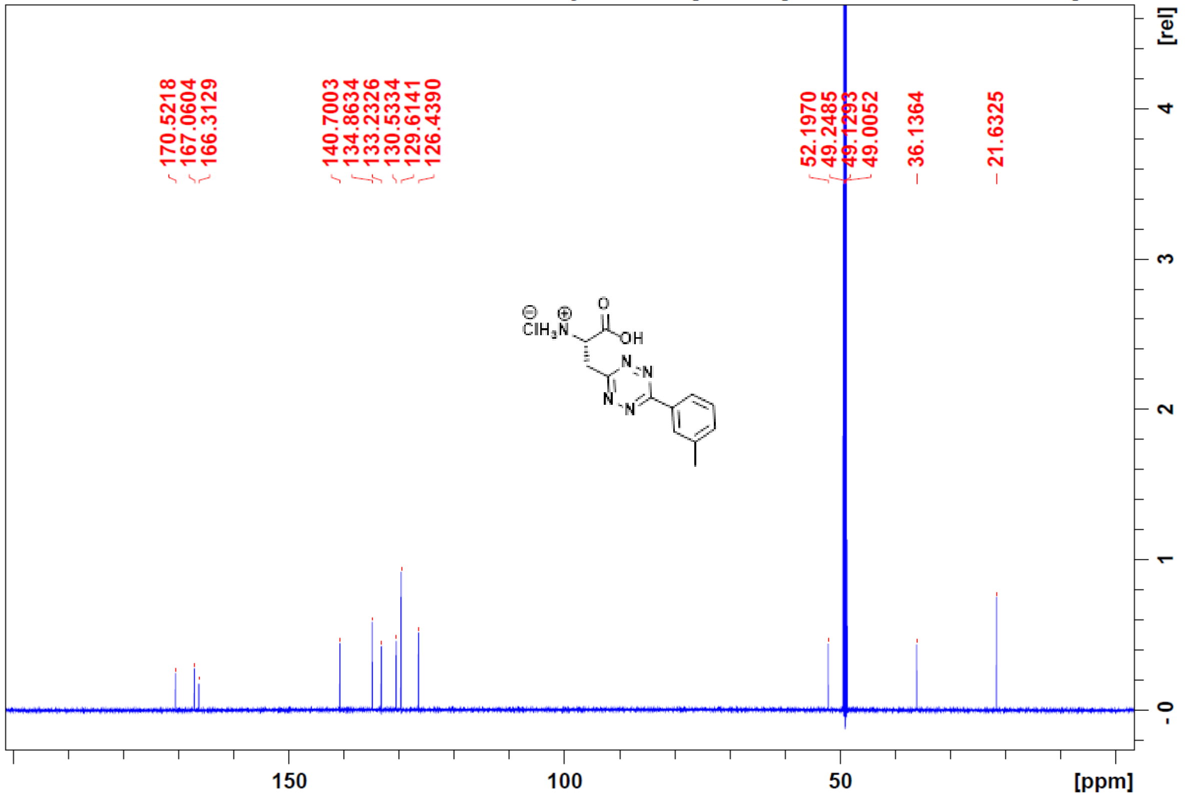

Cl-Para-Tet4.0-Phe-F-New-1H   4   1   "C:\Users\sjana\Desktop\Desktop\NMR\Tet-2-3-4.0 Manuscript"

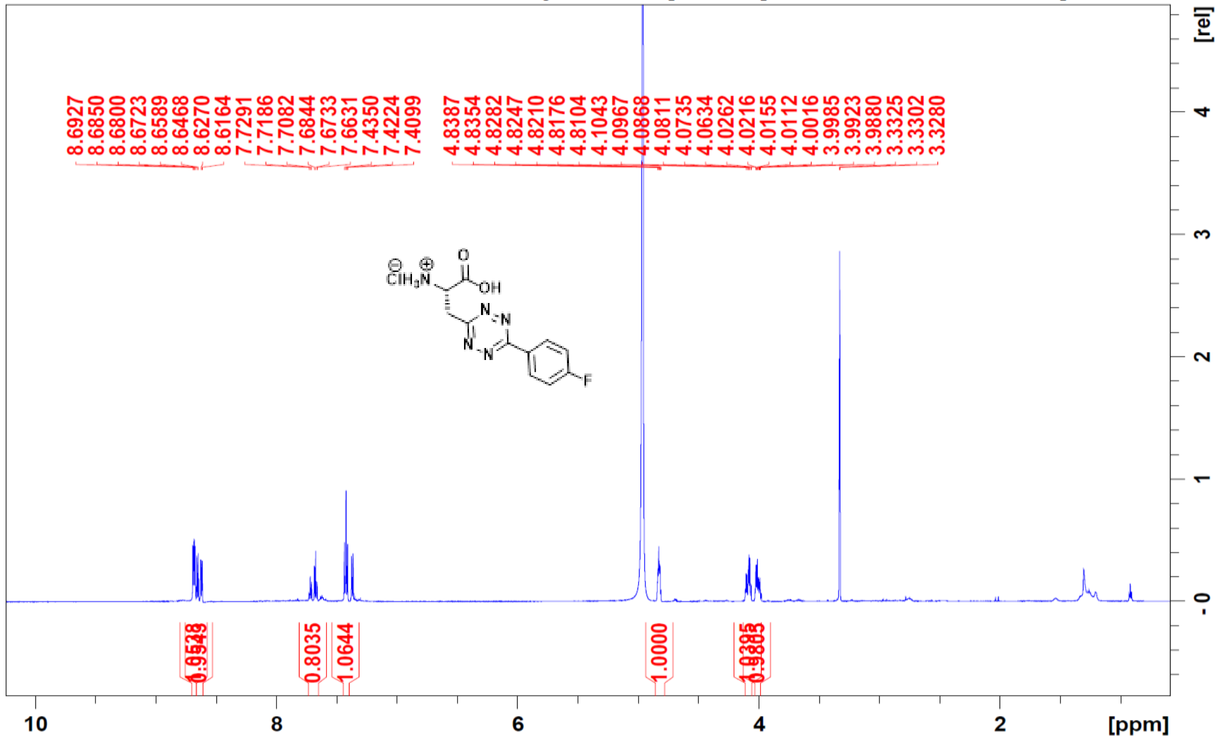

Cl-Para-Tet4.0-Phe-F-New-13C   5   1   "C:\Users\sjana\Desktop\Desktop\NMR\Tet-2-3-4.0 Manuscript"

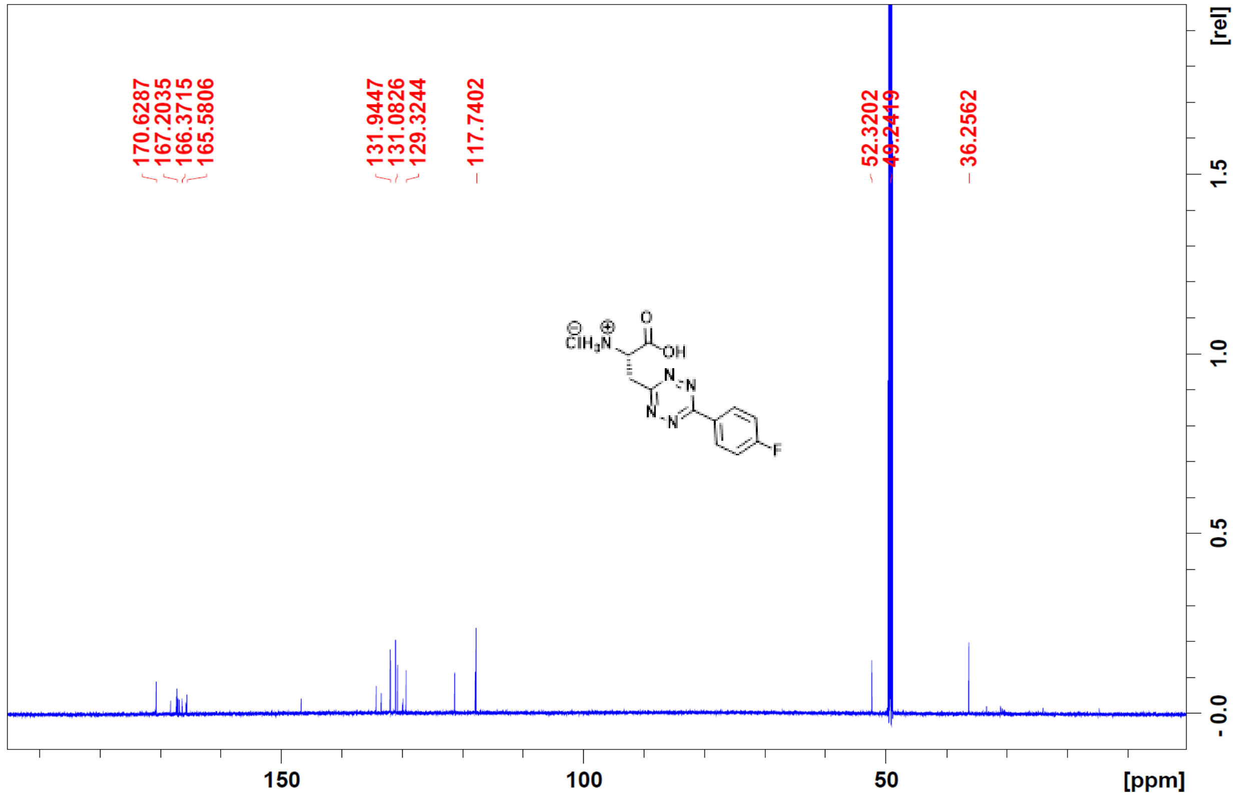

Cl-Meta-Tet4.0-Phe-F-New-1H 4 1 "C:\Users\sjana\Desktop\Desktop\NMR\Tet-2-3-4.0 Manuscript"

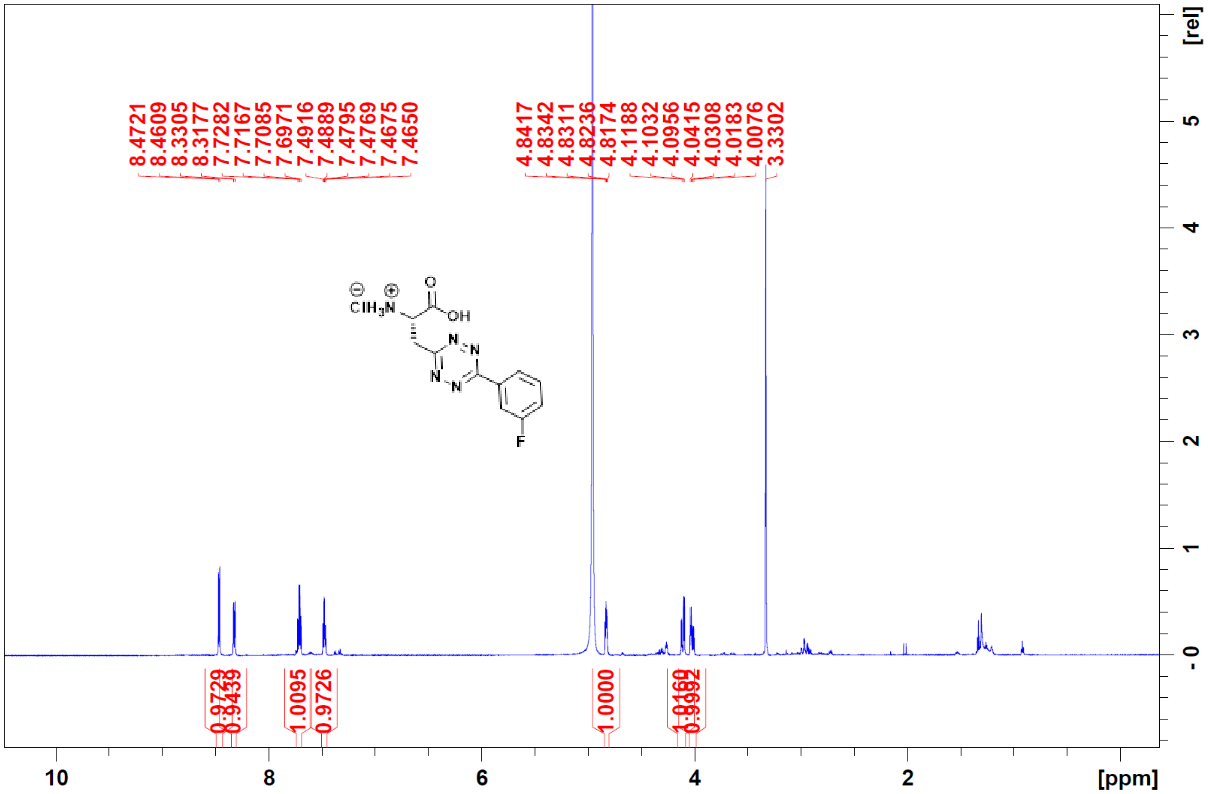

Cl-Meta-Tet4.0-Phe-F-New-13C 5 1 "C:\Users\sjana\Desktop\Desktop\NMR\Tet-2-3-4.0 Manuscript"

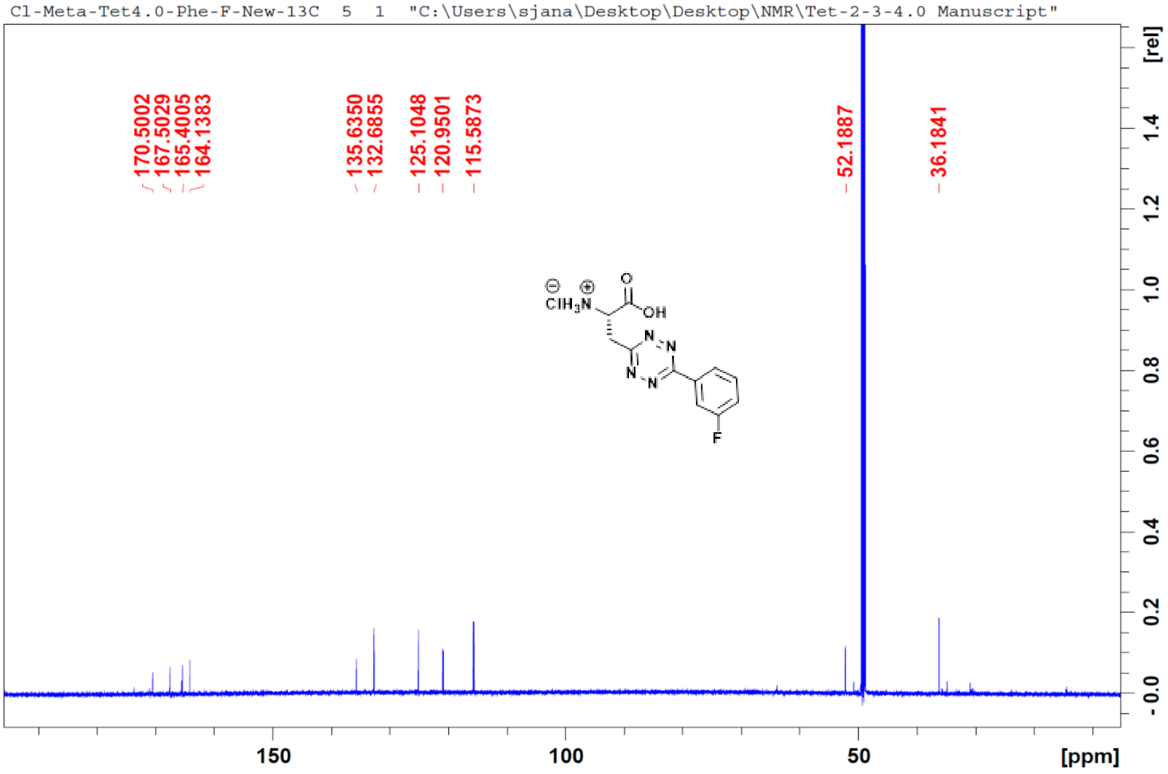

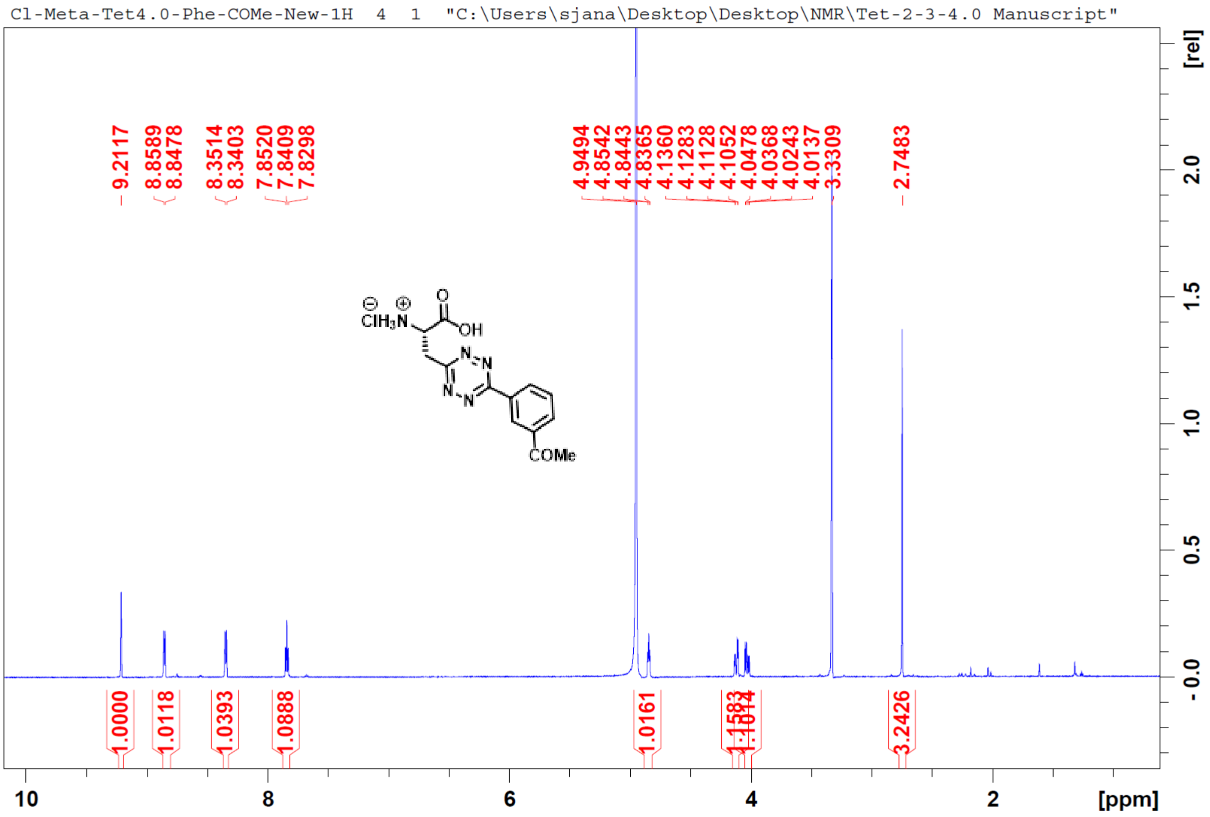
Cl-Meta-Tet4.0-Phe-COMe-New-1H 4 1 "C:\Users\sjana\Desktop\Desktop\NMR\Tet-2-3-4.0 Manuscript"
9.2117
8.8589
8.8478
8.3514
8.3403
7.8520
7.8409
7.8298
4.9494
4.8542
4.8443
4.8365
4.1360
4.1283
4.1128
4.1052
4.0478
4.0368
4.0243
4.0137
3.3309
2.7483
1.0000
1.0118
1.0393
1.0888
1.0161
3.2426
ClH3N
OH
COMe
10
8
6
4
2
[ppm]
[rel]
2.0
1.5
1.0
0.5
-0.0

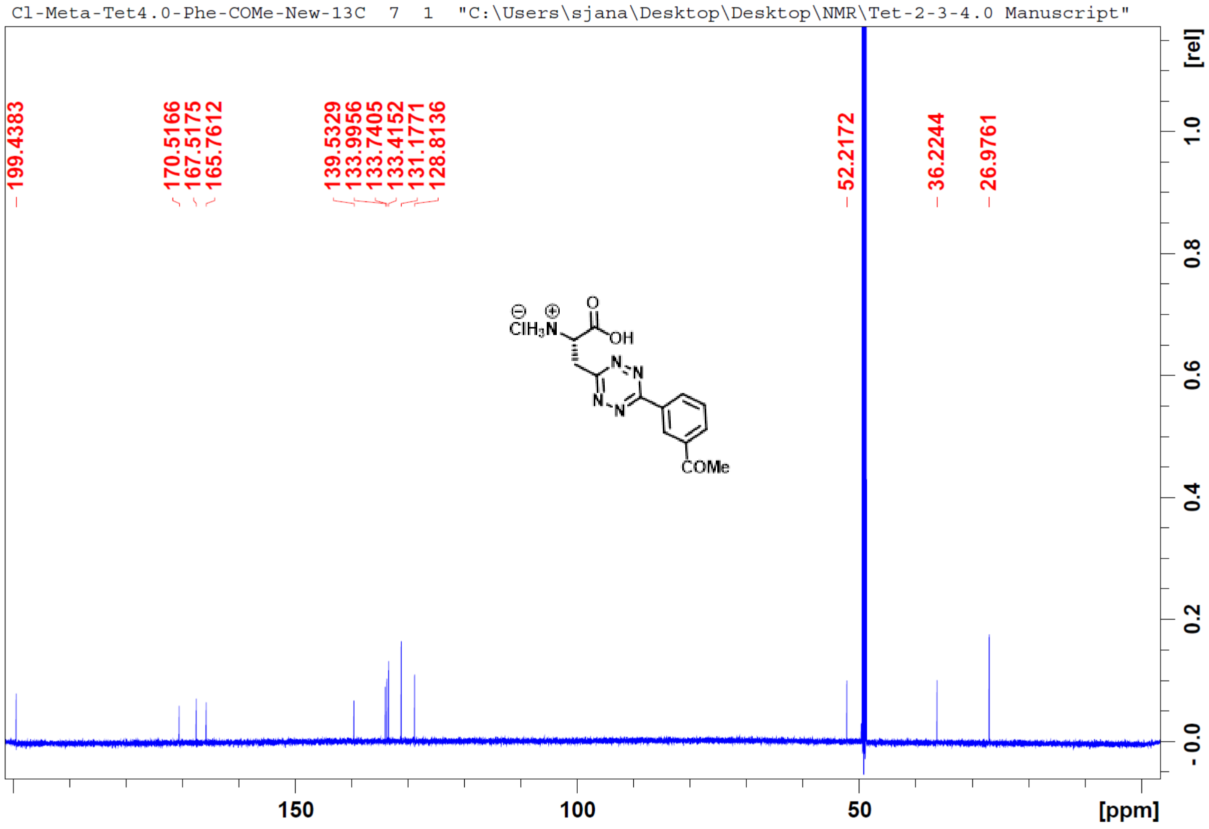
Cl-Meta-Tet4.0-Phe-COMe-New-13C 7 1 "C:\Users\sjana\Desktop\Desktop\NMR\Tet-2-3-4.0 Manuscript"
199.4383
170.5166
167.5175
165.7612
139.5329
133.9956
133.7405
133.4152
131.1771
128.8136
52.2172
36.2244
26.9761
ClH3N
OH
COMe
150
100
50
[ppm]
[rel]
1.0
0.8
0.6
0.4
0.2
-0.0

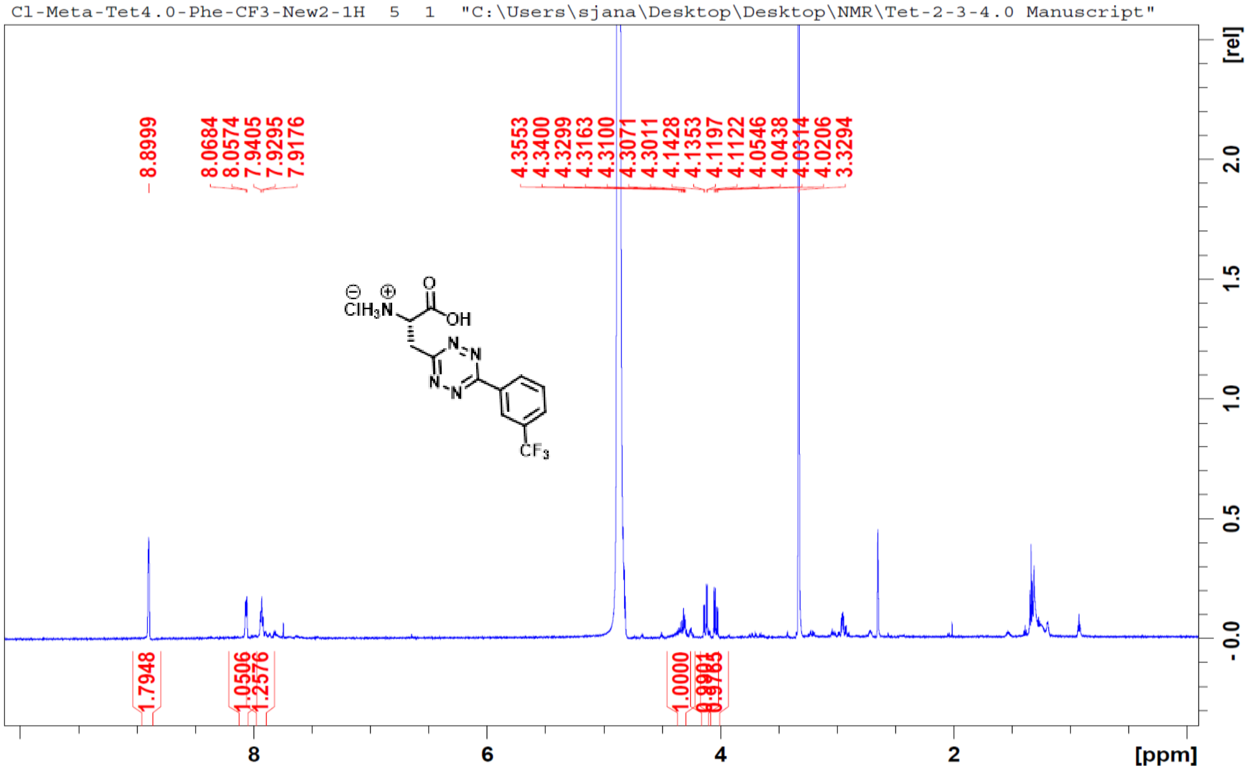
Cl-Meta-Tet4.0-Phe-CF3-New2-1H 5 1 "C:\Users\sjana\Desktop\Desktop\NMR\Tet-2-3-4.0 Manuscript"
8.8999
8.0684
8.0574
7.9405
7.9295
7.9176
4.3553
4.3400
4.3299
4.3163
4.3100
4.3071
4.3011
4.1428
4.1353
4.1197
4.1122
4.0546
4.0438
4.0314
4.0206
3.3294
1.7948
1.0506
1.2576
1.0000
[rel]
[ppm]

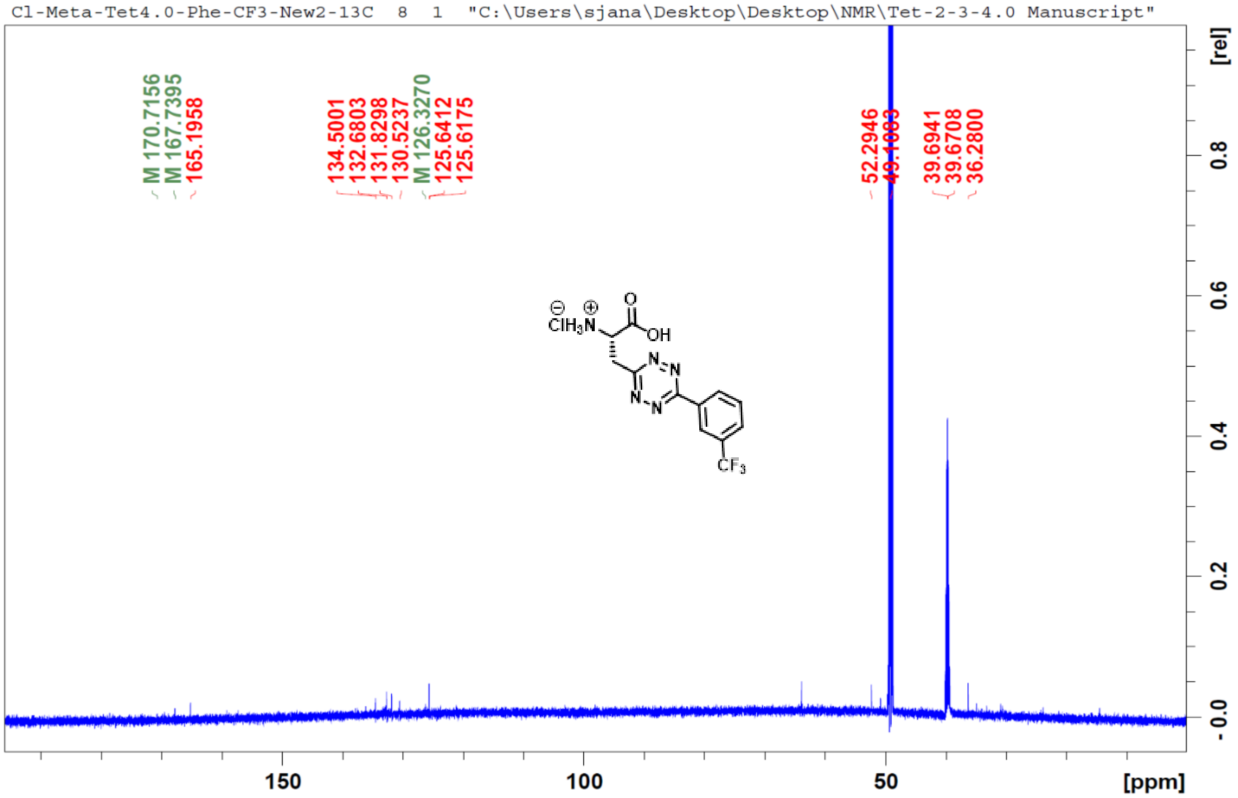
Cl-Meta-Tet4.0-Phe-CF3-New2-13C 8 1 "C:\Users\sjana\Desktop\Desktop\NMR\Tet-2-3-4.0 Manuscript"
M 170.7156
M 167.7395
165.1958
134.5001
132.6803
131.8298
130.5237
M 126.3270
125.6412
125.6175
52.2946
39.6941
39.6708
36.2800
[rel]
[ppm]

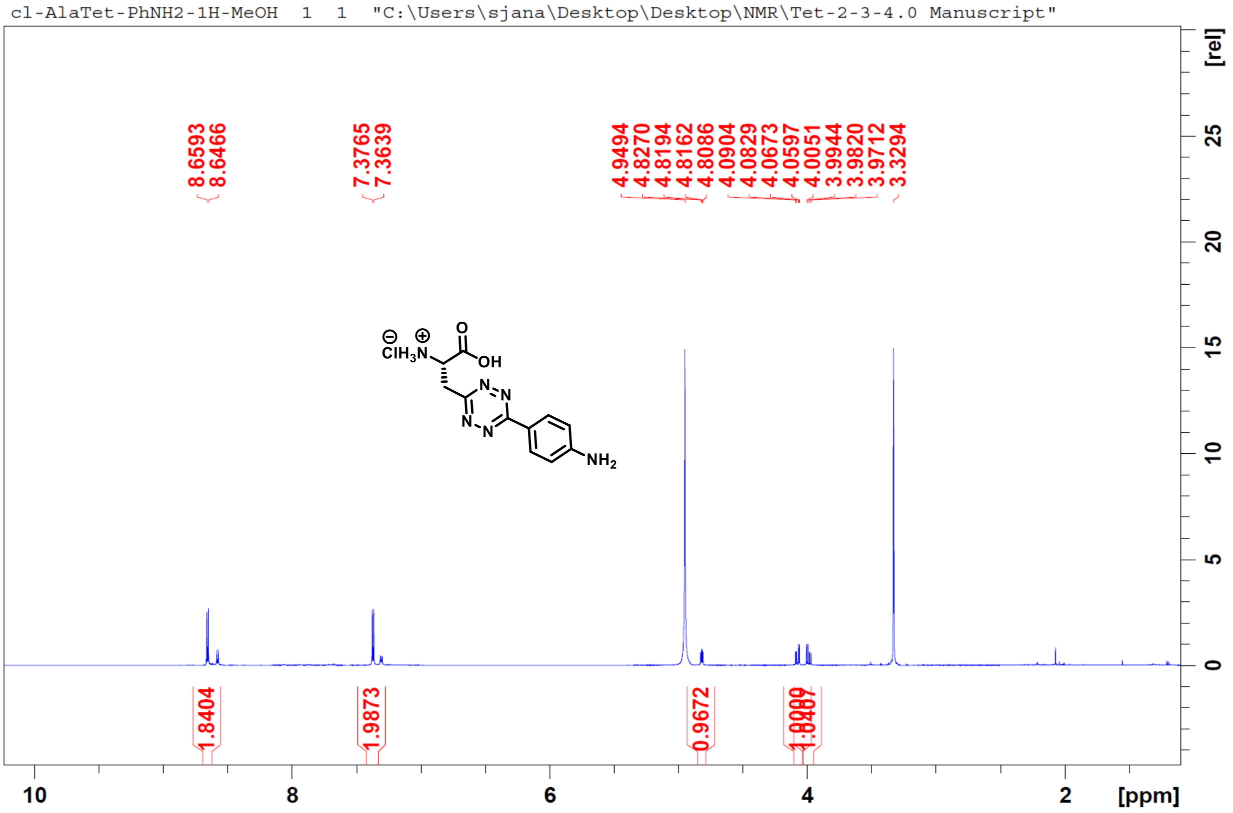
cl-AlaTet-PhNH2-1H-MeOH 1 1 "C:\Users\sjana\Desktop\Desktop\NMR\Tet-2-3-4.0 Manuscript"
8.6593
8.6466
7.3765
7.3639
4.9494
4.8270
4.8194
4.8162
4.8086
4.0904
4.0829
4.0673
4.0597
4.0051
3.9944
3.9820
3.9712
3.3294
1.8404
1.9873
0.9672
[rel]
[ppm]

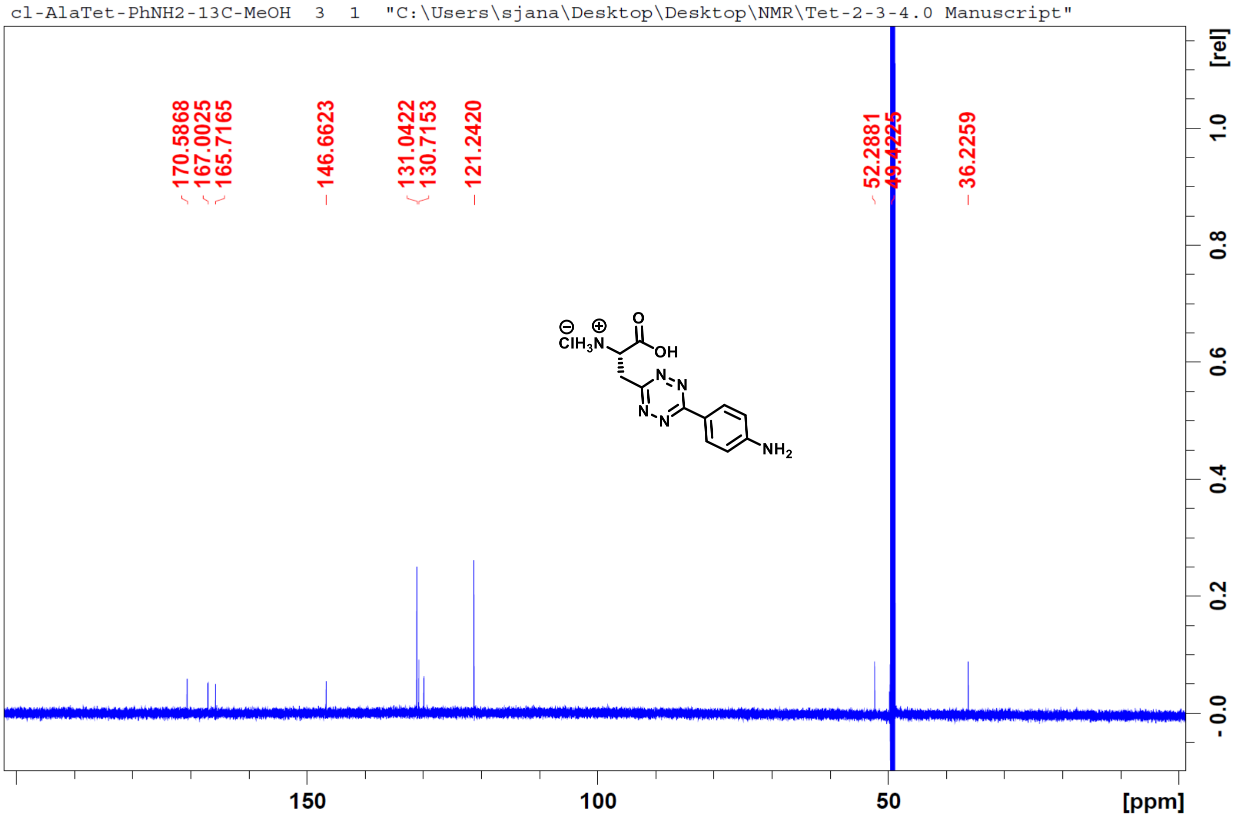
cl-AlaTet-PhNH2-13C-MeOH 3 1 "C:\Users\sjana\Desktop\Desktop\NMR\Tet-2-3-4.0 Manuscript"
170.5868
167.0025
165.7165
146.6623
131.0422
130.7153
121.2420
52.2881
36.2259
[rel]
[ppm]

cl-AlaTet-PhOH-1H-MeOH 2 1 "C:\Users\sjana\Desktop\Desktop\NMR\Tet-2-3-4.0 Manuscript"

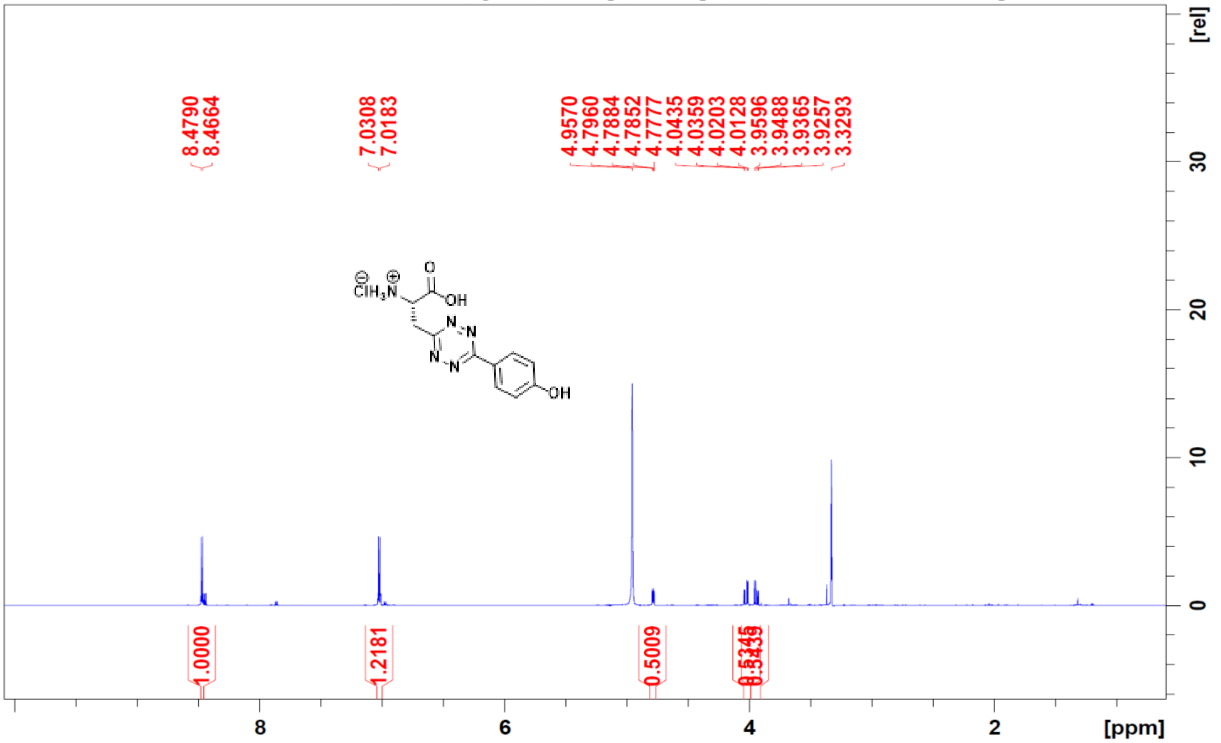

cl-AlaTet-PhOH-13C-MeOH 3 1 "C:\Users\sjana\Desktop\Desktop\NMR\Tet-2-3-4.0 Manuscript"

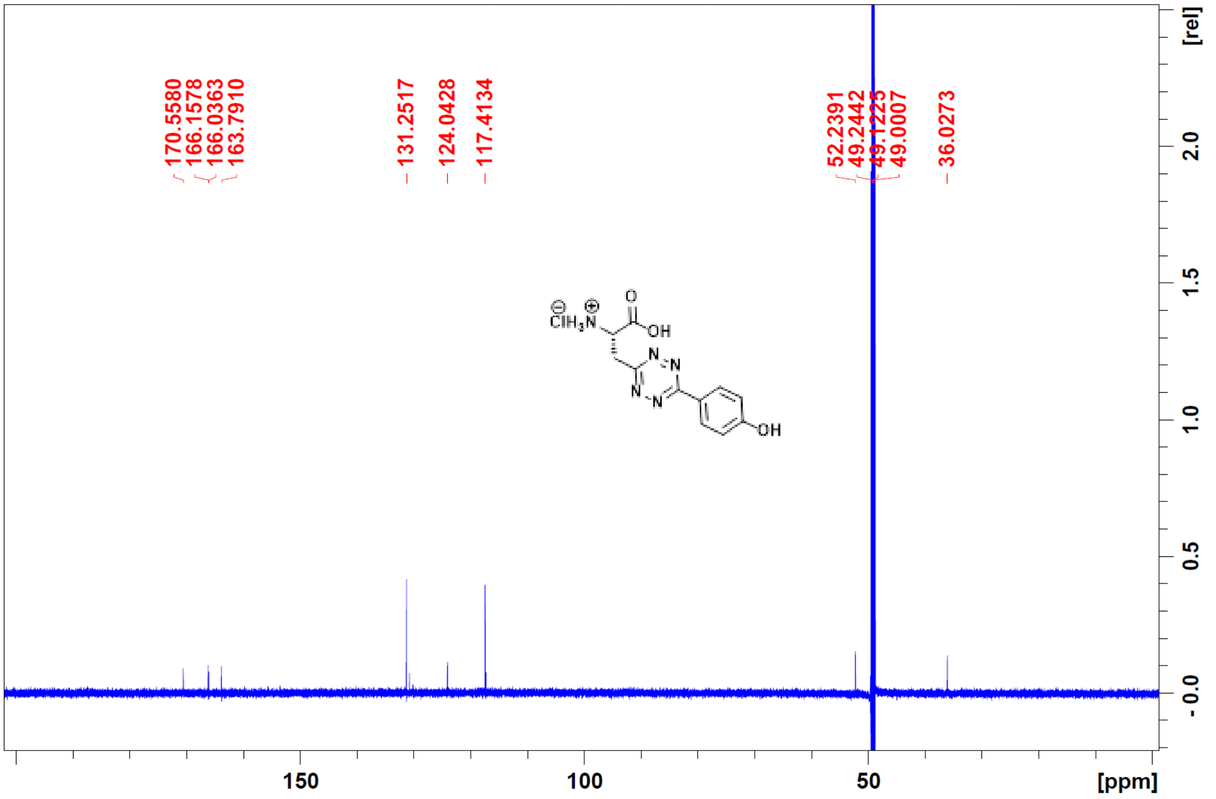

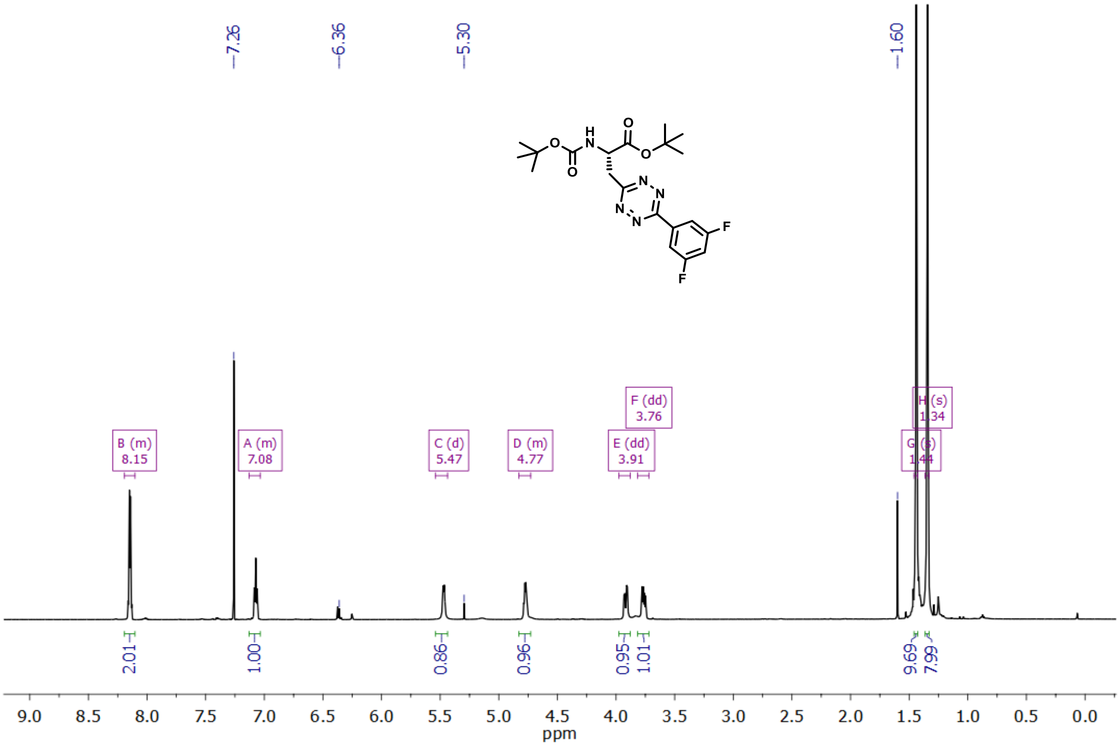
7.26
6.36
5.30
1.60
B (m) 8.15
A (m) 7.08
C (d) 5.47
D (m) 4.77
E (dd) 3.91
F (dd) 3.76
G (s) 1.44
H (s) 1.34
2.01
1.00
0.86
0.96
0.95
1.01
9.69
7.99
ppm

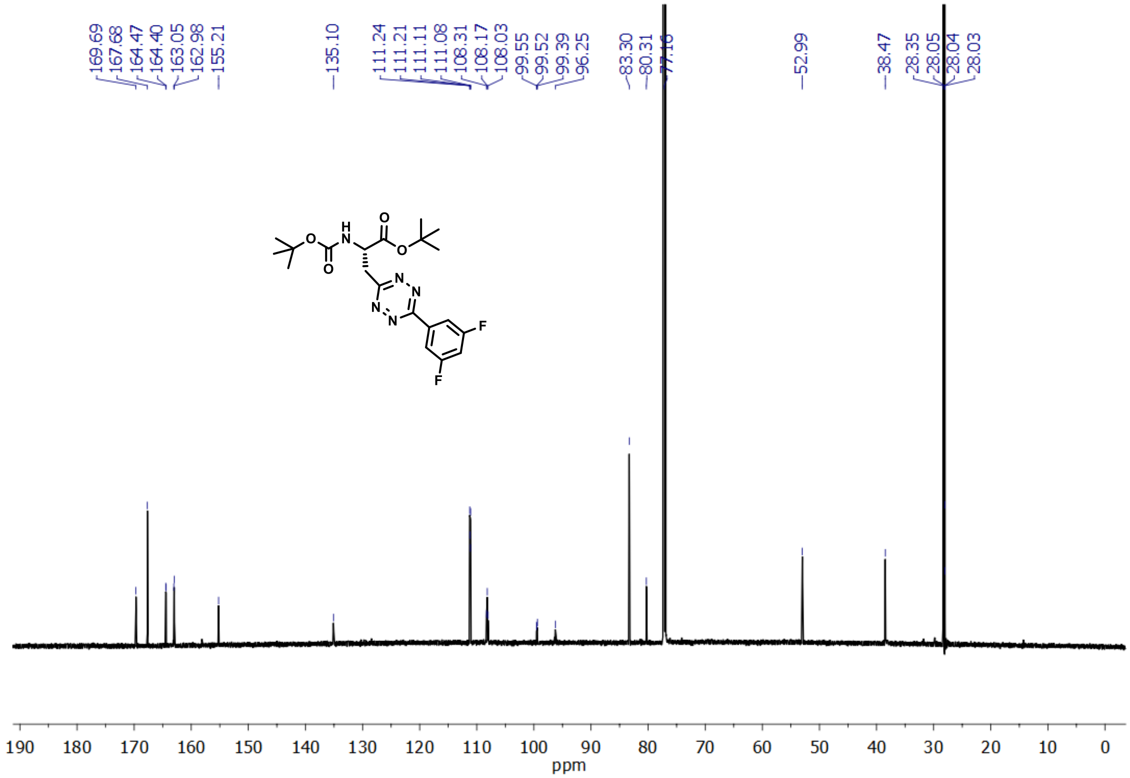
169.69
167.68
164.47
164.40
163.05
162.98
155.21
135.10
111.24
111.21
111.11
111.08
108.31
108.17
108.03
99.55
99.52
99.39
96.25
83.30
80.31
77.16
52.99
38.47
28.35
28.05
28.04
28.03
ppm

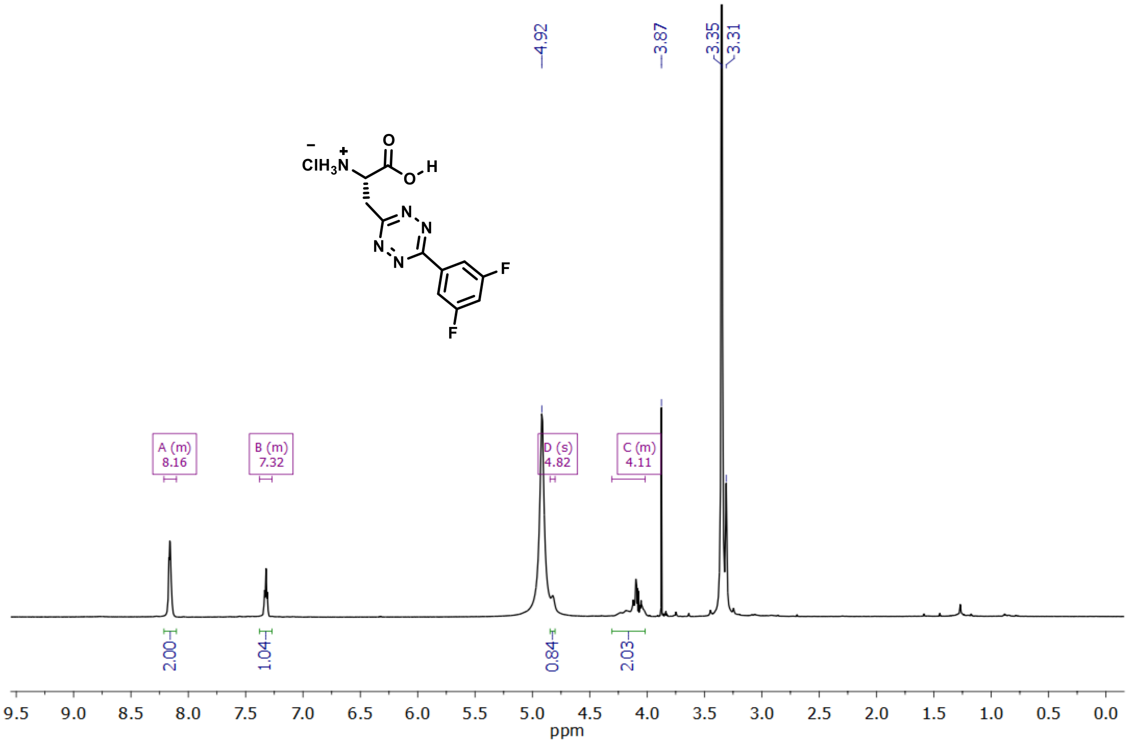
4.92
3.87
3.35
3.31
ClH3N
O
O
H
N
N
N
N
F
F
A (m)
8.16
B (m)
7.32
D (s)
4.82
C (m)
4.11
2.00
1.04
0.84
2.03
9.5 9.0 8.5 8.0 7.5 7.0 6.5 6.0 5.5 5.0 4.5 4.0 3.5 3.0 2.5 2.0 1.5 1.0 0.5 0.0
ppm

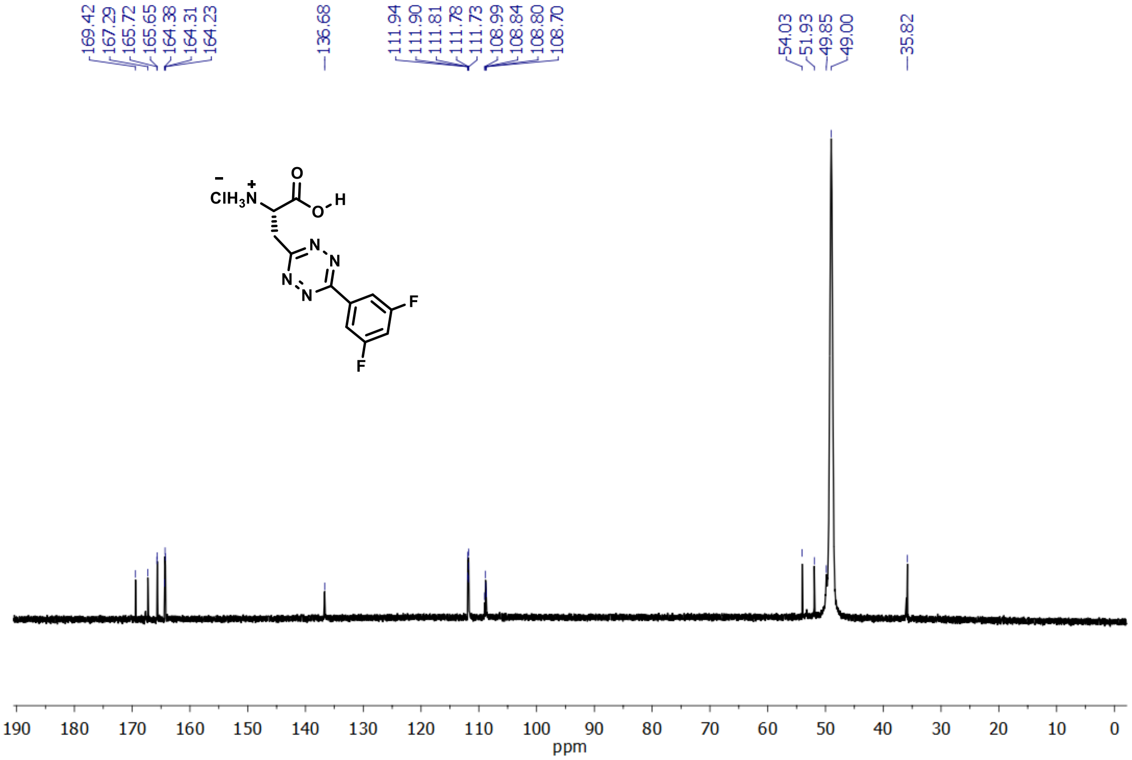
169.42
167.29
165.72
165.65
164.38
164.31
164.23
136.68
111.94
111.90
111.81
111.78
111.73
108.99
108.84
108.80
108.70
54.03
51.93
49.85
49.00
35.82
ClH3N
O
O
H
N
N
N
N
F
F
190 180 170 160 150 140 130 120 110 100 90 80 70 60 50 40 30 20 10 0
ppm

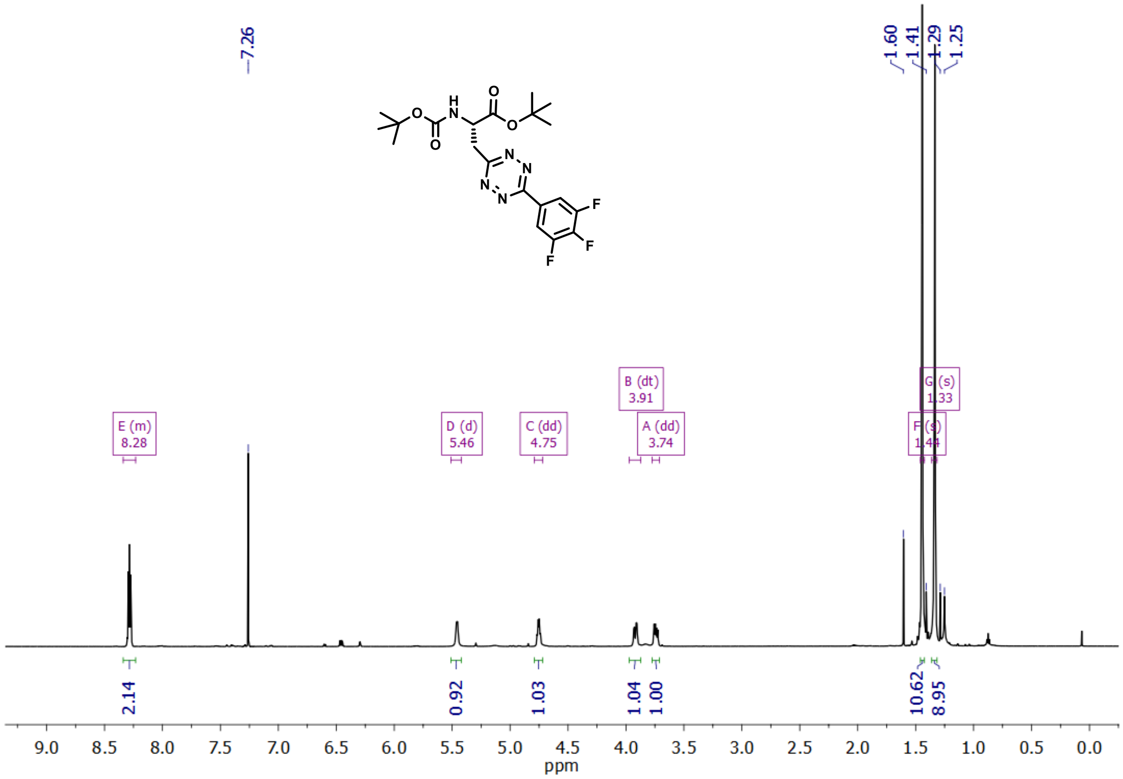

7.26
1.60
1.41
1.29
1.25
E (m) 8.28
D (d) 5.46
C (dd) 4.75
B (dt) 3.91
A (dd) 3.74
F (s) 1.44
G (s) 1.33
2.14
0.92
1.03
1.04
1.00
10.62
8.95
ppm

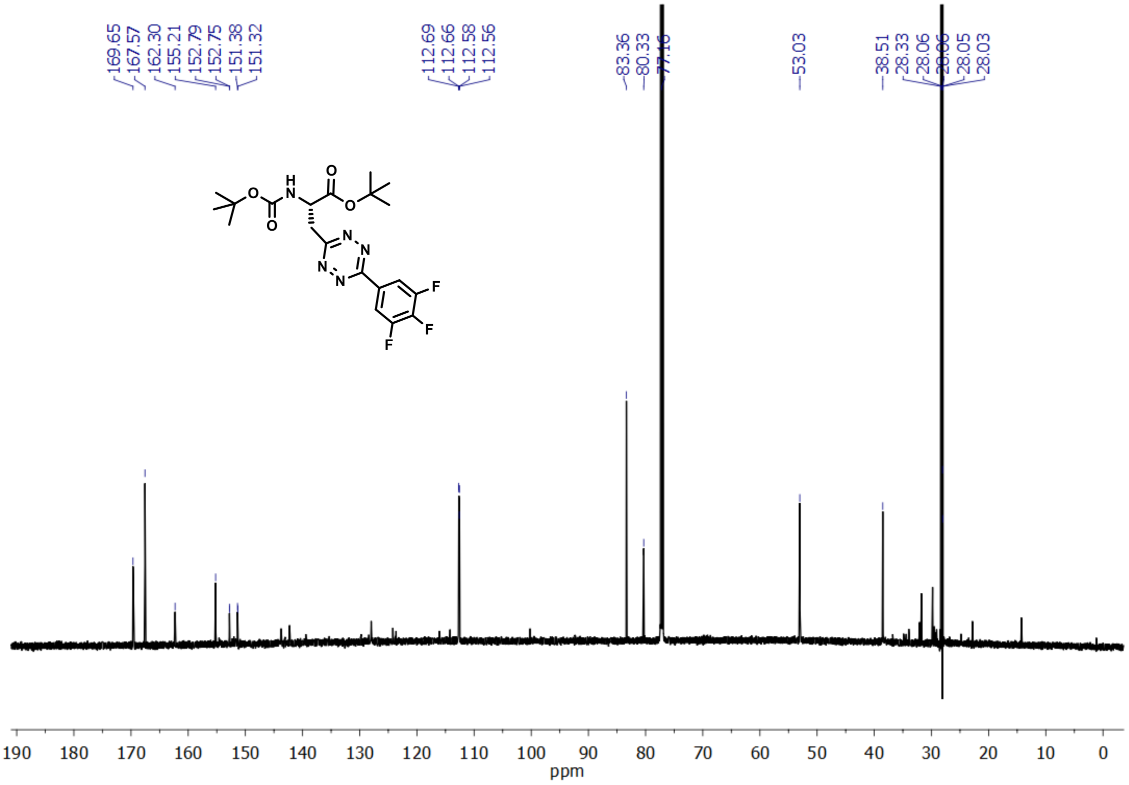

169.65
167.57
162.30
155.21
152.79
152.75
151.38
151.32
112.69
112.66
112.58
112.56
83.36
80.33
53.03
38.51
28.33
28.06
28.05
28.03
ppm

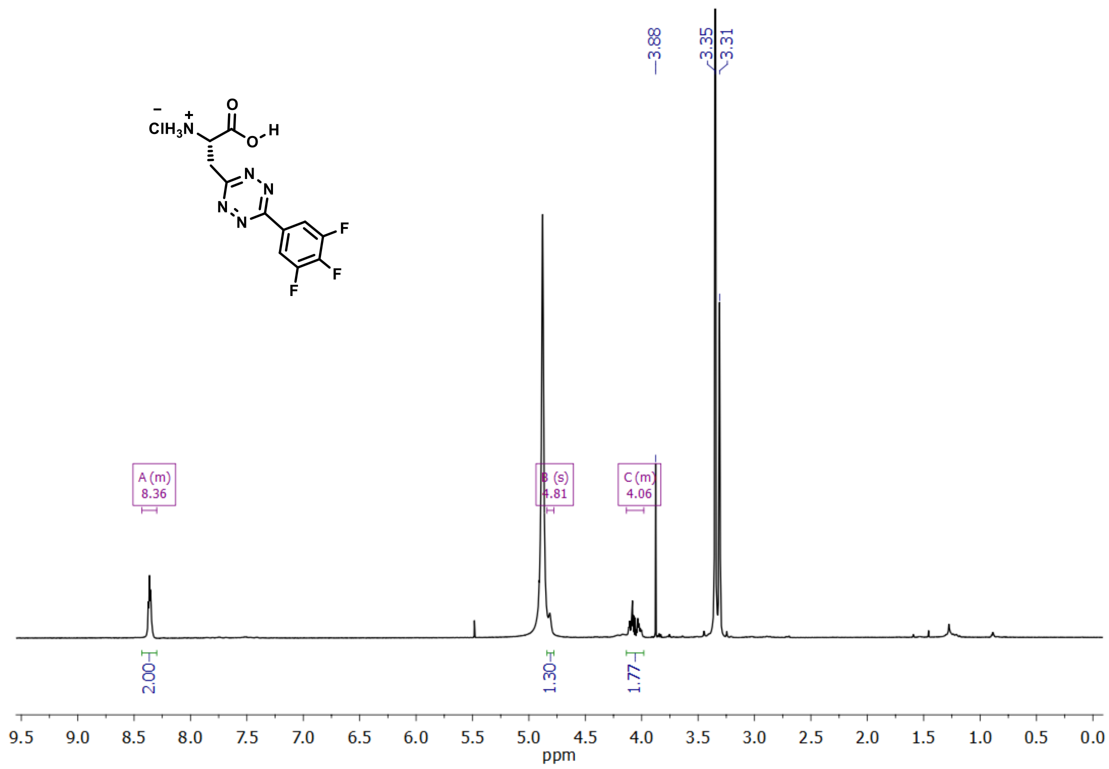

3.88
3.35
3.31
A (m)
8.36
B (s)
4.81
C (m)
4.06
2.00
1.30
1.77
9.5 9.0 8.5 8.0 7.5 7.0 6.5 6.0 5.5 5.0 4.5 4.0 3.5 3.0 2.5 2.0 1.5 1.0 0.5 0.0
ppm

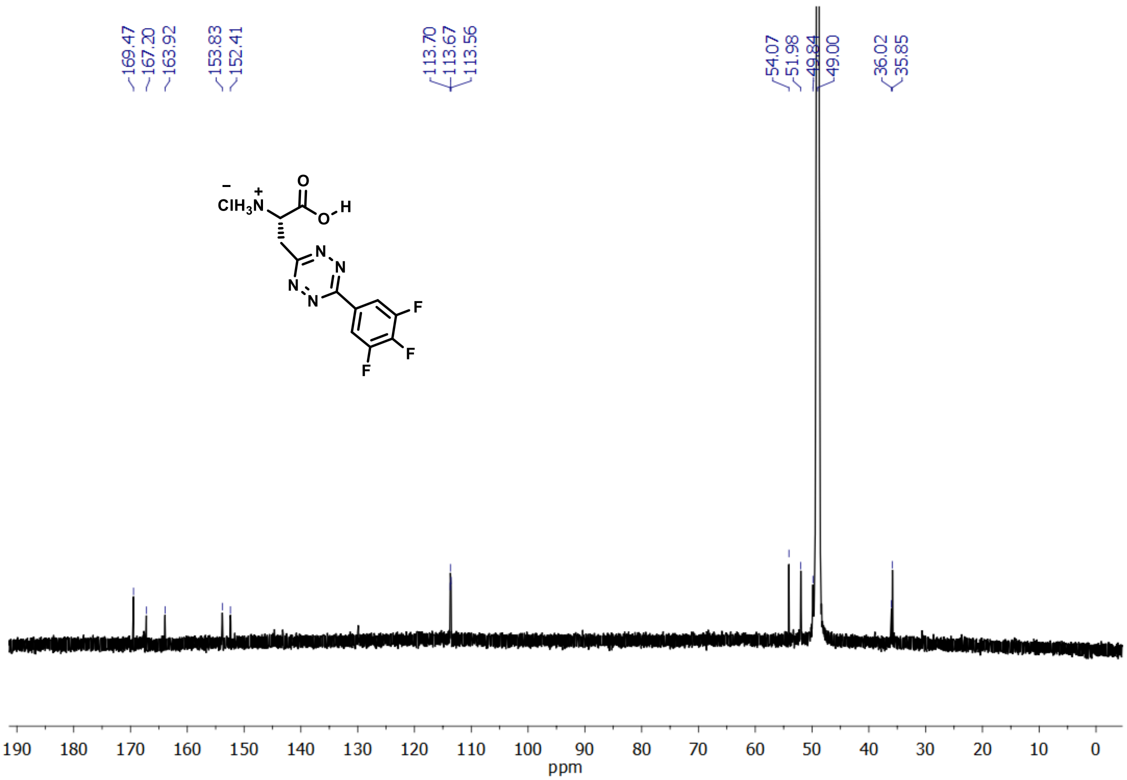

169.47
167.20
163.92
153.83
152.41
113.70
113.67
113.56
54.07
51.98
49.00
36.02
35.85
190 180 170 160 150 140 130 120 110 100 90 80 70 60 50 40 30 20 10 0
ppm
